# Average beta burst duration profiles provide a signature of dynamical changes between the ON and OFF medication states in Parkinson’s disease

**DOI:** 10.1101/2020.04.27.064246

**Authors:** Benoit Duchet, Filippo Ghezzi, Gihan Weerasinghe, Gerd Tinkhauser, Andrea A. Kühn, Peter Brown, Christian Bick, Rafal Bogacz

## Abstract

Parkinson’s disease motor symptoms are associated with an increase in subthalamic nucleus beta band oscillatory power. However, these oscillations are phasic, and there is a growing body of evidence suggesting that beta burst duration may be of critical importance to motor symptoms. This makes insights into the dynamics of beta bursting generation valuable, in particular to refine closed-loop deep brain stimulation in Parkinson’s disease. In this study, we ask the question “Can average burst duration reveal how dynamics change between the ON and OFF medication states?”. Our analysis of local field potentials from the subthalamic nucleus demonstrates using linear surrogates that the system generating beta oscillations is more likely to act in a non-linear regime OFF medication and that the change in a non-linearity measure is correlated with motor impairment. In addition, we pinpoint the simplest dynamical changes that could be responsible for changes in the temporal patterning of beta oscillations between medication states by fitting to data biologically inspired models, and simpler beta envelope models. Finally, we show that the non-linearity can be directly extracted from average burst duration profiles under the assumption of constant noise in envelope models. This reveals that average burst duration profiles provide a window into burst dynamics, which may underlie the success of burst duration as a biomarker. In summary, we demonstrate a relationship between average burst duration profiles, dynamics of the system generating beta oscillations, and motor impairment, which puts us in a better position to understand the pathology and improve therapies such as deep brain stimulation.

**Author summary:** In Parkinson’s disease, motor impairment is associated with abnormal oscillatory activity of neurons in deep motor regions of the brain. These oscillations come in the shape of bursts, and the duration of these bursts has recently been shown to be of importance to motor symptoms. To better understand the disease and refine therapies, we relate the duration of these bursts to properties of the system generating them in the pathological state and in a proxy of the healthy state. The data suggest that the system generating bursts involves more complexity in the pathological state, and we show that a measure of this complexity is associated with motor impairment. We propose biologically inspired models and simpler models that can generate the burst patterns observed in the pathological and healthy state. The models confirm what was observed in data, and tell us how burst generation mechanisms could differ in the disease. Finally, we identify a mathematical link allowing us to infer properties of the burst generating system from burst duration measurements in patient recordings. This sheds some light on the significance of burst duration as a marker of pathology.

## Introduction

The cardinal motor symptoms of Parkinson’s disease (PD) are slowness of movement (bradykinesia) or even inability to initiate movements, as well as rigidity due to increased muscle tone, and tremor. As suggested in [1], increased basal ganglia (BG) oscillatory activity in the beta band (13-35 Hz) has been correlated with worsening of motor symptoms, in particular bradykinesia and rigidity but not tremor [2–6]. It is believed that heightened synchrony in the beta band decreases the information coding capacity of the cortico-basal ganglia network [7], as recently confirmed [8]. PD is caused by a progressive loss of dopaminergic neurons, and can be successfully managed for a number of years by pharmacological treatment (the principal drug is Levodopa, a precursor of dopamine).

Physiological mesoscale beta activity in the cortex is of phasic nature and comes in bursts [9–11] (not to be confused with spike bursting). In PD, besides the average level of synchrony, the temporal patterning of beta activity has more recently been shown to be of importance. Specifically, the proportion of longer bursts of activity in the beta band of subthalamic nucleus (STN) local field potentials (LFPs) OFF medication has been correlated with motor impairment [12]. It was also found that motor symptoms can be ameliorated by shortening beta bursts of longer duration with STN adaptive deep brain stimulation (aDBS) [13]. Since then, STN bursts in PD patients have been shown to impact motor performance at the single trial level [14]. In another task, the percentage of time spent in beta bursts has been shown to be a better predictor of bradykinesia than average beta power [15], and it has been argued that temporal synchrony patterning may be more sensitive to clinical changes than average synchrony [16]. The clinical relevance of temporal synchrony patterning may extend to other PD motor symptoms, and STN beta burst duration has also been suggested as a potential biomarker for freezing of gait in PD [17]. Given this mounting body of evidence, providing insights into the dynamics of burst generation should put us in a better position to understand the pathology and treat it, in particular with targeted neurostimulation. In this study, we therefore ask the following question: can we relate observed changes in beta oscillation temporal patterning in PD between the ON and OFF medication states to changes in dynamical properties of the system generating beta oscillations?

In previous studies, STN beta bursts have been mostly studied based on one arbitrary threshold of the beta envelope as events above the threshold, potentially with a minimum duration condition (examples of various thresholds shown in Fig 1A). Average burst duration and amplitude profiles describing the average burst duration and amplitude for a range of thresholds (see Fig 1A and Fig 1B for an illustration of average burst duration profiles) have been introduced [12, 13]. However, they played a minor role in these studies and have not been considered systematically on an individual patient basis. Here, we leave behind the arbitrary choice of a threshold by relying on profiles across thresholds to provide an unbiased characterisation of beta oscillation temporal patterning. It has been established that STN beta burst duration is a better metric than burst amplitude to distinguish between the healthy and pathological states in an animal model of PD [18]. We begin our study by investigating in STN recordings of PD patients whether average burst duration profiles are better at distinguishing between the ON and OFF medication states than average burst amplitude profiles. We only observe significant changes in the temporal patterning of beta oscillations in average burst duration profiles, and come to the conclusion that burst duration is the better metric as in [18]. We introduce a burst duration specific measure of non-linearity based on linear surrogates, and show that our measure of non-linearity is increasing in recordings from the ON to the OFF state, thereby presenting a first level of description of the dynamical changes associated with changes in the temporal patterning of beta oscillations. To support the relevance of these changes, we study the correlation of our non-linearity measure with motor symptoms.

**Fig 1.**
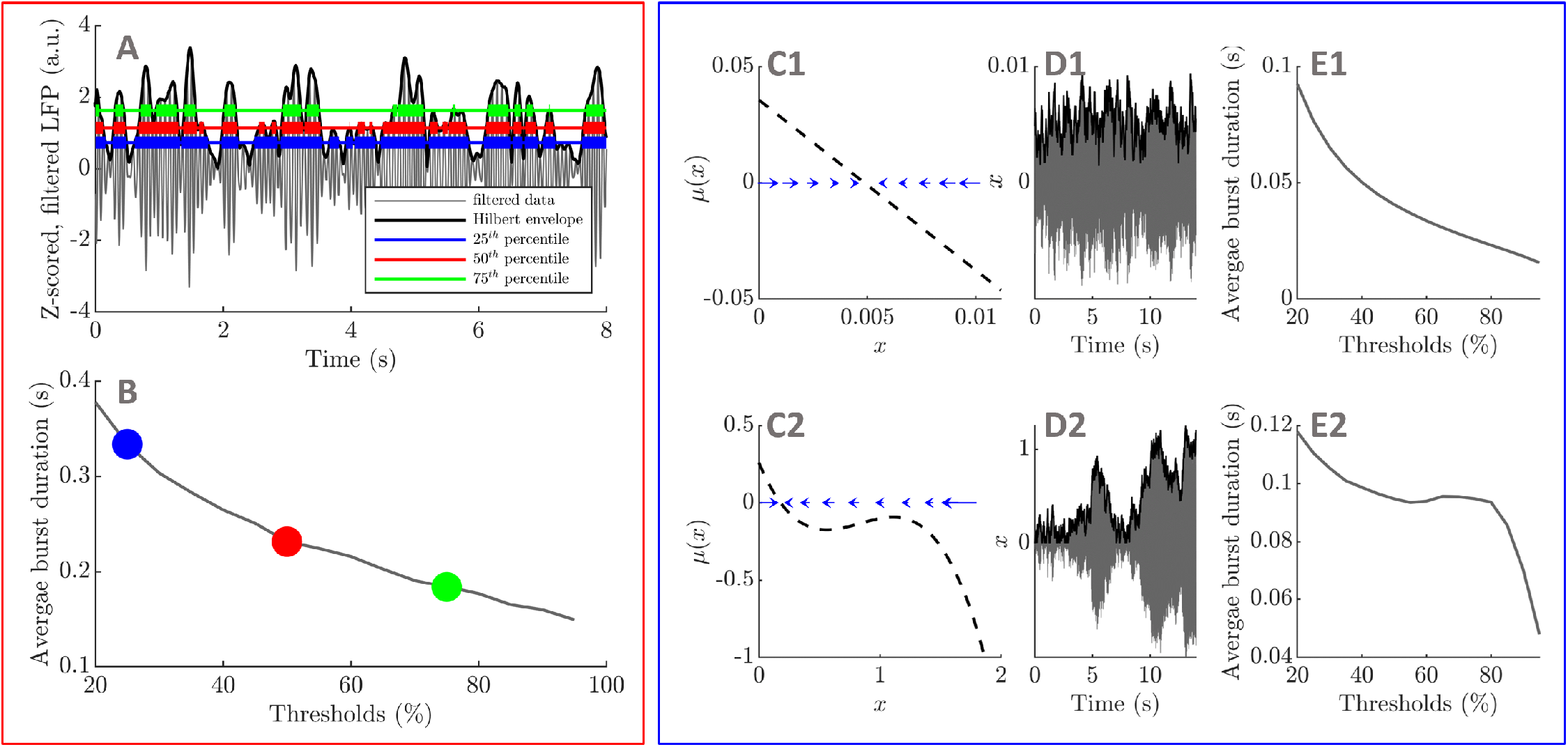
Introducing average burst duration profiles. Average burst duration profiles are obtained by computing beta envelope average burst duration for a range of thresholds. An example is provided for three thresholds, where thick lines highlight the duration of individual bursts for the three thresholds in panel A, and the corresponding averages are identified with the same colour in panel B. Considering the time discretization of simple envelope models of the form *dx*_*t*_ = *μ*(*x*_*t*_)*dt* + *ζdW*_*t*_, where *μ* is the drift function, *W* is a Wiener process, and *ζ* is a constant noise parameter, we illustrate with two example drift functions the link between envelope dynamics (panels C1 and C2, one-dimensional vector field also sketched with blue arrows) and average burst duration profiles (panels E1 and E2). The envelope models produce the black envelopes in panels D1 and D2, and beta oscillations (shown in grey in panels D1 and D2) can be obtained by adding a constant frequency phase equation. In C1, when *x* moves away from the fixed point, it will be strongly attracted back. By contrast, in the case of C2, if *x* is around 1, there is weak attraction towards the fixed point, allowing *x* to stay at an elevated level for longer.

Many modelling studies have reported the generation of sustained beta oscillations in the context of PD, and have identified several potential sources of exaggerated beta oscillations (see [19] for a review). However, reproducing the temporal patterning of beta oscillations has received little attention. It was reported very recently that spike bursting in a small proportion of neurons in the feedback loop formed by the STN and the globus pallidus pars externa (GPe) could contribute to transient beta oscillations [20]. Models of the STN-GPe feedback loop have been shown to generate realistic transient beta oscillations in response to beta frequency inputs from PD patient electroencephalogram (EEG) recordings [21], and in response to biologically inspired input patterns in healthy animals [22]. How model dynamics would need to change in the absence of correlated inputs to account for changes observed in the temporal patterning of beta oscillations in patients ON and OFF medication has not been investigated. To delineate the simplest two-population neural circuit dynamics that can reproduce average burst duration profiles ON and OFF medication, we fit time discretized Wilson-Cowan (WC) models [23] of increasing complexity receiving uncorrelated inputs to selected patient data.

The time discretization of one dimensional stochastic processes can be used to simulate the envelope of beta oscillations (see Fig 1C and Fig 1D). We call these models “envelope models”. Under simplifying assumptions, the envelope of a linear WC model was shown in [24] to be described by a particular envelope model. While the WC model is biologically inspired and describes the STN-GPe circuit, envelope models are simpler, can reproduce average burst duration profiles, and summarize the essence of the underlying dynamics. We use this to our advantage to pinpoint in all the datasets available the simplest polynomial forms of envelope drift function reproducing the ON and OFF medication states, and to derive an analytical expression for average burst duration profiles by identifying a first passage problem. Based on this result, we relate changes in average burst duration to changes in one specific parameter in the linear case.

Different envelope dynamics result in different average burst duration profiles (this is illustrated in Fig 1C, Fig 1D, and Fig 1E). To fully relate temporal patterning of beta oscillations to dynamics, we establish a mathematical link under general assumptions from average burst duration profile to envelope dynamics. This suggests that average burst duration profiles are a direct signature of envelope dynamics, and may be one reason why beta burst duration is found to be an important marker of pathology in experimental studies of PD. In addition, we illustrate the relationship between burst duration and dynamics by recovering envelope dynamics in envelope model synthetic data, and in examples of patient data. This envelope dynamics inference method may find applications in other contexts, as it can be applied to any envelope time series.

Starting with the data, the paper’s narrative will be guided by the following questions. Is the system generating beta oscillations more likely to operate in a non-linear regime OFF medication than ON medication (surrogate analysis, and envelope model subsections)? Can the change in a measure of non-linearity help predict motor impairment (surrogate analysis subsection)? Which are the simplest neural mass models that can reproduce the average burst duration profiles observed in ON and OFF medication data (neural mass model subsection)? Which type of non-linearity is required to explain experimental average burst duration profiles in the ON and OFF states (envelope model subsection)? How can we directly extract this non-linearity from average burst duration profiles (dynamics inference subsection)? The surrogate analysis is based on a statistical method which makes the least assumptions about the system, and is used to link dynamical changes with motor impairment. While the biologically inspired neural mass models describe the STN-GPe circuit, the simpler to fit envelope models are particularly insightful as they can be used to study the dependence of average burst duration on model parameters, and provide a direct link from average burst duration to drift function.

## Results

### Comparing bursting features ON and OFF medication

#### Choice of bursting features

Neural activity at beta frequency comes in the shape of bursts, and several features could be analysed to capture differences in beta bursting dynamics between medication states in STN LFPs. Differences in average beta power between medication states have been well documented in STN LFPs, see for instance [2, 25–28]. We therefore investigate bursting dynamics beyond simple differences in average beta power by individually z-scoring each dataset. To quantify the duration of bursts, we use as bursting feature the average burst duration profile. To quantify the amplitude of bursts, we use as bursting features the average burst amplitude profile, and the probability density function (PDF) of the envelope amplitude. To quantify the rate of occurrence of bursts, the average burst rate profile can be easily obtained from the average burst duration profile (see derivation in S1). The average burst rate profile therefore does not bring any additional information and is not included in the analysis.

Bursts are commonly defined as events corresponding to the envelope being above a predefined threshold for more than 100 ms [12, 13], but burst profiles across thresholds do not depend on an arbitrary threshold choice. Additionally, given a short time series, burst profiles make more efficient use of the data available than the distribution of burst durations at a single threshold. This is because they rely on a mean value for a given threshold and the same time series is reused for all thresholds. Although the duration of a given burst will necessarily decrease with increasing threshold, average burst duration profiles can be more complicated than a simple decreasing function of thresholds. This is illustrated in Fig 2, where the decrease in burst duration of individual bursts is more than compensated by the decrease in the proportion of shorter bursts when going from the 70^th^ percentile to the 80^th^ percentile. Importantly, dynamical properties of the system are revealed by average burst duration profiles as will be detailed later.

**Fig 2.**
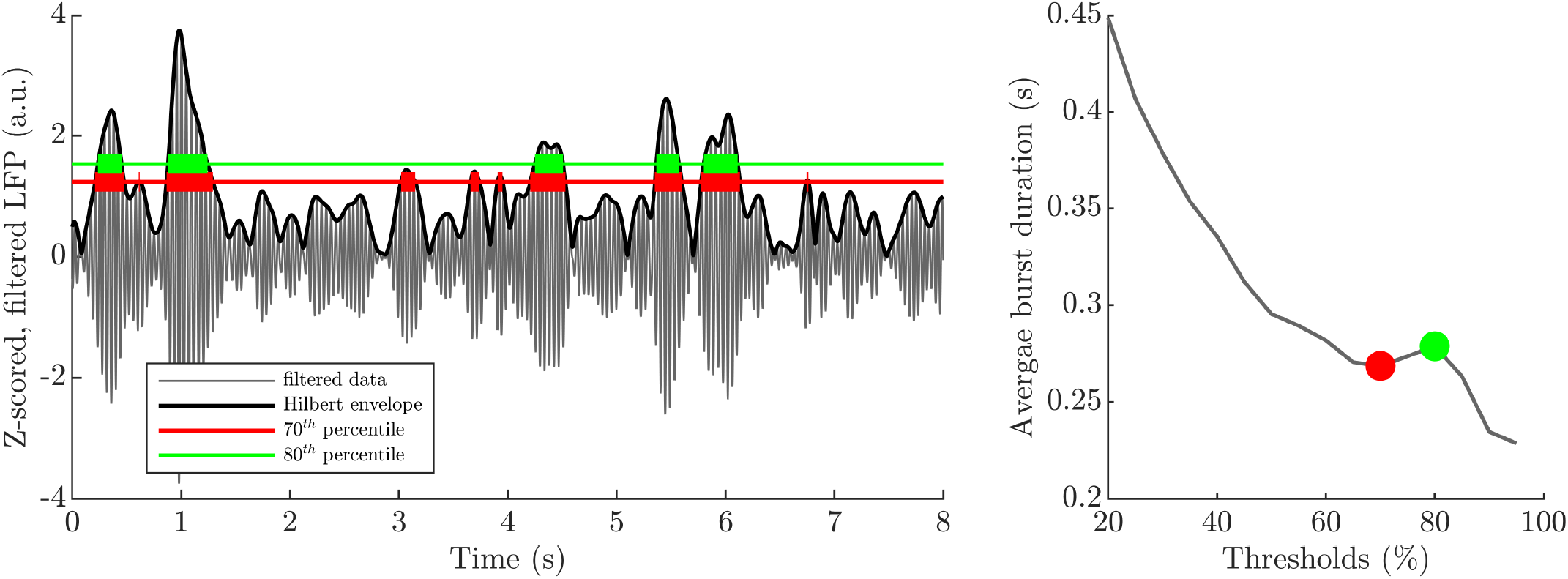
Average burst duration profiles can have complex shapes. Thick lines highlight the duration of individual bursts for two thresholds in the left panel. The corresponding averages are identified with the same colour in the average burst duration profile in the right panel. Longer bursts are still present at the 80^th^ percentile, and shorter bursts are significantly more frequent at the 70^th^ percentile than at the 80^th^ percentile. In addition, high amplitude, longer bursts are sharp (quick rise and fall), and therefore their duration is only shortened slightly from the 70^th^ to the 80^th^ percentile. As a result, the average burst duration profile is non-monotonic.

We extracted the power spectrum density (PSD) and the three bursting features mentioned above from filtered bilateral STN LFP recordings of 8 patients with advanced Parkinson’s disease ON and OFF Levodopa [2]. A detailed description of the data, as well as our data processing and feature extraction methodologies are provided in the subsection “Extracting power spectra and bursting features” in the Methods section.

#### The average burst duration profile is the relevant feature to discriminate bursting dynamics ON and OFF medication

Our statistical analysis reveals that the average burst duration profile is the most powerful of the three bursting features analysed (average burst duration profile, average burst amplitude profile, envelope amplitude PDF) to discriminate bursting dynamics between conditions. Significant differences in mean burst duration and mean burst amplitude between the ON and OFF states were assessed by t-tests (*n* = 5, *d.f.* = 8) with false discovery rate (FDR) control at 5% (more details on FDR control in the subsection “FDR control” in the Methods section). We evaluated significant differences in envelope PDFs between the ON and OFF conditions using cluster-based permutation testing (10^6^ permutations). The power spectra and the three bursting features are shown for the right hemispheres of the eight patients in Fig 3. Besides differences in the power spectra which are known to be statistically significant, only average burst duration profiles exhibit significant differences after FDR correction between the ON and OFF states. These significant differences are seen in most patients, although they are consistent across thresholds only for half of the datasets. Besides a difference in mean power removed by z-scoring, amplitude statistics were not found to be significantly different ON and OFF medication as exemplified by the lack of significant differences in both average burst amplitude profiles and envelope PDFs for all patients. A similar picture is seen in left hemisphere LFPs (see Supplementary Fig A in S2). We show in Supplementary Fig B and C in S2 that profiles obtained ON medication are different from similarly filtered pink noise (1/f noise) for the majority of hemispheres. This confirms that ON profiles are physiologically meaningful, despite the choice of filtering windows always centered on the beta peak OFF medication. We will therefore focus on average burst duration profiles in the rest of the paper, and we begin with a linear surrogate analysis of the changes in these profiles.

**Fig 3.**
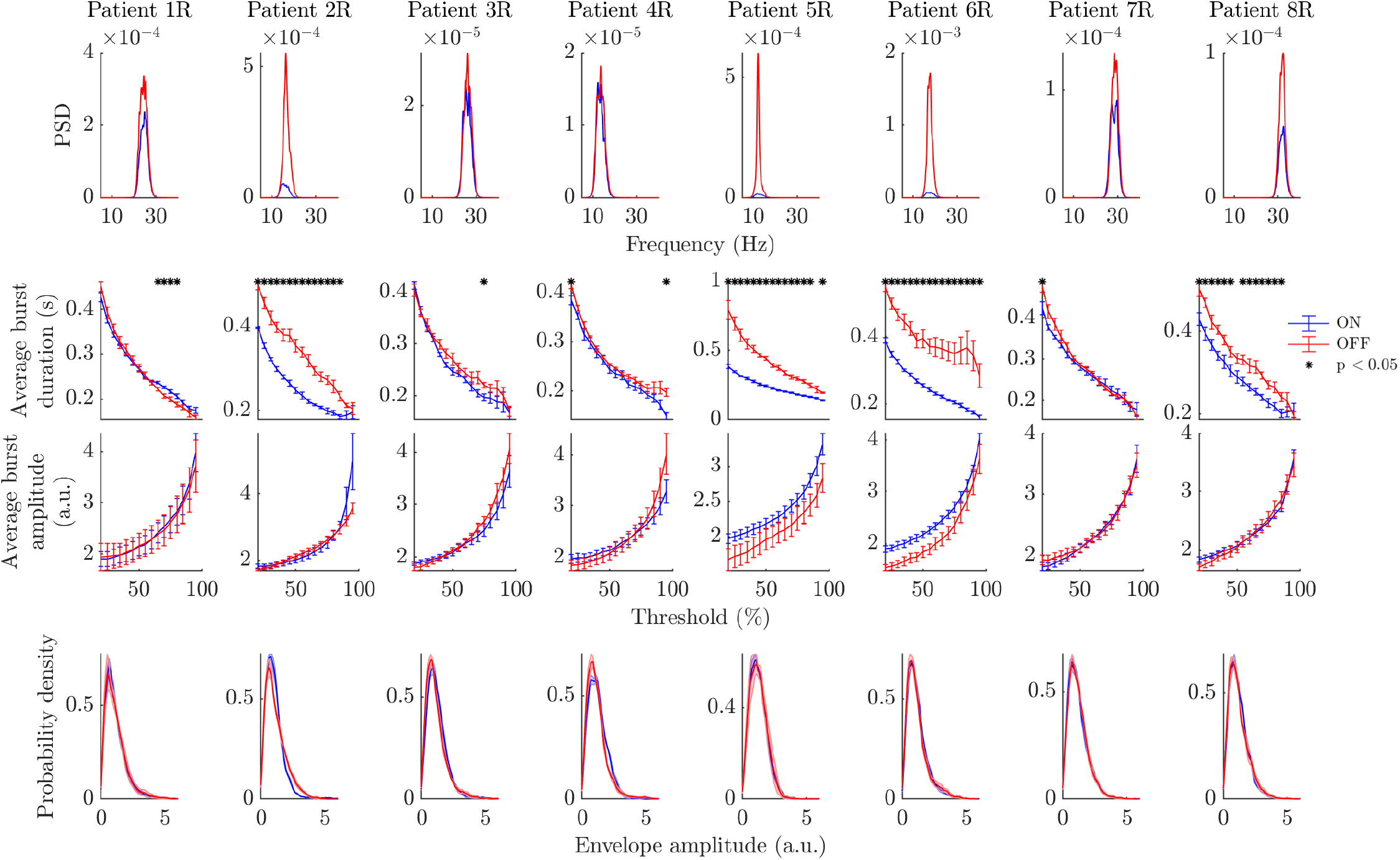
Power spectra and bursting features ON and OFF Levodopa (right hemispheres). Each column corresponds to the right hemisphere of one of the eight patients. Each row corresponds to a feature, the ON state is in blue, and the OFF state in red. The first row shows power spectra, the second row average burst duration profiles, the third row average burst amplitude profiles, and last row envelope amplitude PDFs. Statistically significant differences under FDR control are indicated by black stars (three bursting features only). Error bars represent the SEM.

### Investigating changes in dynamics using a linear surrogate analysis of average burst duration profiles

Our first approach to relating changes in the temporal patterning of beta oscillations between the ON and OFF states to dynamical changes is to consider the degree of non-linearity in the ON and OFF states. It is obtained by comparing average burst duration profiles ON and OFF medication to the profiles of their respective linear surrogates. In addition, we provide support for the relevance of these changes by reporting correlations with motor impairment.

#### Linear surrogates

Linear surrogates provide a way of testing for the presence of non-linearity in the system that generated an observed time-series [29]. Here linear system refers to a stationary linear stochastic process, an example of which is a stationary linear Gaussian process (in discrete time an auto-regressive moving average model or ARMA model). An ARMA(p,q) model can be described as

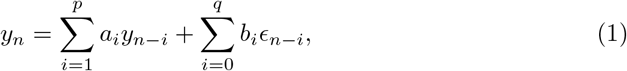

where *ϵ*_*i*_ are independent Gaussian white noise increments, and *a*_*i*_ and *b*_*i*_ are constant coefficients. The value at time *n* is a linear combination of past values and noise terms.

Linear surrogates preserve the linear properties of the data (linear correlations) but erase any potential non-linear structure. Besides the mean and standard deviation, linear properties are limited to the auto-correlation at all lags in the time domain, which is equivalent to the power spectrum in the frequency domain via the Wiener-Khinchin theorem. Thus, linear surrogates preserve the PSD.

The most common linear surrogates, Fourier transform (FT) surrogates and iterated amplitude adjusted Fourier transform (IAAFT) surrogates assume that the data is stationary (more details on these methods are given in “From FT surrogates to GWR surrogates” in the Methods Section). Depending on their duration, nonstationarity may be present in LFP recordings, and could be mistaken for non-linearity by FT and IAFFT surrogates. In this work, even if the recordings used are short (on the order of 250 s), we rely on gradual wavelet reconstruction (GWR) surrogates, which can mitigate the nonstationarity issue. GWR surrogates are provided along a continuum parametrised by *ρ*, where *ρ* = 0 corresponds to IAAFT surrogates, and *ρ* = 1 corresponds to the data [30]. In addition to addressing the influence of nonstationarity, this continuum allows to quantify effect strength. More details on the GWR method can be found in “From FT surrogates to GWR surrogates” (Methods section). Nineteen GWR surrogates were computed from the filtered data for each patient and hemisphere, ON and OFF medication, and for each *ρ* level ranging from 0 to 0.9 in steps of 0.1 with an additional value at 0.99 (as close to the data as possible). At *ρ* = 0.1, the largest wavelet coefficients making up 10% of the total wavelet energy are left out of the randomization procedure, which ensures that most of the data temporal variability is included in surrogates (see surrogates shown in the Methods Section for a range of *ρ* levels).

#### Changes in a burst duration specific measure of non-linearity between the ON and OFF states

To evaluate dynamical changes between the ON and OFF states, we define a burst duration specific measure of non-linearity based on linear surrogates, and show that it is significantly greater OFF than ON medication.

We obtain average burst duration profiles and PSDs from the filtered data and surrogates as detailed in “Extracting power spectra and bursting features” (Methods Section), except that no segmentation is done and the surrogates are not filtered (as they already reproduce the spectrum of the filtered data). Average burst duration profiles of linear surrogates and filtered data are shown for the right hemispheres of the eight patients in Fig 4 for *ρ* = 0. Similar figures are provided for a range of *ρ* levels for both hemispheres (Supplementary Fig D and E in S2), and show that surrogate PSDs very closely match data PSDs as expected.

**Fig 4.**
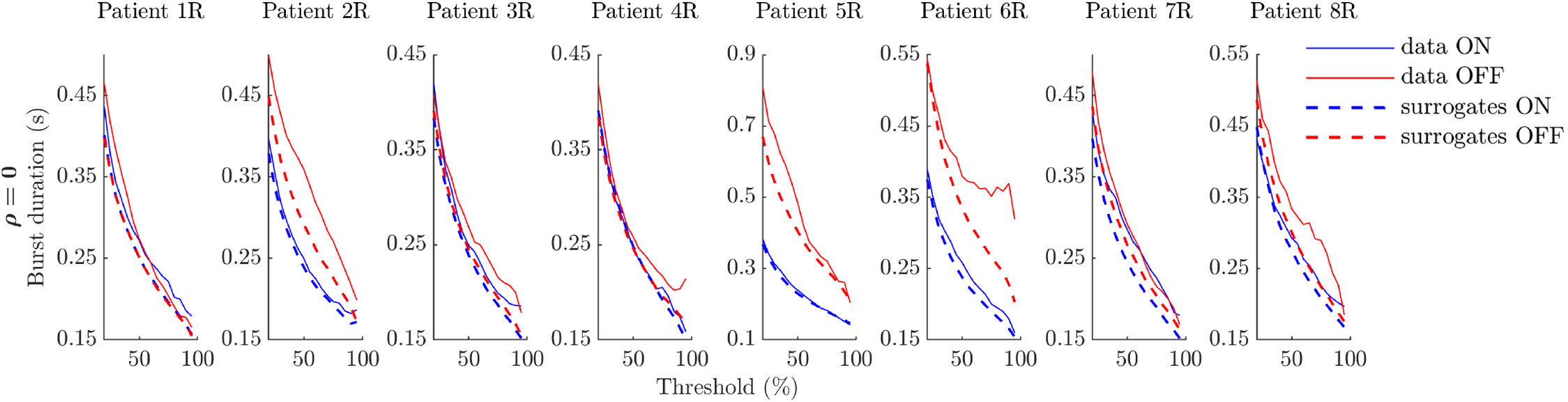
Average burst duration profiles ON and OFF medication for data and GWR surrogates at *ρ* = 0 (right hemispheres). In all panels, data profiles are solid lines, while linear surrogate profiles are dashed lines. The OFF medication state is indicated in red, and the ON state in blue.

As shown in Fig 5A, we define a non-linearity measure specific to burst duration profiles as the sum of the squared differences between filtered data and linear surrogate average burst duration profiles, relative to the square of the mean value of the surrogate average burst duration profile. We refer to this measure as BDDLS, which stands for burst duration distance to linear surrogate. We call the difference in BDDLS OFF and ON medication BDDLSdiff.

**Fig 5.**
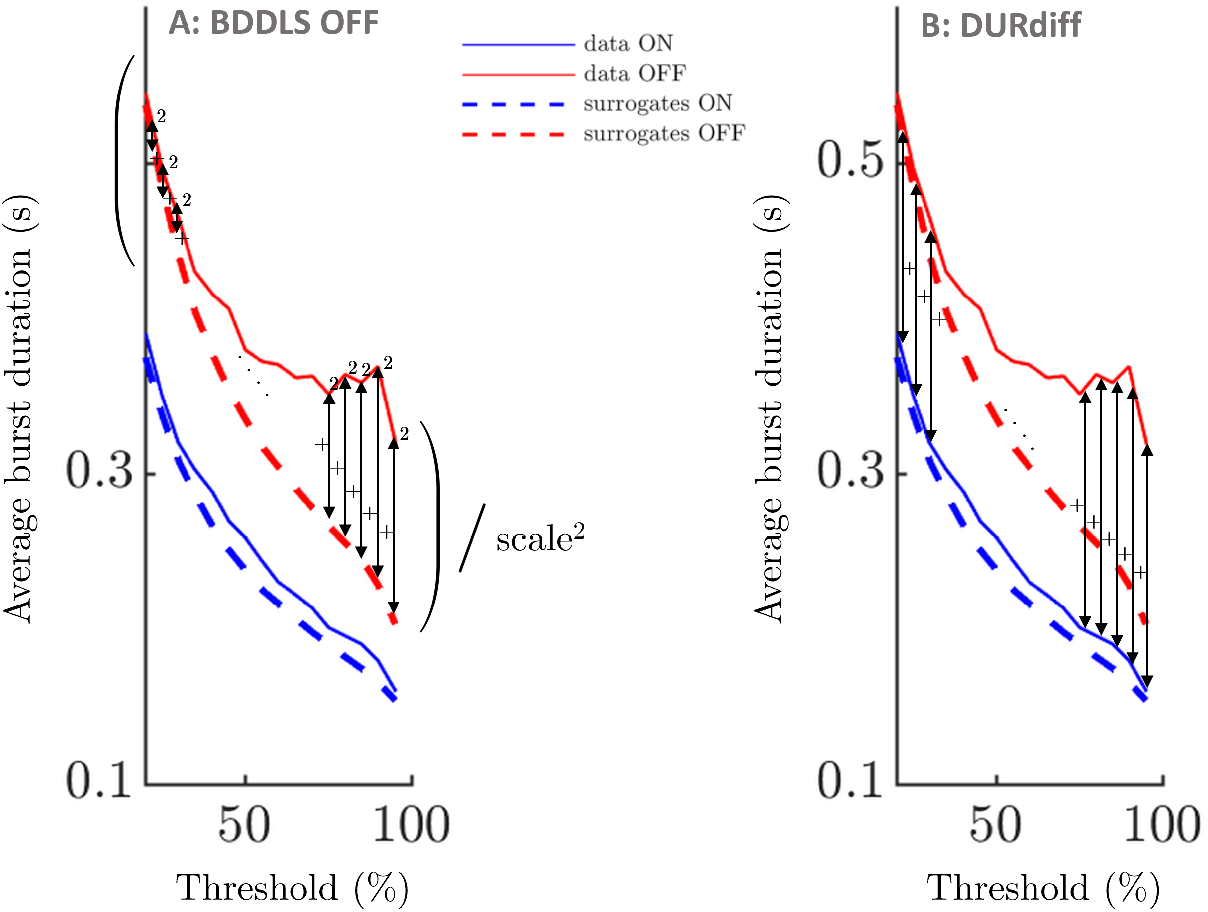
Sketch of burst duration metrics. A: illustration of burst duration distance to linear surrogates in the OFF state (BDDLS OFF). BDDLS OFF is defined as the sum of squared differences between data and linear surrogate average burst duration profiles for the OFF condition divided by the square of a scale. The scale is taken as the mean value of the OFF linear surrogate average burst duration profile. BDDLS ON is defined in a similar way, and BDDLSdiff is BDDLS OFF medication minus BDDLS ON medication. B: DURdiff is defined as the sum of the differences across thresholds between burst duration profiles OFF and ON medication. In this figure, summation is indicated by the symbol +, division by the symbol */*, and squaring by the symbol ^2^.

Our burst duration specific measure of non-linearity, BDDLS, was found significantly greater OFF than ON medication from *ρ* = 0 up to *ρ* = 0.6 under FDR control as shown in Table 1 (one-tailed Wilcoxon signed rank test, all patients and hemispheres, *n* = 16 per condition). Scatter plots corresponding to all of the analysed *ρ* levels are provided in Supplementary Fig F in S2. Since the BDDLS measures include a scaling by the square of the mean value of the surrogate average burst duration profile, the effect of medication state cannot be due to the profiles of data and surrogates having overall larger values OFF medication. Two conclusions can be drawn from the effect being significant up to *ρ* = 0.6. Firstly, the effect is not due to nonstationarity in the data (as mentioned earlier, GWR surrogates at *ρ* = 0.1 already look very similar to the data, and thus capture the major non-stationary features that may be present). Secondly, significance up to *ρ* = 0.6 implies that a limited amount of phase randomization in the surrogates is enough to start seeing a significant difference between BDDLS OFF and ON medication, hinting at a strong effect. More details on *ρ* can be found in “From FT surrogates to GWR surrogates” (Methods section).

**Table 1.** Statistical significance of medication state effect on BDDLS. Showing p-values for the test that BDDLS is greater OFF than ON medication (sign rank test, all patients, both hemispheres, *n* = 16 per condition) as a function of the GWR surrogate parameter *ρ*. P-values in bold are smaller than 5%, while green indicates significance under FDR control.

| $\rho$ | 0 | 0.1 | 0.2 | 0.3 | 0.4 | 0.5 | 0.6 | 0.7 | 0.8 | 0.99 |
| --- | --- | --- | --- | --- | --- | --- | --- | --- | --- | --- |
| p-val | **0.0010** | **0.0009** | **0.0010** | **0.0009** | **0.0003** | **0.0081** | **0.0012** | 0.259 | 0.510 | 0.330 |

As a control, we calculated BDDLS OFF and ON medication for data band-pass filtered ± 3 Hz around 35 Hz, for *ρ* = 0. The control condition aims to show some frequency specificity of the difference in BDDLS between conditions. The choice of *ρ* = 0 is therefore conservative (a difference is more likely when the surrogates are the most different from the data). No effect of medication state on BDDLS was found (p = 0.449, *n* = 16 per condition, and see Supplementary Fig G and Fig H in S2). Although band-pass filtered data around 35 Hz may be filtered noise (at least partially), we observe that differences in power do not necessarily translate into differences in BDDLS, as PSDs vary considerably between the OFF and ON states in the controls, but BDDLS OFF and ON medication are similar. As BDDLS is greater in the OFF state, we investigate below whether BDDLSdiff correlates with motor impairment.

#### Clinical correlations

To show that the difference in BDDLS OFF and ON medication, BDDLSdiff, is indicative of motor impairment, we also consider two other metrics as possible predictors of motor impairment. The first one is the relative difference between PSD OFF and ON medication called PSDdiff (the difference has to be relative to allow an analysis across patients as PSD levels vary greatly), and the second one is the sum of the differences between burst duration profiles OFF and ON medication across thresholds called DURdiff (similar scale across patients). DURdiff is illustrated in Fig 5B. Motor impairment was measured using the unified Parkinson’s disease rating scale (UPDRS) OFF medication. To increase test sensitivity, the rating was done using half points as opposed to integers only [2]. The correlations we report are for predictors and hemibody UPDRS OFF medication averaged across sides. The Spearman’s correlations described below are therefore for *n* = 8, *d.f.* = 6.

BDDLSdiff correlates with UPDRS OFF medication. The correlation is statistically significant under FDR control from *ρ* = 0.1 to *ρ* = 0.5 (Spearman’s correlation coefficients and associated p-values are shown in Table 2). This suggests a robust effect as explained previously. The other factors of interest PSDdiff (relative change in PSD) and DURdiff (change in burst duration) are also correlated with UPDRS OFF medication (Spearman’s correlation of 0.500, p = 0.216, and of 0.476, p = 0.243, respectively). The lack of statistical significance for these factors is likely due to the small sample size (eight subjects). The relationship between predictors and changes in clinical scores is plotted in Supplementary Fig I in S2.

**Table 2.** Spearman’s correlations between BDDLSdiff and UPDRS score OFF medication. Values are presented as a function of the GWR surrogate level *ρ*. P-values in bold are smaller than 5%, while green indicates significance under FDR control. Predictors and hemibody UPDRS OFF are averaged across sides (*n* = 8, *d.f.* = 6).

| $\rho$ | 0 | 0.1 | 0.2 | 0.3 | 0.4 | 0.5 | 0.6 | 0.7 | 0.8 | 0.99 |
| --- | --- | --- | --- | --- | --- | --- | --- | --- | --- | --- |
| correlation | 0.571 | 0.738 | 0.810 | 0.929 | 0.833 | 0.810 | 0.333 | -0.048 | -0.571 | 0.595 |
| p-val | 0.1511 | **0.0485** | **0.0218** | **0.0022** | **0.0154** | **0.0218** | 0.428 | 0.935 | 0.151 | 0.132 |

The three predictors considered are correlated, although with one exception not significantly so. Spearman’s correlations are 0.500, p = 0.216 between BDDLSdiff for *ρ* = 0.2 and PSDdiff, 0.476, p = 0.243 between BDDLSdiff for *ρ* = 0.2 and DURdiff, and 0.952, p = 0.001* between DURdiff and PSDdiff, where * denotes significance under FDR correction. The lack of significance in the remaining comparisons may be due to the relatively low number of subjects. The relationship between the predictors is plotted in Supplementary Fig J in S2. However, scaling BDDLS by the mean value of the surrogate average burst duration profile decorrelates as much as possible BDDLSdiff from DURdiff. Moreover, non-linear correlations cannot be recovered from the PSD (the power spectrum only captures linear correlations) which implies that BDDLSdiff should contain information not present in PSDdiff if the system is non-linear and non-linear behaviour is reflected in its average burst duration profile.

We have made apparent that changes in the temporal patterning of beta activity between medication states can be related to an increase in our burst duration specific non-linearity metric OFF medication, and that changes in the BDDLS metric are correlated with motor symptoms. Average burst duration profiles OFF medication rank higher on the BDDLS metric and can therefore be thought of as being generated by a system more likely to operate in a non-linear regime than ON medication. By non-linear regime we mean a regime where the generated signal covers enough of the dynamical system vector field that the system non-linearity translates into non-linear structure in the generated signal. To get a clearer idea of which specific dynamical changes could be at play between the ON and OFF states, we proceed to model chosen datasets ON and OFF medication using neural mass models, and identify the simplest dynamical systems that can fit the data in each state.

### Investigating changes in dynamics using neural mass models

To investigate which particular changes to the dynamics of a biologically inspired model of the STN-GPe loop could account for the changes in the temporal patterning of beta oscillations observed between the OFF and ON condition, we fit WC models of increasing complexity to patient data. Because fitting neural mass models to patient data is computationally expensive, we select the top two datasets with the largest BDDLS for *ρ* = 0 (these are OFF medication datasets), and the corresponding ON datasets. These datasets are patient 6, right hemisphere (which we denote patient 6R), and patient 4, left hemisphere (which we denote patient 4L). Datasets 6R OFF and 4L OFF display the most striking average burst duration profile deviations from their respective linear surrogates (see Fig 4 and Supplementary Fig D in S2). Datasets 6R ON and 4L ON have average burst duration profiles typical of the ON state.

#### Fitting a linear Wilson-Cowan model to ON state datasets

We start by fitting to patient 6R ON and 4L ON the time discretization of a linear WC model which describes the interactions between an excitatory and an inhibitory population of neurons. The discrete linear WC model can be seen as a multivariate version of an ARMA model (equation (1)). The WC model is a natural choice as it can be mapped onto the basal ganglia STN-GPe loop as shown in Fig 6, the STN being modelled as the excitatory population, and the GPe as the inhibitory population [31–33]. While the STN-GPe circuit is characterised by complex currents at the microscopic level, our choice to work with the mesoscopic WC model is based on the mesoscopic character of the data available. As a heuristically derived mean-field model, the WC model benefits from a low number of parameters while retaining some level of description of a microscopic biological reality [23]. The LFP recordings used in this study have been obtained from DBS electrodes implanted in the STN, and therefore we model the LFP signal by the activity of the excitatory population. The STN and GPe are reciprocally connected, and the STN receives a constant excitatory input from the cortex, while the GPe receives a constant inhibitory input from the striatum, and is also self-inhibiting. Uncorrelated inputs, specifically Gaussian white noise, are added to each population. The STN activity, *E*, and the GPe activity, *I*, are described by the stochastic differential equations

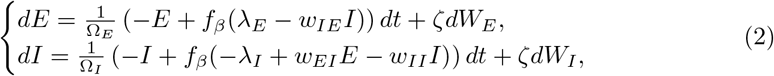

with *w*_*PR*_ the weight of the projection from population “P” to population “R”, *λ*_*P*_ the constant input to population “P”, and Ω_*P*_ the time constant of population “P”. In addition, *W*_*E*_ and *W*_*I*_ are Wiener processes, and *ζ* is the noise standard deviation. In this attempt to describe simple ON state average burst duration profiles, we use a linear activation function simply given by *f*_*β*_ (*x*) = *βx*, where *β* is the slope.

**Fig 6.**
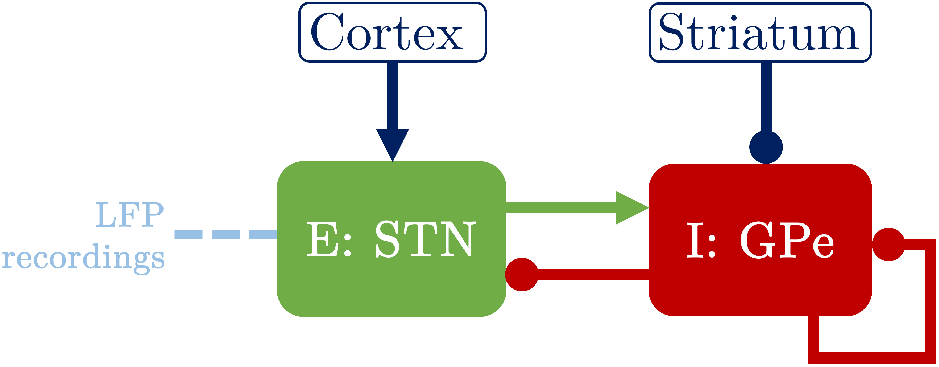
Mapping of the Wilson-Cowan model onto the STN-GPe loop. The excitatory population E and the inhibitory population I model the basal ganglia STN and GPe, respectively. Arrows denote excitatory connections or inputs, whereas circles denote inhibitory connections or inputs. The DBS electrode is implanted in the STN (indicated by a dashed light blue line) and records the STN LFP. The STN also receives an excitatory input from the cortex, while the GPe receives an inhibitory input from the striatum, and also has a self-inhibitory loop.

The time discretized model is fitted to two features (also known as summary statistics) of the data, namely the data PSD and the data average burst duration profile. How fitting is carried out is described in “Fitting procedure” in the Methods section. As the model output *E* models the centered LFP recordings, a model of the beta envelope is obtained by considering the Hilbert amplitude of *E* (modulus of the analytic signal of *E*).

The best fits to patients 6R ON and 4L ON are shown in the first and third rows of Fig 7, and we report the corresponding model parameters in Supplementary Table A in S3. The linear WC model is able to reproduce both the data PSD and average burst duration profile in both cases (coefficient of determination *R*^2^ = 0.756 for 6R ON, see Fig 7C1 and Fig 7D1, and *R*^2^ = 0.798 for 4L ON, see Fig 7C3 and Fig 7D3). The input parameters *λ*_*E*_ and *λ*_*I*_ only contribute to transients in the linear WC model, and are therefore set to zero here. They will have an influence on the model output when non-linearity is introduced.

**Fig 7.**
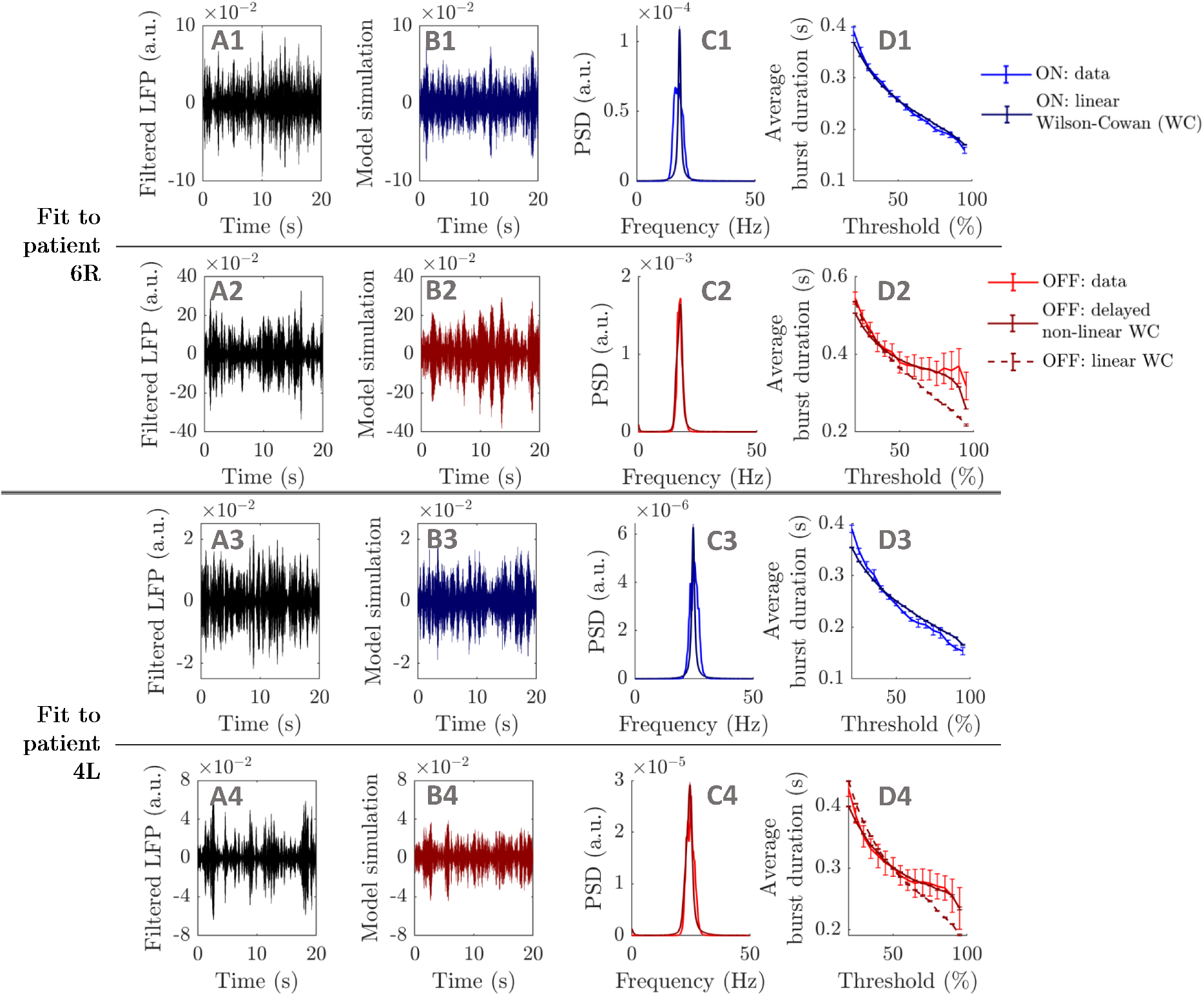
Best WC model fits. Showing best fits to datasets 6R (ON: first row, OFF: second row), and 4L (ON: third row, OFF: fourth row). The first column shows twenty seconds of filtered LFP recording (A panels), while the same duration of model oscillatory activity output is plotted in the second column (B panels). Data and model PSDs are compared in the third column (C panels), and data and model average burst duration profiles are shown in the last column (D panels, SEM error bars). In the first and third rows, all model outputs correspond to fits of the linear WC model. In the second and fourth rows, dark red solid lines correspond to fits of the delayed non-linear WC model, and dark red dashed lines to fits of the linear WC model (shown for comparison, D panels only).

We have shown that a linear WC model can fit to patients 6R ON and 4L ON, and we next fit increasingly complex WC models to patients 6R OFF and 4L OFF to investigate the OFF medication case.

#### Fitting Wilson-Cowan models to the highest BDDLS patients

The same procedure as before is used to fit the time discretized linear WC model to patients 6R OFF and 4L OFF. The linear WC model is able to reproduce the data PSD (not shown) but does not fit well to the data average burst duration profile in both cases (see dashed lines in Fig 7D2 and Fig 7D4, *R*^2^ = 0.312 for patient 6R OFF and *R*^2^ = 0.635 for patient 4L OFF). We report the corresponding model parameters in Supplementary Table A in S3.

We next introduce the time discretization of a non-linear WC model, which is identical to the linear WC model (see Fig 6), except that its activation function

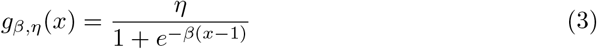

is non-linear and that connections carry a delay. The system of equations (2) is therefore modified as

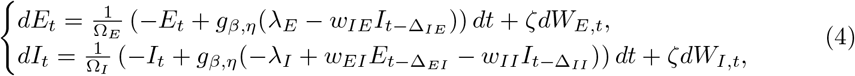

where Δ_*PR*_ is the delay from population “P” to population “R”, and *η* is a scaling parameter.

The best fits to patients 6R OFF and 4L OFF obtained with the same fitting methodology as before are shown in the second and fourth rows of Fig 7 (corresponding model parameters in Supplementary Table B in S3). The non-linear WC model with delays is able to reproduce the data average burst duration profile and the data PSD well in both cases (*R*^2^ = 0.91 for patient 6R OFF, see dark red lines in Fig 7C2 and Fig 7D2, and *R*^2^ = 0.931 for patient 4L OFF, see Fig 7C4 and Fig 7D4). The Bayesian information criterion (BIC) corresponding to the average burst duration profile fit is lower in both cases for the delayed non-linear WC than for the linear WC (ΔBIC = 12.26 for patient 6R OFF, and ΔBIC = 19.55 for patient 4L OFF). This difference in BIC highlights the superior fit of the non-linear model despite the increase in model complexity.

In summary, this subsection has demonstrated that in a biologically inspired excitatory/inhibitory (E/I) model, reproducing the most striking average burst duration profiles in the OFF condition requires the addition of delays and a non-linear (sigmoidal) activation function compared to typical ON profiles. The need for non-linearity agrees with the surrogate analysis carried out previously, but more importantly the present results also delineate the simplest type of biologically inspired model required to reproduce the two conditions studied.

In some cases, the envelope of E/I models can be related to what we call in this work “envelope models” (illustrated in Fig 1). Indeed, it was shown in [24] that the envelope of a linear WC model is a Rayleigh process, assuming the ratio of *E* to *I* envelope amplitudes is constant, and the phase delay between *E* and *I* oscillations is also constant. A Rayleigh process is in fact a particular type of envelope model. We next investigate whether a similar conclusion to that of this subsection on E/I models holds for envelope models, which provide a direct, simpler description of envelope dynamics and can more easily be fitted to all datasets. Additionally, we will study envelope models analytically to obtain additional insights.

### Investigating changes in dynamics using envelope models

To obtain the simplest polynomial dynamics describing the OFF and ON conditions, we consider envelope models directly describing the LFP envelope and fit them to data. Furthermore, to describe the dependence of burst duration on model parameters, we derive an approximate analytical expression for the average burst duration profile, and apply it to the model most representative of the ON condition.

#### Fitting envelope models to burst duration profiles

The time discretization of the simplest stochastic process with state dependent drift and uncorrelated white noise, the Ornstein-Uhlenbeck (OU) process, is in fact enough to model the average burst duration profile of the majority of ON medication datasets, and of a few OFF medication datasets. The OU process is described by the stochastic differential equation

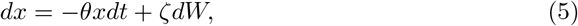

where *W* is a Wiener process, *θ* a positive decay parameter, and *ζ* the constant noise standard deviation. While *x*(*t*) directly models the envelope, the phase equation

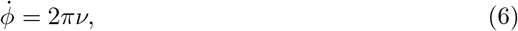

where *ν* is a constant frequency can be used to generate oscillatory activity as *z*(*t*) = *x*(*t*)cos *ϕ*(*t*).

To reproduce the average burst duration profiles of the datasets whose envelopes cannot be described by an OU process, we extend the drift function of the OU model to include non-linear polynomial terms. We are thus considering the time discretization of models of the form

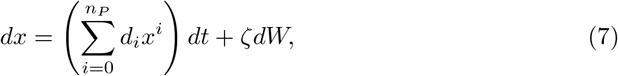

where *n*_*P*_ is the degree of the polynomial drift, and 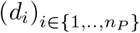 are constants with 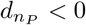.

The resulting envelope models are fitted to the average burst duration profile of all datasets. For each dataset, we fit envelope models of increasing polynomial degree starting with *n*_*P*_ = 1 (OU) until a fit to the average burst duration profile with *R*^2^ *>* 95% is achieved. We call these models “minimal models”. Model parameters are optimised (procedure in “Fitting procedure” in the Methods section) to obtain the best fit to the data average burst duration profile with *x*(*t*) directly modelling the beta envelope. With frequencies *ν* adjusted to match the data peak frequencies, the models are scaled so that the standard deviation of *z* approximately matches that of the data.

The best fits of minimal models to all datasets are shown in Fig 8. The coefficients of determination *R*^2^ of all models leading to, and including, minimal models are reported in Supplementary Tables C and D in S3. We verified that minimal models have the lowest BIC of all models considered for a given dataset (see Supplementary Tables E and F in S3). We report minimal model parameters in Supplementary Tables G, H, I, and J in S3. The fitted drift functions of minimal models (second and fourth rows in Fig 8) were found more often linear ON medication (10 cases out of 16) than OFF medication (5 cases out of 16). Barnard’s exact test [34] shows a trend (p = 0.0551, n = 16 per condition) in the medication state affecting whether the minimal model is linear or non-linear. The minimal model degree is strictly greater OFF than ON medication in 10 out of 16 cases, and this difference is statistically significant (p = 0.020, binomial test with parameters (16,0.3438)).

**Fig 8.**
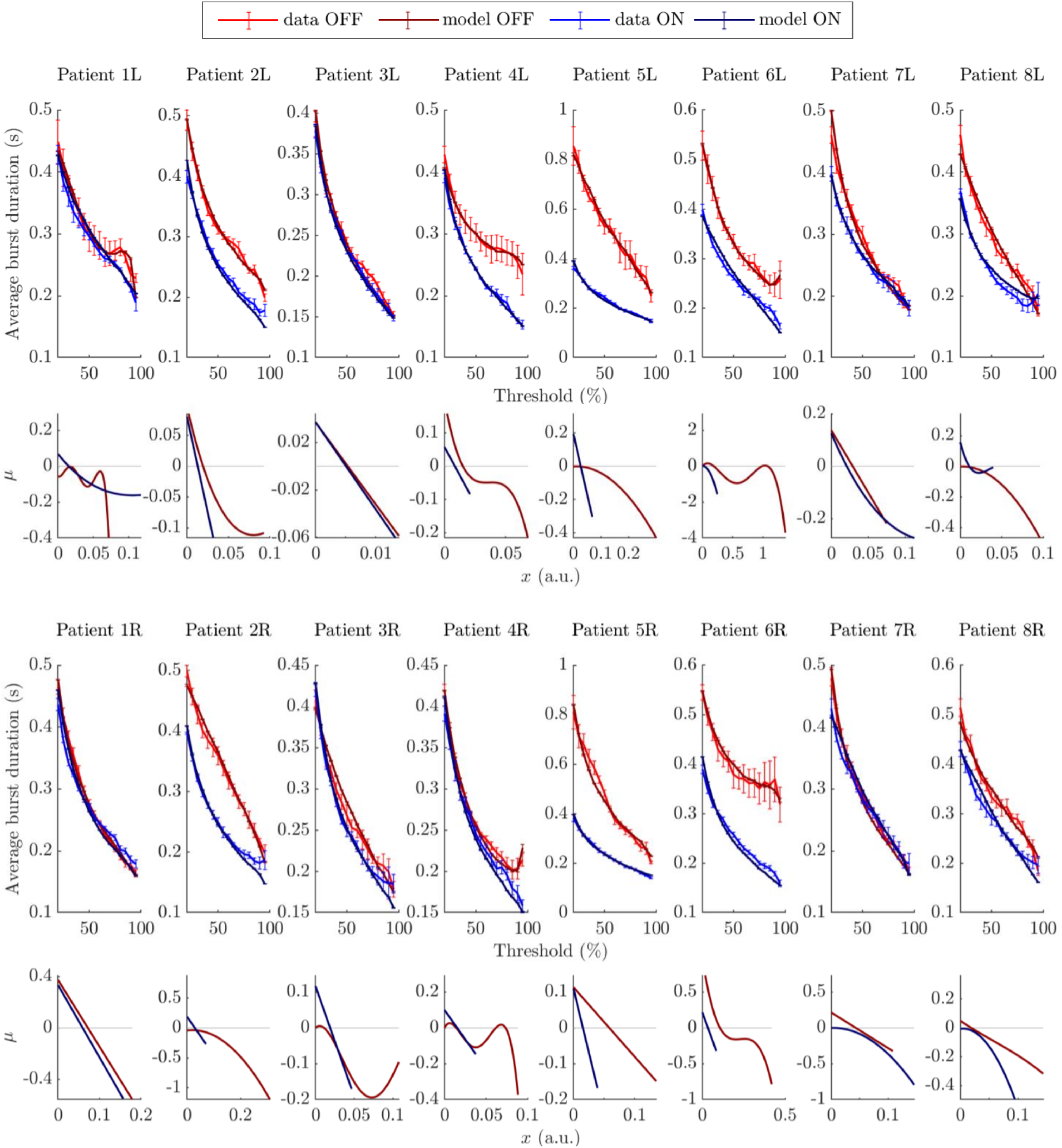
Envelope model fits to all datasets. The top two rows correspond to left hemispheres, the bottom two rows to right hemispheres, and each column corresponds to one patient. Patient’s average burst duration profiles and corresponding model fits are shown in the first and third rows (SEM error bars). The OFF state is represented in red for data, and in dark red for model fits. The ON state is represented in blue for data, and in dark blue for model fits. The drift functions (*μ*) of the fitted envelope models are shown in the second and fourth rows. The range of *x* values shown for each drift function corresponds to the range spanned by the corresponding model, and the light grey line represents *μ* = 0.

In line with the previous subsections, envelope model fits suggest that reproducing average burst duration profiles OFF medication is more likely to require envelope models with non-linear drift functions, such as polynomial drift functions of degree greater than one. We have shown that envelope models can successfully model patient average burst duration profiles, and we next study analytically how to express average burst duration profiles as a function of envelope model dynamics.

#### Average burst duration and exterior problem mean first passage time

In order to study the dependence of burst duration on envelope model parameters, we seek an analytical expression for the average burst duration profile of envelope models. As recently noted in [24], the concept of average burst duration is related to the concept of mean first passage time (MFPT, also known as mean exit time) in the exterior problem. Considering a threshold or boundary *L* and a starting point *x*_0_ = *L* + *δ*, with *δ >* 0, we denote the continuous-time MFPT from *x*_0_ to *L* by 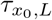. It is defined by the expectation of the random variable inf {*t ≥* 0 : *x*_*t*_ ≤ *L*}. At this point it should be clarified that the average burst duration of a continuous-time stochastic process with Gaussian noise is always zero, as all trajectories starting from the threshold *L* will cross it again in vanishingly short times. In addition, MFPTs in continuous time are systematically biased towards shorter first passage times since a continuous stochastic model is more likely to cross the boundary when close to it than its discrete-time counterpart. However, we can adapt classical MFPT results from continuous-time stochastic process literature to analytically approximate the average burst duration of the corresponding models in discrete time. In what follows, we use a tilde to distinguish quantities that can be measured readily in a time discretized system from the continuous-time quantities.

To study the associated discrete-time model with time step *dt*, we are considering the continuous-time case of a stochastic process

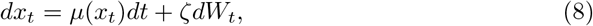

where we have assumed that the real valued drift function *μ* can depend on *x*_*t*_, and that *ζ* is constant in time and space. Following a derivation similar to the derivation reviewed in [35], we show in the Methods section (see “Continuous model MFPT”) that the MFPT of the continuous system is given by

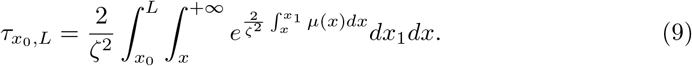

For the integral to converge, it is necessary that *μ*(*x*) is non-vanishing and negative at +∞, which is the case in models describing neural oscillations. Equation (9) cannot be applied to white noise only, where *μ*(*x*) = 0.

From equation (9), we derive an expression for the average burst duration of the discretized model with time step *dt*. We show in the Methods section (see “Discretized model”) how to correct for the systematic bias of the continuous model (Steps 1 and 2). We also detail therein how to relate average burst duration and MFPT (Step 3): when crossing a threshold *L* from below at the start of a burst, a discrete model will always overshoot the threshold, and the average burst duration at threshold *L* can be related to MFPTs by taking into account the overshoot distribution at *L* (see Fig 9). We finally obtain to first order in 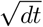 the average burst duration 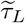 for the discrete-time model as

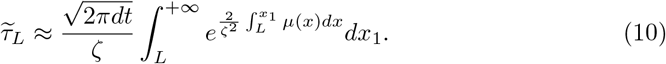

**Fig 9.**
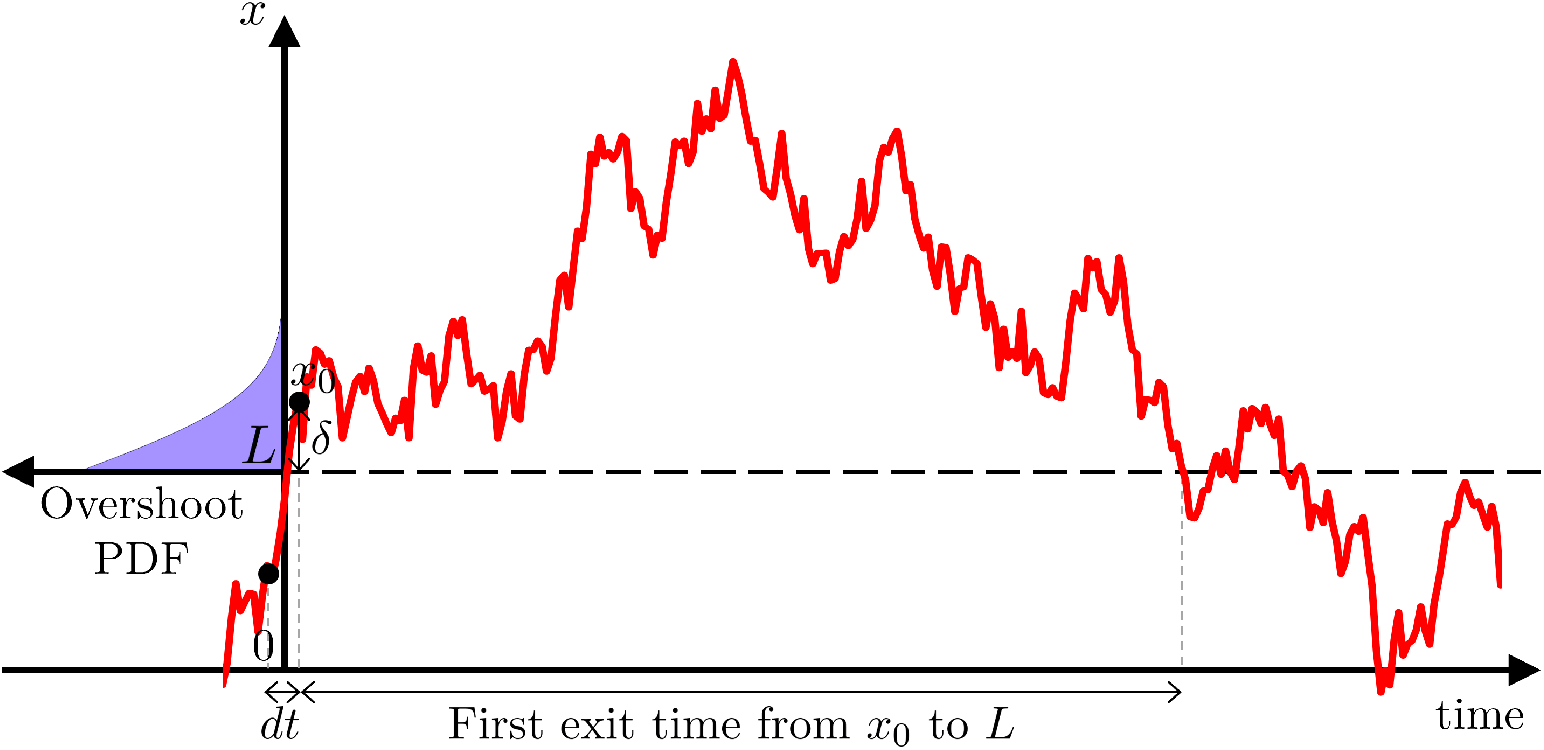
First exit time and overshoot distribution. Showing a discrete realisation of an OU process overshooting the threshold *L* by *δ*, with a sketch of the overshoot probability density at *L* in purple. The first exit time from *x*_0_ to *L* is the time taken to get below *L* for the first time when starting from *x*_0_.

Equation (10) provides a general relationship from drift function to average burst duration profile for the discretization of stochastic envelope models described by equation (8). As shown in Fig 10, this result applied to the OU model and a third degree polynomial envelope model (equation (11) below and equation (48) in the Methods section) is very close to simulations.

**Fig 10.**
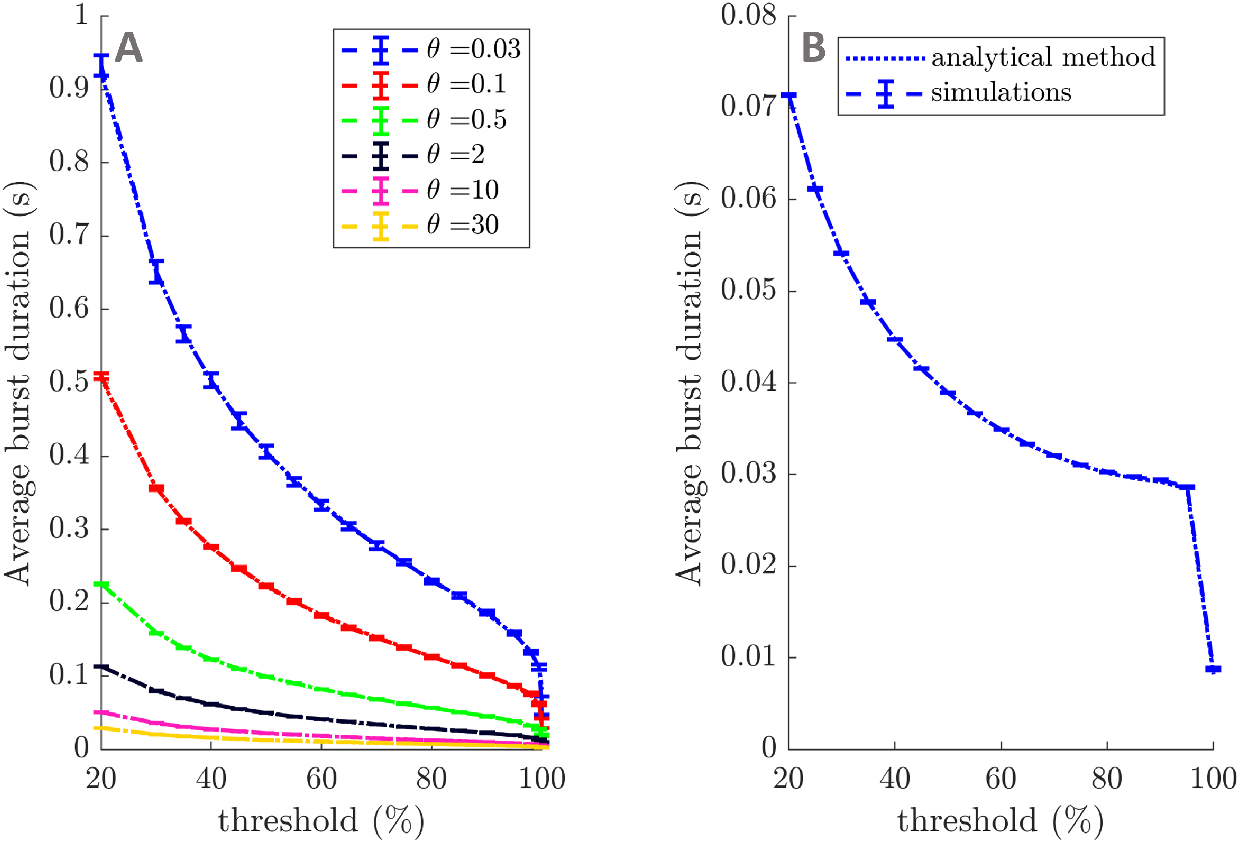
Simulations of average burst duration profiles for OU processes and a third degree polynomial model. Average burst duration profiles from simulations of OU processes are compared to equation (11) for a range of decay parameters and *ζ* = 1 in panel A. Similarly, average burst duration profiles from simulations of a third degree polynomial envelope model are compared to equation (48) in panel B. Simulations are indicated by dashed lines (SEM error bars), and analytical results by dotted lines. Simulations consist of five repeats of 10^5^ s, with a time step of 1 ms (OU process simulations use the exact updating equation (17)).

#### Increasing average burst duration in the OU model

Building on the previous result, we highlight the dependence of average burst duration on model parameters and show how bursts can be lengthened in the discretized OU model, which is representative of most datasets in the ON condition and of linear systems in general (see Supplementary Fig K in S2). We are considering here the discretization of an OU process centered on zero (equation (5)). Equation (10) gives, to first order in 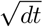

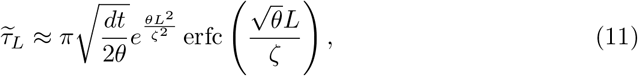

where erfc is the complementary error function. The dependence of average burst duration on parameters is easier to see when *L* is expressed as a percentile of the time series values, which is also how we plot average burst duration profiles. Because of this choice, the shift introduced to ensure the envelope stays positive (see “Fitting procedure” in the Methods Section) will have no impact on the result.

We express the average burst duration of a discrete OU process as a function of threshold percentile using the stationary probability distribution of the OU process (see “OU” paragraph in the Methods section). We obtain

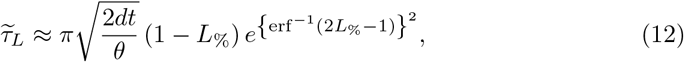

where *L*_%_ is the percentile rank (between 0 and 1) corresponding to threshold *L*. Equation (12) does not depend on the noise parameter *ζ* (which only scales trajectories), and makes apparent a simple dependence of the average burst duration on the decay parameter. We can conclude that increasing the average burst duration for a given threshold corresponds to decreasing the decay in the OU model, which is what is intuitively expected.

We have demonstrated that discretized envelope models can reproduce the average burst duration profile of all the datasets available with *R*^2^ *>* 95%, and we have provided the polynomial forms of the drift functions required. In addition, the average burst duration profile of discretized envelope models have been approximated analytically, which yields insights into how model parameters influence average burst duration profiles. This analytical result opens the question of a reverse link from average burst duration profile to dynamics, which we investigate next.

### Average burst duration profiles are a signature of envelope dynamics

In this subsection, we establish that average burst duration profiles are a signature of envelope dynamics by showing that the envelope drift function can be recovered from the average burst duration profile. This clarifies how a change in the temporal patterning of beta oscillation signals a change in dynamics.

#### Relationship between envelope drift function and average burst duration profiles

To highlight the importance of average burst duration as a marker of dynamics, we establish a link for a general class of stochastic processes with additive noise between average burst duration, noise standard deviation, and dynamics. We are considering a time discretization (time step *dt*) of the stochastic process given by equation (8). As a reminder, we assume that the real valued drift function *μ* can depend on *x*_*t*_, and that the diffusion term *ζ* is constant in time and space. We show in “Passage method” in the Methods section, that to first order in 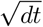,

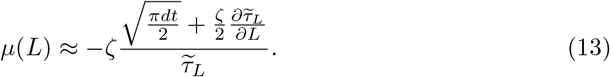

Equation (13) highlights a direct relationship from average burst duration in discrete time to envelope dynamics (drift function). This is essentially a local relationship (at threshold *L*), and the drift function can be estimated where the average burst duration profile is known. We call this estimation procedure the “passage method”, and show next that it can recover envelope dynamics in synthetic data and patient data.

#### Inferring dynamics in synthetic data and in patient data with the passage method

To validate the method, we first test the passage method on synthetic data. We are considering envelope time series of 250 s (roughly the same length as patient data) and of 1000 s generated from an OU envelope model and a fifth degree polynomial envelope models. Dynamics from these time series are recovered using the passage method. We are considering 300 thresholds, and applying smoothing to 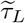 and its derivative (full details provided in “Passage method” in the Methods section). The results are shown in the first two rows of Fig 11. Approximations of the drift functions *μ* are recovered (Fig 11A1 and Fig 11A2), and the average burst duration profiles and inverse cumulative distribution functions (CDFs) obtained from simulating the recovered dynamics approximate the synthetic data well (Fig 11B1 and Fig 11B2, and Fig 11C1 and Fig 11C2, respectively). Recovered dynamics and associated features are better approximations when recovered from the longer time series than from the shorter ones. Performance is however reasonable for the shorter time series, which are approximately the duration of patient data recordings.

**Fig 11.**
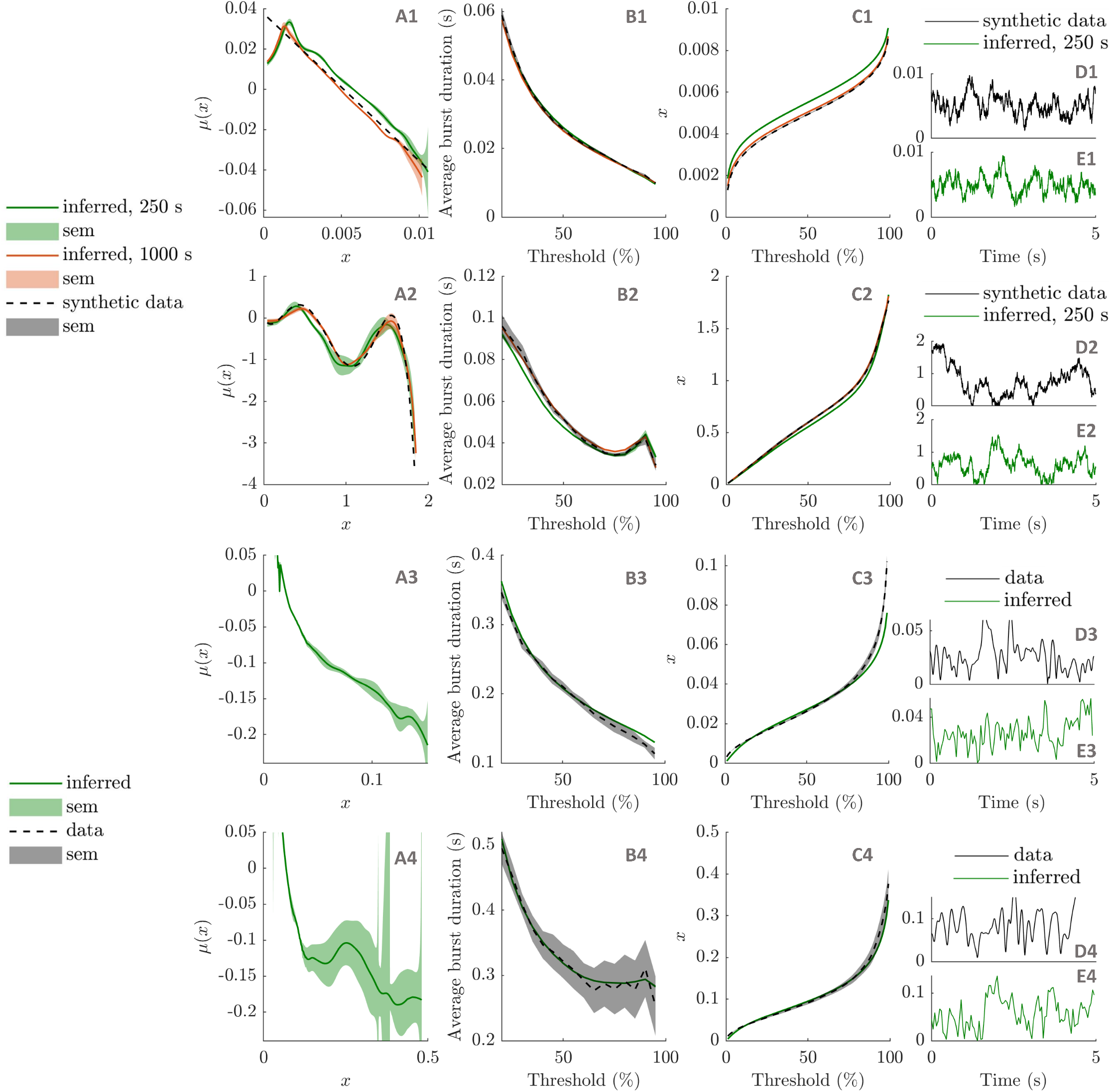
Inferring envelope dynamics with the passage method. The method is applied to synthetic data (OU model in the first row, fifth degree polynomial model in the second row), and patient data (patient 6R, ON medication in the third row, and patient 6R, OFF medication in the last row). The recovered drift functions (*μ*) are shown in the first column, and the average burst duration profiles and the inverse CDFs simulated from the recovered dynamics in the second and third column, respectively. Ground truths are provided when available (black dashed lines). Synthetic data results are presented for 250 s and 1000 s of synthetic data. In the last column, 5 s of simulated data from the recovered dynamics are compared to training data.

Given its success on synthetic data, we test the method on patient data. We pick the patient that scores the highest on our burst duration specific non-linearity measure, patient 6R OFF, and the same patient and hemisphere ON medication for comparison. The method is applied as before, with the additional estimation of the noise parameter and the time step (specifics of the application of the method to patient data detailed in “Passage method” in the Methods section). The results are shown in the last two rows of Fig 11. There is no ground truth to compare the recovered dynamics against (Fig 11A3 and Fig 11A4), but the average burst duration profiles and inverse CDFs obtained from simulating the recovered dynamics are a good match to the data. We note that the dynamics inferred ON medication are closer to a straight line, as reported for this dataset in the previous subsections.

#### Comparison of the passage method with a direct method

To evaluate the performance of the passage method in extracting envelope dynamics from data, we compare our method with a simpler envelope recovery method that directly estimates time derivatives from the envelope time series. Specifically, time derivatives are calculated from consecutive envelope values using first order differences. The resulting derivative estimates are binned in 300 bins of equal width (same number of bins as the number of thresholds in our method) within which derivative values are averaged.

Both the simple method and the passage method are applied to envelope time series of a range of durations from 100 s to 1000 s generated by the OU model and the fifth degree polynomial model used in previous tests. The passage method is applied as described in “Passage method” (Methods section), except that no smoothing is applied to either the passage method or the direct method for a fair comparison. The sum of the squared errors between the recovered drift function *μ* and the corresponding ground truth is calculated for both methods for each time series, and the results are presented in Fig 12. The passage method is at a slight advantage for a range of time series lengths in the OU case, and for all time series lengths in the fifth degree polynomial case.

**Fig 12.**
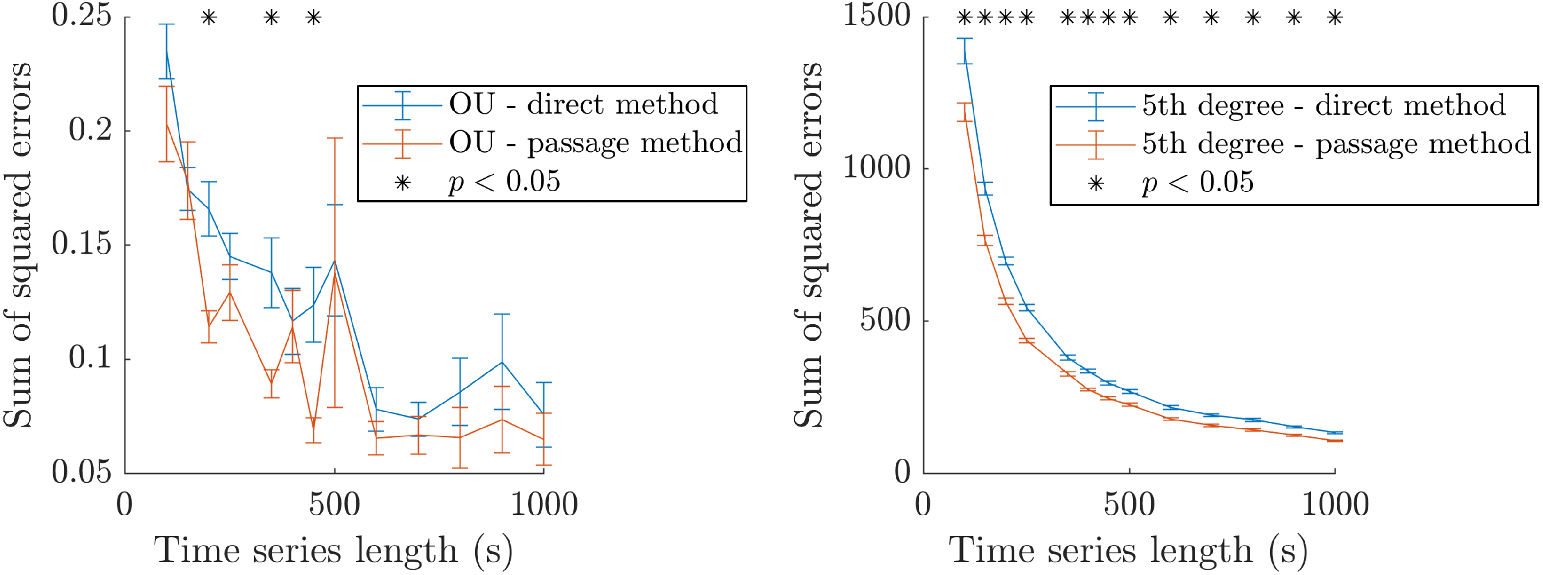
Comparison of the passage method and the direct method on synthetic data. Showing the sum of squared errors between the inferred and ground truth *μ* functions for both methods as a function of the duration of synthetic data used for inference. The synthetic data were generated from an OU process (left) and a fifth degree polynomial (right). In both cases, the mean value of the sum of squared errors and the SEM error bars are obtained for a large number of repeats (150 repeats from 100 s to 250 s, 100 repeats from 350 s to 450 s, and 50 repeats from 500 to 1000 s). Significant differences are highlighted by black stars (t-tests under FDR control, same sample sizes as for error bars).

## Discussion

In this study, we analysed PD patients’ STN LFP recordings in the beta band and first motivated the choice of average burst duration as a marker of the differences in bursting dynamics ON and OFF medication. We found a burst duration specific measure of non-linearity based on linear surrogates to significantly increase from the ON to the OFF medication state, and the change in this non-linearity measure to be correlated with motor impairment. We further narrowed-down dynamical changes underlying the changes in beta oscillation temporal patterning between medication states by fitting models to data. The simplest biologically inspired neural mass model to reproduce typical ON average burst duration profiles was a WC model with a linear activation function and uncorrelated inputs, while the most striking OFF profiles required the addition of a non-linear activation function and of delays. Envelope models were fitted to all datasets, and reproducing average burst duration profiles OFF medication was more likely to require non-linear polynomial drift functions, while the majority of ON medication datasets could be modelled with linear drift functions (OU processes). The simplicity of envelope models enabled us to derive an approximate expression for average burst duration profiles (equation (10)). This expression clarified how model parameters affect average burst duration in the linear case. In addition, we showed that the non-linearity can be directly extracted from average burst duration profiles using our “passage method” (based on equation (13)), demonstrating that average burst duration profiles are a signature of envelope dynamics. This sheds light on why burst duration has been suggested in multiple studies as an important biomarker in PD.

### Bursting features

Average burst duration profiles are an insightful and exhaustive way of characterising bursting dynamics. Average profiles across thresholds benefit from information from a range of thresholds and capture more about the system than the burst duration distribution at one threshold (which may only integrate dynamical information, and predominantly above the threshold). In fact, we demonstrate that average burst duration profiles are closely related to average burst rate profiles, and more importantly, to the dynamics of the beta envelope. Some previous studies have used wavelet amplitude as the beta envelope to study bursting features [12, 13], while we used the Hilbert envelope of the filtered signal. We found little difference between the methods as far as average burst duration is concerned (not reported). While a number of recent studies define bursts as Hidden Markov Model states [36, 37] or using support-vector machines [38], we aim at providing a more complete dynamical picture of beta dynamics. If both STN LFP and single unit activity are available, the temporal structure of beta synchronisation can be investigated using first-return maps of the signal phases [39].

Besides the PSD, our statistical analysis of patient STN LFP recordings identified average burst duration across thresholds as the analysed bursting feature most suited to distinguish between burst dynamics in the ON and OFF states. We highlight the importance of individually z-scoring the filtered data when comparing average burst amplitude profiles between medication states to control for differences simply due to differences in mean beta power. A similar result has been reported in a non-human primate study [18] where beta burst duration was found to be a better differentiator of healthy and pathological episodes in the STN than burst amplitude. Deleting longer beta episodes resulted in a greater decrease of parkinsonian activity than deleting stronger episodes. In our patient analysis, the ON medication state gives an approximation of the physiological state. It is hypothesised that shorter bursts are more likely to be physiological, whereas longer burst are more likely to be pathological [12, 13]. Longer bursts are known to be associated with more synchronization in the STN, but also across the motor network [40]. This results in less information coding capacity [7, 8] and may underlie the motor symptoms.

### Surrogate analysis

Our burst duration specific measure of non-linearity, BDDLS, provides a new dimension that can be used to analyse LFP recordings. Non-linearity here should be understood as non-linear correlations in the time series of interest (a sinusoid is considered linear, as it can be represented by an ARMA model, see “Linear surrogates”). A high BDDLS can come from a longer average burst duration for all thresholds than what would be expected for a linear system (e.g. dataset 2R OFF), and/or a change in burst duration distribution for certain thresholds only (e.g. dataset 6R OFF, and see Fig 2 where the proportion of short bursts is lower at high thresholds).

We showed that, in the case of PD, the difference in BDDLS between medication states in STN LFPs is correlated with motor impairment. In our small patient sample, the correlations for *ρ* = 0.1 to 0.5 were greater than 0.73, while the correlation obtained with the relative difference in power and the difference in burst duration were 0.500 and 0.476, respectively. However, given the uncertainty arising from the small sample size, the correlation between BDDLSdiff and motor impairment may have been overestimated. Therefore, we do not claim that BDDLSdiff is superior to PSDdiff or DURdiff in predicting motor impairment. The correlation between BDDLSdiff and motor impairment should be re-evaluated on larger datasets. Nevertheless, predicting burst duration based on PSD alone will not be accurate for patients that have different average burst duration profiles than their linear surrogates, which is the case for most patients OFF medication. This is because the PSD only reflects linear correlations.

The lower signal to noise ratio (SNR) ON medication may play a role in the difference in BDDLS observed between medication states. However, if the difference in SNR between medication states were the primary driver of BDDLSdiff, we would expect BDDLSdiff to be strongly correlated with PSDdiff. This is not the case as highlighted by the left panel in Supplementary Fig J in S2 (Spearman’s correlation is 0.500, p = 0.216 for *ρ* = 0.2).

Changes between the ON and OFF states can be classified into three types. The first one is mostly “linear” changes (e.g. patient 5R, in Fig 4, where both the ON and OFF average burst duration profiles are close to their surrogates and there is a large difference between surrogates). The second one is mostly “non-linear” changes (e.g. patient 4R, in Fig 4, where the ON profile matches its surrogate and the OFF surrogate, and the OFF profile differs from its surrogate). The third one is both “linear” and “non-linear” changes (e.g. patient 6R, in Fig 4, where the ON profile is close to its surrogate, and there is a large difference between surrogates and between the OFF profile and its surrogate). These categories could provide a basis to stratify patients, in particular if similar results could be obtained from non-invasive EEG recordings.

The scaling by the mean value of the surrogate profile that was applied to BDDLS helped decorrelate BDDLSdiff from the difference in burst duration (DURdiff). We defined DURdiff as a difference between data burst duration profiles for simplicity, but it may be the case that defining DURdiff as the difference between linear surrogates would decorrelate BDDLSdiff and DURdiff further and would improve predictive power when all three metrics are considered. Given more data, the contribution of the metrics to predict motor impairment should be further assessed, for instance by comparing multivariate regressions based on various combinations of the metrics.

Surrogate testing has been used to explore non-linearity in brain recordings in pathological states. In particular it has been reported that epileptic seizures are associated with non-linear brain dynamics, while little non-linearity was found in Alzheimer’s disease (reviewed in [41]). In the context of PD, we found an increase in a burst duration specific measure of non-linearity OFF medication compared to ON medication. From an information theoretic perspective, non-linear correlations between frequencies within the beta band may further reduce information coding capacity, and impair motor function. Non-linear correlations between different frequencies in STN LFP rhythms have in fact been reported to be greater OFF than ON medication [42]. On a related note, scalp EEG recordings over the sensorimotor cortex of PD patients were found to have more pronounced non-sinusoidal features in the beta band OFF medication compared to ON medication [43] and OFF DBS compared to ON DBS [44]. In addition, non-linearity has been identified in inter-spike interval series in PD patients OFF medication [45]. STN LFPs were also described as more non-linear during resting tremor in PD [46], but our non-linearity measure is specific to burst duration in the beta band and may therefore be more specific to bradykinesia and rigidity. Very recently, a non-linearity measure related to a higher order version of auto-correlative signal memory was found to be greater OFF than ON medication in STN LFPs filtered in the beta band [47].

Non-linear correlations are not seen in average power, and we showed that the degree of non-linearity is correlated with motor impairment, which has implication for aDBS in PD patients. A slow variant of aDBS provides stimulation according to average beta power on a 50 s time scale [48, 49]. Slow aDBS will not address selectively the aspect of the pathology we highlighted in this study, since average power cannot reflect non-linear correlations. Providing aDBS according to a predictive algorithm based on the frequency information contained in a 300 ms window before burst onset was reported in PD to be very close in performance to optimised aDBS, i.e. to threshold based stimulation [50]. Moreover, it appears that pathological synchronization is established earlier than common thresholds used to define beta bursts [51]. Non-linear information not present in the windowed power spectra taken as features in [50] may provide additional predictive power, and linear surrogates estimated on a slower time scale than real time could provide a useful baseline to compare current recordings against. The computational cost of GWR surrogates rules them out, but computationally cheaper IAAFT surrogates may be better cut out for the task. Although GWR surrogates were used in our study to account for potential nonstationarity in LFP recordings, a correlation with clinical scores was observed for *ρ* = 0, which corresponds to IAAFT surrogates. Average burst duration profiles and BDDLSdiff have been studied on a slow time scale (about 250 s of data). How little data can be used to reliably estimate BDDLSdiff remains to be explored, and other surrogate based non-linearity measures may be more suited to real-time use. Nevertheless, closed-loop DBS targeting non-linearity in the drift function of beta envelope models might prove more selective in suppressing pathological oscillations than closed-loop DBS approaches based on amplitude. Additionally, amplitude thresholds in aDBS would need to change with medication and activity level, and a stimulation strategy not requiring an amplitude threshold would be simpler.

### Modelling burst duration profiles

When fitting models to patient data, model complexity was increased gradually to avoid overfitting and identify the simplest models that could fit the data. Linear models could fit to the simplest average burst duration profiles. The linear models chosen were a linear WC model without delays and an OU envelope model. On the one hand, the architecture of the former can be mapped onto the STN and GPe populations [31–33], and it only has a few parameters, making it easier to constrain and less likely to overfit the data. On the other hand, the latter is perhaps the simplest stochastic envelope model with a non constant drift term, and is a simplification of the Rayleigh envelope model, which has been shown to describe the envelope of the linear WC model under certain assumptions [24] as mentioned in the Results Section. To model more complex average burst duration profiles (higher burst duration specific measure of non-linearity), complexity was introduced gradually in neural mass models and envelope models. It was verified that a linear WC model without delays could not fit the data, and that a non-linear WC model without delays could not fit the data either (not shown), before successfully fitting the non-linear WC model with delays. With envelope models, polynomial drift functions of increasing degree were fitted to the data until satisfactory minimal models were obtained.

Dynamical changes identified in neural mass models and envelope models between medication states deserve further discussion. While it could be tempting to conclude from the WC fits that delays are necessary to reproduce the data from patients 6R OFF and 4L ON, the corresponding third degree polynomial envelope models suggest that non-linearity is enough to reproduce the data and that delays are not necessarily required in envelope models. Changes in WC model parameters between medication states cannot be interpreted directly because of non-identifiability (many sets of parameters can lead to the same output). The levelling-off at high thresholds of several average burst duration profiles OFF medication (e.g. 1L OFF, 4L OFF, 6R OFF) could be explained by larger oscillations getting stability from an additional attractor. Multistability could be a dynamical interpretation of such features in the average burst duration profile, and could open the door to a dynamical definition of bursts as events in the vicinity of an attractor at larger amplitude. However multistability turns out to not be a requirement since the minimal models fitted to datasets 1L OFF, 4L OFF, and 6R OFF do not exhibit multistability (horizontal axis not crossed, light grey line in Fig 8) but produce average burst duration profiles close to the data. This is also the case for the dynamics inferred by the passage method applied to dataset 6R OFF (see Fig 11A4, and Fig 11B4). While the “bumps” or levelling-off seen in the drift functions of datasets 1L OFF, 4L OFF, and 6R OFF are not indicative of multistability, they suggest an attractive influence at the corresponding amplitude level. This attractive influence brings more stability to oscillations of the corresponding amplitude. It is unclear how this attractive influence could be interpreted in terms of physiological processes at this point. Multistability is however present in the minimal envelope models fitted to datasets 6L OFF and 4R OFF. In contrast, we note that in the ON condition, drifts of minimal envelope models are either straight lines or are shaped like simple parabolas and do not display the more complex features observed in the OFF condition.

The classification of changes between the ON and OFF states in three types (“linear” changes, “non-linear” changes, and a combination of both) mentioned when discussing the surrogate analysis are reflected in the fitted envelope models. For patient 5R (given as an example of mostly “linear” changes), the minimal models representing the ON and OFF states are both linear, but with different slopes. For patient 4R (given as an example of mostly “non-linear” changes), the ON state is represented by a linear model, and the OFF state by a fourth degree polynomial drift. The ON and OFF drift functions significantly overlap, and the OFF drift function adds non-linearity on top of the ON drift function. The ON state could be thought of as a sub-regime of the OFF state. For patient 6R (given as an example of both combined “linear” and “non-linear” changes), the ON state is represented by a linear model, and the OFF state by a third degree polynomial drift. The ON and OFF drift functions do not overlap.

The drift function and noise parameter of a minimal envelope model characterise the system generating the observed beta oscillations in a concise manner. In this study, we have identified differences in the drift function of minimal envelope models between medication states in PD patients. The drift function and noise parameter are however more than just features of the beta state, and can be used to generate synthetic beta envelopes with average burst duration profiles (and equivalently average burst rate profiles) similar to the data, which could contribute to in-silico design and testing of closed-loop DBS strategies.

It is well known that simple WC models of the STN-GPe loop can generate sustained beta oscillations without the need for correlated inputs [33, 52], but whether that is the case for transient beta oscillations has not been reported. The finding that a simple STN-GPe WC model with uncorrelated noise as inputs can reproduce the top two most complex average burst duration profiles of the dataset, i.e. realistic transient beta oscillations, is consistent with the theory that the STN-GPe loop could play a predominant role in the generation of pathological beta activity [53–55]. There is no consensus on how exaggerated beta oscillations arise, and other theories include excessive beta oscillations originating in the longer cortico-BG-thalamic loop [33, 56], or directly within interconnected populations of striatal neurons [57]. Our WC model does not exclude a role of influences external to the STN-GPe, such as cortical and striatal influences. Indeed the model receives constant inputs from these populations as well as white noise. However, reproducing average burst duration profiles ON and OFF medication did not require correlated external inputs.

### Analytical expression of average burst duration in envelope models

Standard first-passage results are available for continuous-time stochastic processes (see for instance [35]), but their application to time-discretized systems is not immediate. While we derive in this study a three step correction to obtain the average burst duration profile of envelope models in discrete time (assuming time-independent but possibly state dependent drift function and constant diffusion term across space and time), other related approaches are worth mentioning. Perhaps the most relevant is a continuity correction introduced in the financial literature to price continuous barrier options [58]. The correction consists in shifting the continuous barrier to account for discrete monitoring of prices, but is not applicable when the initial price is close to the barrier. As a result, this approach cannot be successfully used for bursts, since they start very close to the barrier (the distribution of initial points is given by the overshoot distribution). Alternatively, algorithms have been suggested to approximate continuous first passage time distributions with simulations [59–61]. In the specific case of the OU model, approximations are directly available for the first passage density of the discretized model, but their complexity makes them unpractical [62, 63].

Discrete models are required to model burst duration, however the dependence of our results on the time step is worth discussing. As mentioned in the Results Section, discrete models are required to model burst duration since the average burst duration for a continuous stochastic process with Gaussian noise is always zero. As expected, the average burst duration given by equation (10) goes to zero as the time step goes to zero. Moreover, the dependence of average burst duration on the time step predicted by equation (10) matches what is observed when simulating the OU and third degree models used in Fig 10 for various time steps (not shown). Importantly, the square root of the time step is only a constant factor that scales the average burst duration as seen in equation (10), and it does not influence the shape of the profile. The time step can be seen as a parameter of the bursting model, and can be chosen to match the roughness of the data envelope, which is done when applying the passage method to data. We also highlight that our analytical approach differs from our experimental approach in that the minimum burst duration of 100 ms considered for data analysis and fitting is not modelled.

Mean first passage time analysis in the context of gamma bursts has been used in [24], where an expression is given for the mean duration of a typical burst in a Rayleigh envelope model. A typical burst is defined therein as starting and ending at the envelope median, and peaking at the envelope mean plus one standard deviation. The present work contributes a result for a more general class of models, without relying on an arbitrary burst threshold and peak. Our result also includes the influence of time-discretization, and the dependence on the threshold used to define bursts. Equation (10) is applicable to the time-discretization of a Rayleigh model 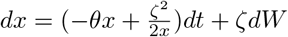, and the corresponding average burst duration at threshold *L* is simply 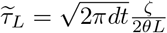.

It is noteworthy that a simple analytical expression could be obtained for the average burst duration profile of OU processes (equation (12)), making the analysis of the influence of model parameters on average burst duration possible. Besides thresholds, only the decay parameter and the time step play a role in the expression. For a given time step, it is therefore possible to identify an envelope decay in a dataset by fitting an OU average burst duration profile to the dataset’s average burst duration profile. The decay could be assimilated to the real part of the eigenvalues of a broader class of linear envelope models. In the context of simple integrate and fire neuron models, firing can be related to the first passage of an OU process, and the associated decay parameter is estimated in [64] by considering inter-spike interval statistics. In our work, Equation (12) clarifies the role played by the decay parameter, and predicts that a linear system modelling beta bursts will be made more pathological when its decay parameter is reduced, which makes intuitive sense. This may correspond to changes in average burst duration described as “linear” in the “Surrogate analysis” part of the discussion. Additionally, since we have expressed the average burst rate as a function of the average burst duration in S1, we provide an approximate analytical expression for the average burst rate of OU processes (equation (S.4) in S1). Burst rate is also of interest as it is used to characterise bursting in some experimental studies (see for example [15, 65, 66]). In the third degree model analysed in Fig 10, dependence on model parameters cannot be studied directly contrary to the OU case, but the integral expression given by equation (48) could facilitate numerical analysis as it can be evaluated numerically much faster than the model can be simulated and subjected to burst duration quantification. Also of experimental interest, the burst duration distribution for a given threshold can be obtained analytically for OU processes as an infinite series, but is hard to evaluate numerically since it involves confluent hypergeometric functions of the second kind, their zeros and derivatives (equation (90) in [35]).

### From average burst duration profiles to dynamics

The link we established from average burst duration profiles to drift function suggests that average burst duration profiles are a window into envelope dynamics. As detailed in the introduction, burst duration has been experimentally identified as an important marker of pathology for PD, and our finding sheds some light on the dynamical significance of burst duration. The uncovered link (equation (13)) is valid for envelope models with a general drift function that can depend on the position, and additive noise constant in time and space (“complicated dynamics, simple noise”). However, the model cannot include delays, and dynamics cannot be recovered in the vicinity of the derivative of the MFPT being zero at the threshold considered. As is manifest in equation (13), when the average burst duration and its derivative are considered only at a given threshold, only local information on the dynamics is available.

The relationship naturally provides a method to infer envelope dynamics from data, which we call the “passage method”. We applied the passage method successfully to synthetic data, and recovered envelope dynamics from patient data for patient 6R, ON and OFF medication. In this dataset, envelope dynamics OFF medication were found more non-linear (a straight line fit would be worse) than ON medication. Comparing the inferred drifts (Fig 11A3 and 11A4) to drifts of the corresponding minimal models obtained from fitting (Patient 6R in Fig 8), shapes are similar, and differences in the scale of *μ*(*x*) come from differences in the time steps used. Other differences may come from the fact that the minimum burst duration of 100 ms considered when fitting envelope models cannot be modelled in the passage method.

The passage method favorably compared to a simple, direct method in the absence of smoothing. The slightly better performance of the passage method on synthetic data may be related to a better robustness to noise. The noise parameter has to be estimated and is accounted for in the passage method, which is not the case in the direct method. This might be advantageous provided that constant additive noise is a good approximation for the dataset at hand. Comparison of the passage method with state of the art dynamics inference methods that account for stochasticity such as dynamical Bayesian inference [67–69] or Langevin regression [70] is out of the scope of this paper, but would provide more insight into the performance of the passage method. Dynamical Bayesian inference and Langevin regression were not specifically designed to recover envelope time series, and assume basis functions to represent dynamics, but could be applied to envelope dynamics inference. Additionally, the direct method can be improved using multiple linear regressions and derivative estimates with various time lags [71], in particular for linear dynamics.

Beyond movement disorders, the bidirectional link between average burst duration and envelope dynamics may find productive applications in other parts of the neurosciences and beyond where transient oscillatory dynamics are of interest. In particular, beta and gamma bursts [65], as well as the duration of sharp wave ripples [72] have been found to be related to memory. State anxiety was shown to be related to beta burst duration changes in the sensorimotor and prefrontal cortex [73]. Perception [11] and pain [74] are some other examples of fields where the importance of transient neural oscillations has been demonstrated.

## Methods

We present in this section some methodological details on data analysis, model fitting, the derivation of average burst duration in envelope models, and the passage method.

### Extracting power spectra and bursting features

We extracted the PSD and the three bursting features considered (average burst duration profile, average burst amplitude profile, and envelope amplitude PDF) from previously reported bilateral STN LFP recordings of eight patients with advanced Parkinson’s disease ON and OFF Levodopa [2]. Recordings were obtained three to six days postoperatively, while leads were externalized. Patients were withdrawn from antiparkinsonian medication overnight, and recordings were performed at rest (patients quietly seated). In the ON medication condition, a dose of Levodopa was administered about one hour prior to the recordings. Recordings ranged from 137 s to 366 s (mean duration 233 s) and were obtained from adjacent contact pairs. Signals were amplified and filtered at 1–250 Hz using a custom-made, high-impedance amplifier, and sampled at 625 Hz or 1 kHz.

Features were obtained from filtered data. For a given patient and hemisphere, the contact pair with the highest beta peak was selected, and both the ON and OFF medication data were band-pass filtered ± 3 Hz around the beta peak found in the OFF state (defined as the power spectrum maximum in the 13 to 35 Hz range). Power spectra were directly obtained from the filtered data (power spectra of both filtered and unfiltered data are shown in Supplementary Fig L in S2). As in [12], we exclude one patient reported in [2] who had an outlier response to treatment with Levodopa (relative difference between filtered PSD OFF and ON medication 2.5 standard deviations from the mean, more beta power ON medication). Next, the filtered data were individually z-scored to remove amplitude differences that could arise simply from a difference in mean beta power between the ON and OFF states, and highlight instead any possible differences in the temporal dynamics of beta amplitude. This z-scoring step has no effect on burst duration profiles as thresholds considered are individual to each time series. Beta envelopes were obtained as the smoothed modulus of the analytic signal of the filtered, z-scored data (smoothing span of 5 ms, about a tenth of a beta cycle). To allow for statistical analysis, time-series were then divided into five parts of equal length. For each segment, beta envelope PDFs were estimated, and burst duration and burst amplitude profiles were built as the average burst duration and amplitude for thresholds ranging from the 20^th^ to the 95^th^ percentiles of the envelope in steps of 5%. This range includes the typical 75^th^ percentile used in previous studies [12–15]. Thresholds lower than the 20^th^ percentile were not included since they are less characteristic of bursting (the measured average burst duration at the 0^th^ percentile is the duration of the recording). The 100^th^ percentile was left out as it corresponds to an average burst duration of zero and an average burst amplitude given by the maximum value of the envelope (only one value). We used the definition of bursts given in “Choice of bursting features” (including the minimum burst duration of 100 ms), and burst amplitude was defined as the maximum amplitude recorded during a burst.

### FDR control

Multiple statistical tests were performed in this study under FDR control at 5%. This ensures that the expectation of the number of false positives over the total number of positives is less than 5% when many statistical tests are performed. The null hypothesis was rejected relatively frequently in this study (for example 13 out of 25 times in the linear surrogate analysis section). Thus, the total number of tests is not a good estimator of the number of true null hypotheses in this study. The total number of tests is the estimator used in the original Benjamini and Hochberg procedure [75]. Instead, we rely on a better estimator of the number of true null hypotheses [76], and use an FDR control procedure based on it (adaptive linear step-up procedure, reviewed in [77]).

### From FT surrogates to GWR surrogates

FT surrogates are the most straightforward implementation of the linear surrogate idea: the Fourier phases of the data Fourier transform are randomized, but its Fourier amplitudes are kept, which ensures that upon inverse Fourier transform, the generated time series will share the same linear properties as the data but none of their potential non-linear properties. This corresponds to generating surrogates according to a stationary linear Gaussian process (ARMA in discrete time) that shares the same power spectrum as the data. Note that the coefficients and the order of the model need not be estimated and are not determined by the procedure.

IAAFT surrogates improve on FT surrogates by providing an exact match of the values in the surrogates and the values present in the data [78]. This corresponds to generating surrogates according to a stationary linear Gaussian process rescaled by an invertible, time-independent, non-linear measurement function:

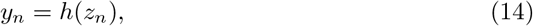

where *h* is the measurement function, *z*_*n*_ an ARMA model as described in “Linear surrogates” (Results Section), and *y*_*n*_ is constrained by the procedure to have the same PDF as the data, and approximately the same power spectrum. Due to *h*, the FT surrogate requirement that *y*_*n*_ has a Gaussian noise structure is relaxed. The IAAFT surrogate method in particular, but also a number of other linear surrogate methods, have been successfully applied in various fields. For in-depths reviews of the methods available and their applications, see [79–81].

As mentioned in the Results Section, FT and IAAFT surrogates assume that the data is stationary. This shortcoming can be addressed by using GWR surrogates instead. The GWR method is based on a maximal overlap discrete wavelet transform (MODWT) implementation of the IAAFT algorithm first described in [82] and later refined in [83]. The algorithm details are available in [30], but the basic ingredients are fixing a proportion of the MODWT wavelet coefficients, and applying the IAAFT algorithm to each scale and to the dataset as a whole. Surrogates can be computed along a continuum parametrised by *ρ*, where *ρ* = 0 corresponds to IAAFT surrogates, and *ρ* = 1 corresponds to the data. The fixed coefficients are related to *ρ* as follows.

The energy of a signal of length *N* = 2^*J*^ is proportional to

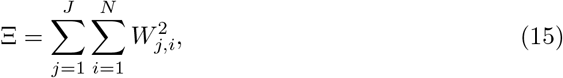

where *W*_*j,i*_ is the wavelet coefficient at scale *j* and temporal position *i*. Let us define an energy threshold Ξ_0_ = *ρ*Ξ. Going from the largest to the smallest, squared wavelet coefficients are summed irrespective of scale and position until Ξ_0_ is reached.

Coefficients that contributed to the sum are fixed for this level of *ρ*, i.e. left out of the IAAFT phase randomization. An example of surrogates for a range of *ρ* values is given in Fig 13.

**Fig 13.**
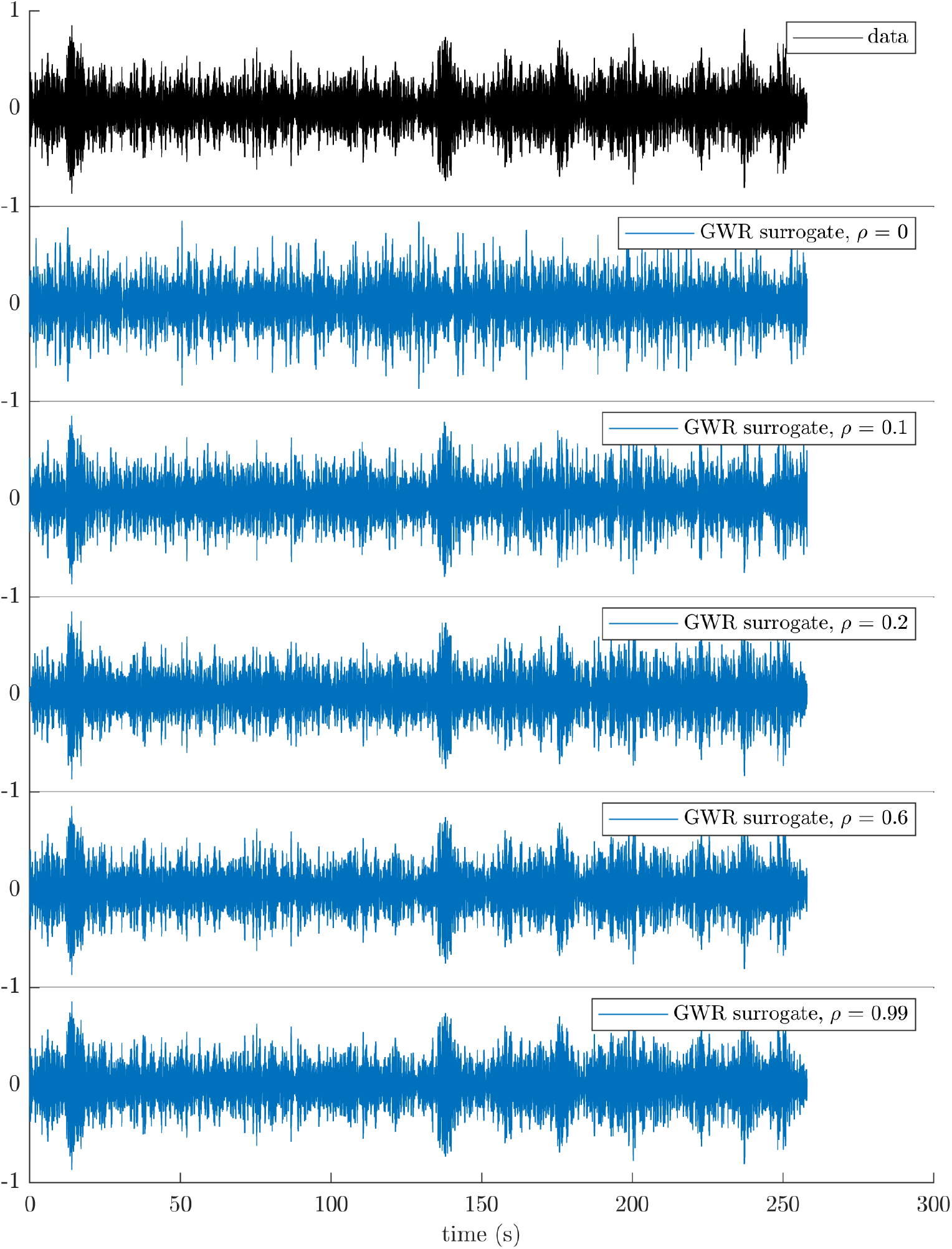
Filtered LFP for patient 6L OFF (black), and corresponding GWR surrogates for a range of *ρ* levels (blue). Most of the data temporal variability is already accounted for at *ρ* = 0.1. The plots share the same time axis.

### Fitting procedure

We describe in this subsection the fitting procedure used to fit all the models of this study to data. Differences between models will be highlighted when applicable. The general fitting procedure is similar to what was used in [84]. For each fit, random sets of model parameters are generated from uniform distributions with appropriate bounds. For envelope models, all parameters are accepted. For WC models, parameters impacting the PSD and the average burst duration profile are coupled, and to improve optimisation efficiency, parameters are accepted when the PSD peak of the corresponding model is within 1 Hz and 30% in magnitude of the data PSD peak. The first 5000 sets of parameters for WC models and 200 sets of parameters for envelope models are optimised in parallel on a supercomputer with the generalized pattern search algorithm [85, 86]. We use Matlab’s implementation of the algorithm with the “positive basis 2N” poll method. Parameters are put on a similar scale to improve search robustness. A mesh size of 10^*−*5^, and a function call budget of 600 calls are used.

At each optimisation step, the optimiser returns the cost

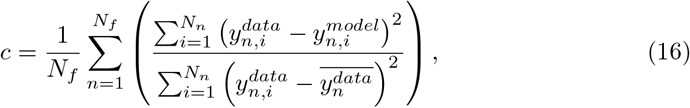

where *N*_*f*_ is the number of features, *y*_*n*_ the features considered, *N*_*n*_ the length of *y*_*n*_, and 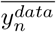 the mean of the data feature *n*. For envelope models, the PSD is not considered in the optimisation, while for WC models both the PSD and the average burst duration profiles are considered. At the end of the procedure, we reject fits with an envelope PDF very different from the data if any (specifically, when the envelope PDF *R*^2^ is smaller than zero), and the fit with the highest *R*^2^ = 1 − *c* is deemed the best fit.

Model simulations are performed with a Euler–Maruyama scheme, except for the OU model which is simulated according to the exact updating equation [87]

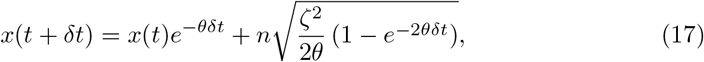

where *n* are independent samples of the standard normal distribution. In the simple OU case, the envelope is prevented from being negative by shifting it by the absolute value of its 0.1^th^ percentile, and setting values below that to zero. This shift does not affect average burst duration profiles (with thresholds expressed as percentiles), and makes OU models equivalent to the form given by equation (7), with *n*_*P*_ = 1 and *d*_0_ *>* 1, which is the form we use to report fitted OU model parameters. For higher degree models, a positive envelope is enforced by retaining only the absolute value of the next point at each integration step, which has a negligible impact on positive thresholds. The time step used in all cases was 10^*−*3^ s, roughly equivalent to the data sampling rate. At each optimisation step, the features (average burst duration for all models, plus PSD for WC models) are computed on five repeats of 1000 s and are averaged over the five repeats. The average burst duration profiles are computed from the model envelope (model output for envelope models, modulus of the analytic signal of the model output for WC models), based on the same thresholds and minimum burst duration as for data analysis (see “Extracting power spectra and bursting features”).

### Average burst duration in an envelope model

#### Continuous model MFPT

To derive the average burst duration of the time discretization of the stochastic process described by equation (8), we start by drawing on classic results pertaining to the continuous-time stochastic process itself. We consider the MFPT problem on [*L,* + ∞) with a single boundary at *x* = *L*, and a reflective boundary at *x* = +∞. The backward Fokker-Planck operator corresponding to the continuous-time stochastic process reads

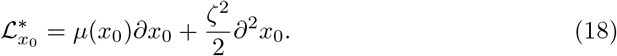

As reviewed in [35], the MFPT from *x*_0_ to *L* which we denote 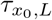 can be found as the solution of the differential equation

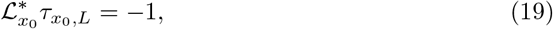

where 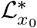 is the backward Fokker-Planck operator acting on the starting point *x*_0_.

Equation (19) is a first order differential equation in 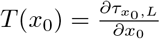 given by

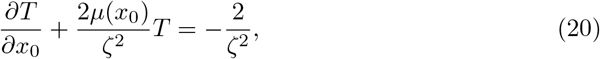

whose solution reads

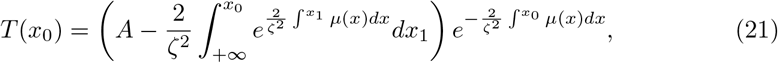

where *A* is an integration constant, which is zero in the continuous case (reflective boundary at +*∞*). We can therefore write *T* as

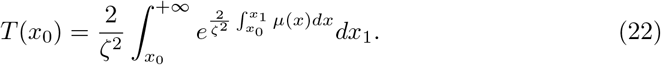

Integrating *T* (*x*_0_), we obtain the MFPT as

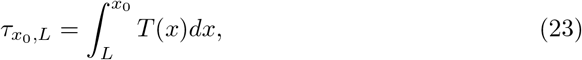

where we have used the absorbing boundary condition at *L* (*τ*_*L,L*_ = 0). This leads to the expression for the MFPT in the continuous model presented in the Results Section (equation (9)). Finally, for *x*_0_ = *L* + *δ* and *δ « L*, the continuous-time MFPT close to the boundary *τ*_*L*+*δ,L*_ can be approximated by

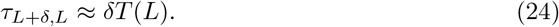

#### Discretized model

We now consider a time discretization of the stochastic process described by equation (8) with time step *dt*, and derive an expression for the average burst duration of the discretized model in three steps. We correct for the systematic bias of the continuous model MFPT and MFPT derivative in steps 1 and 2, respectively. In step 3, we relate average burst duration and MFPT. As a reminder, we use a tilde to distinguish quantities that can be readily measured in a discretized system from the continuous system quantities introduced earlier. For clarity of exposition, we also denote with a hat and double hat intermediate quantities that will be introduced as the derivation progresses. The three steps will make use of the transition probability for small *dt* = *t − t′* to first order in 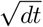,

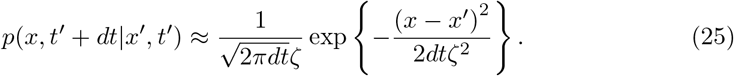

##### Step 1: MFPT bias

When close to the boundary *x* = *L*, the continuous-time model is more likely to cross the boundary earlier, which makes for a systematic under-estimation of the discrete MFPT by the continuous model. The first correction to the continuous MFPT 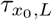 is therefore to add a constant correction 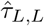 to it, which describes the average additional time taken by the discrete model once close to the boundary:

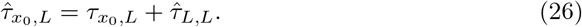

Another way of looking at it is that in the continuous model, all trajectories starting from the threshold will cross the boundary in vanishingly short times, which is reflected by the boundary condition *τ*_*L,L*_ = 0. This is not the case for the discrete model, as some trajectories starting at *L* will end up above *L* a time step later. In fact, by considering trajectories starting at *L* and the transition probability for one time step *dt*, 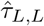 can be estimated as

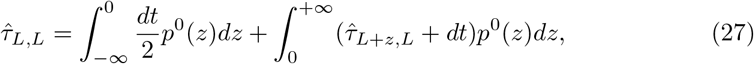

where *p*^*x*^(*z*) = *p*(*L* + *x* + *z, t′* + *dt L* + *x, t′*), and the burst duration of a trajectory starting at *L* but ending below *L* a time step later has been approximated by 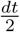. Using equation (26), we find

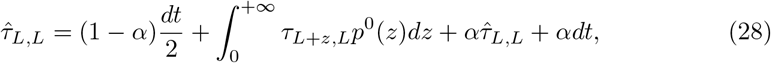

with 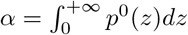. We therefore have

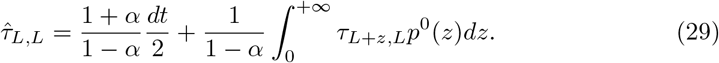

As *p*^0^ decays quickly away from *L* for small *dt*, we use equation (24), and obtain to first order in 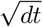

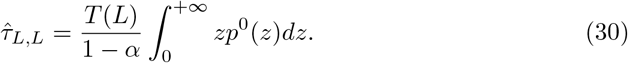

To first order 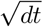, *p*^0^(*z*) = *p*(*L* + *z*, *t’* + *dt|L*, *t’*) in is an even function of *z* (see equation (25)) and 1 *− α* = *α*. Therefore,

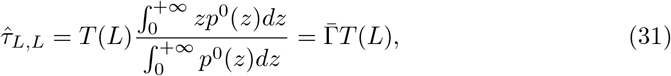

where 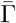 is the average step above *L* starting from *L*. To first order in 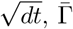 does not depend on *μ* and *L* and is given by

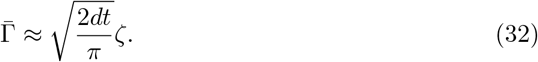

##### Step 2: MFPT derivative

As the discrete model is less likely to cross the boundary when close to it, the derivative of the MFPT at the boundary will also be affected. We model this effect with a correction to the continuous model through a non-zero *A* (integration constant in equation (21)). We define

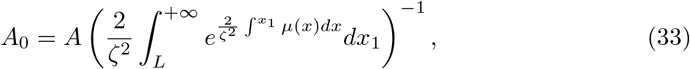

in such a way that the corrected MFPT derivative at the boundary 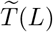 equals (*A*_0_ + 1)*T*(*L*). To show that *A* should not be zero in the discrete-time case and provide an approximation of the value of *A*_0_, we are going to approximate 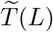 by

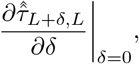

where 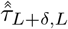 is defined next and will explicitly take into account the distribution of the first time step from *L* + *δ*, similarly to what was done in the previous paragraph. We are assuming here that most of the difference in the derivative with the continuous case comes from the first time step, and that the MFPT will subsequently evolve according to equation (26). Starting from *L* + *δ* and considering explicitly the first time step and its associated transition probability, we can write

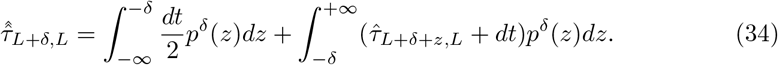

With the change of variable *x* = *δ* + *z*, we obtain to first order in 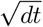

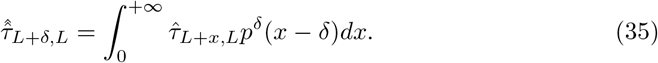

Making use of 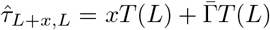 (see equations (24), (26), and (31)), we express

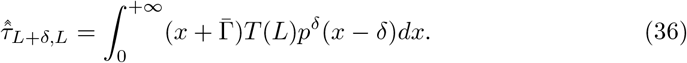

From there, we obtain to zeroth order in 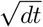 (*A*_0_ + 1 will multiply a 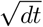 factor in the final expression)

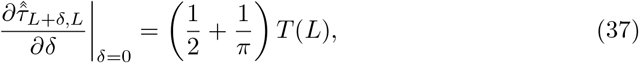

which yields 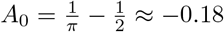. This provides a rationale for a non-zero *A*_0_ in the discretization and a good approximation of the optimal value of *A*_0_, which was found empirically to be 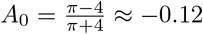 (the discrepancy may come from considering only the contribution from the first time step). We use this latter value in what follows, and for *x « L*, the MFPT in the discretized model is obtained as

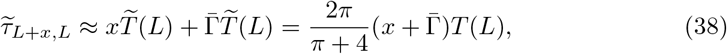

with *T*(*L*) given by equation (22).

##### Step 3: overshoot distribution

We have just obtained an approximation for the MFPT in the discretized model (equation (38)), and the last step is to derive from this result the average burst duration for the discretized model as a function of the threshold *L*. In the discretization, unless simulations are started exactly at *L* before every single burst is analysed (which is not compatible with modelling data), the distribution of boundary overshoots from below has to be taken into account (see Fig 9 in the Results Section). Averaging equation (38) over the overshoot distribution of density *χ* gives the average burst duration 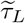 that can be measured from one long time series of the discretized model as

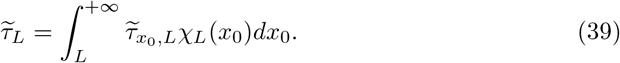

Contrary to 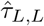, the notation 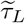 does not refer to an exact start of trajectories from *L* before each burst. We also note that the contribution of boundary overshoots from above when getting back to *L* at the end of the burst is negligible compared to stopping exactly at the threshold as it cannot change 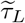 by more than a time step. This is not the case at the beginning of a burst as the further away from the threshold, the less likely a premature end of the burst is, which has a drastic influence on burst duration. As *χ*_*L*_ decays quickly away from *L* for small *dt* (the average overshoot scales with 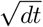 as we will see in equation (43)), equation (39) can be approximated by

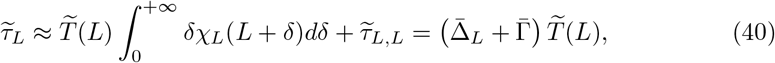

where 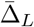 is the average overshoot at threshold *L*. An approximation for the average overshoot can be obtained by considering the probability of the state *x* given that *dt* before, the state was anywhere below *L*, which is given by

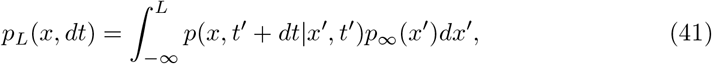

where *p*_∞_(*x*) is the stationary probability density of the process normalised by its integral over (−∞, *L*] and *p* is the transition probability to first order in 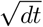 introduced earlier (equation (25)). The overshoot density is

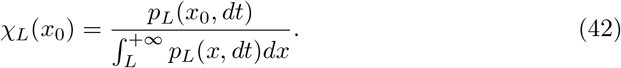

For small *dt* the stationary probability density contribution reduces to a constant that cancels out in the overshoot density. The average overshoot is obtained to first order in 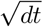 as

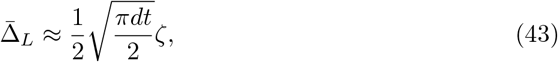

which does not depend on the drift function *μ* (to first order in 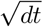) and as a result does not depend on *L* either. We will therefore drop the subscript *L* and simply denote the average burst duration by 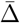.

This leads to a direct relationship between 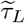 and *T*(*L*),

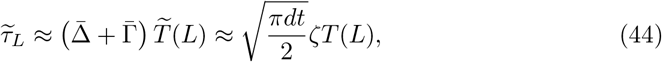

and the full expression presented in the Results Section for the average burst duration of a discretized envelope model to first order in 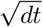 is

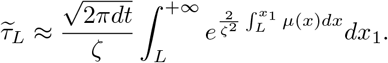

#### Application to two envelope models

**OU** We are considering here the discretization of an OU process centered on zero (equation (5)). To highlight the dependence of average burst duration on parameters, we are going to express equation (11) with *L* as a percentile of the time series values. Equation (11) is found in the Results section, and is a direct application of the previous result. The stationary probability density for our centered OU process is

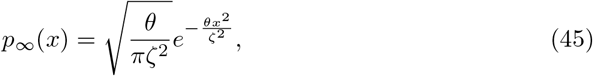

and the associated CDF Φ reads

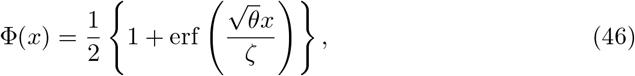

where erf is the error function. Let us denote by *L*_%_ the percentile rank (between 0 and 1) corresponding to the threshold *L*. By definition, Φ(*L*) = *L*_%_, and therefore

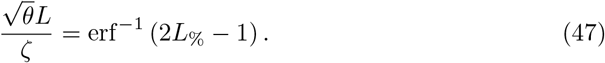

This leads to an expression for the average burst duration of a discrete OU process as a function of threshold percentile rank:

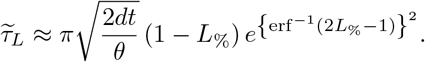

### Third degree polynomial

We are considering here the discretization of a third degree polynomial envelope model (equation (7) with *n*_*P*_ = 3). The direct application of the general expression for 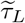 obtained previously provides an approximation for the average burst duration of this model to first order in 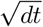, which reads

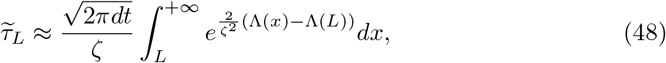

with

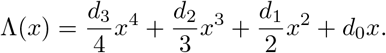

The average burst duration expression for a third degree polynomial model is not as simple as in the OU case, but it can easily be evaluated numerically.

## Passage method

### From average burst duration profile to envelope dynamics

We are concerned here with the time discretization of the continuous-time stochastic process described by equation (8), where we are assuming that the real valued drift function *μ* can depend on *x*_*t*_, and the diffusion term *ζ* is constant in time and space. Provided that the derivative of the first passage time *T* (*x*_0_) is never zero, the differential equation satisfied by *T* (equation (20)) can be re-arranged into

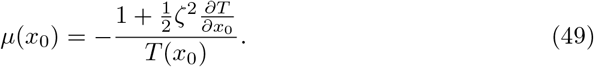

We have shown previously that in the discretized system considered, there is a simple relationship to first order in 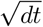 between the average burst duration 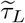 at threshold *L* and *T*(*L*) (equation (44)). Thus, to first order in 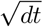,

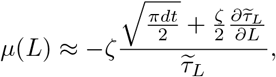

which we discuss in the Results Section.

### Data preparation

Simulation of the synthetic data used to test envelope dynamics recovery was performed with the same methods as in “Fitting procedure”. Synthetic data were generated by an OU model (same parameters as patient 3L ON minimal model, see Supplementary Table I in S3) and a fifth degree polynomial envelope model (equation (7), with *n*_*P*_ = 5, and parameters given in Supplementary Table K in S3). OU synthetic data were shifted as before, while for the fifth degree model, only the absolute value of the next point was retained at each integration step. For patient data the envelope was obtained as the modulus of the analytic signal of the filtered data as in “Extracting power spectra and bursting features”.

### Envelope dynamics recovery

Three hundred thresholds equally spaced from one fiftieth to 90% of the maximum envelope value are considered (extreme thresholds will not yield reliable measures). At each threshold, the average burst duration 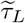 is obtained, and the resulting curve is smoothed across thresholds (LOWESS method, which stands for locally weighted scatterplot smoothing, smoothing span of the threshold range divided by eight). The derivative of 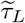 is then estimated numerically and smoothed (LOWESS, smoothing span of the threshold range divided by five). In the case of patient data, the first 10% to 20% of both 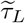 and its derivative are not smoothed as the fast decay of 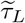 at the left edge of the profile is not handled properly by the smoothing function. Finally, the drift function *μ* is reconstructed based on equation (13). The noise parameter *ζ* is simply taken as the known value used to generate the envelope for synthetic data, and is estimated for data (24% and 27% of the time series standard deviation for OFF medication and ON medication, respectively). These values were obtained for patient data by adjusting the noise parameter so the burst duration profile and the envelope inverse CDF of the inferred dynamics best match the data. The time step *dt* is taken as the time step used for forward simulation for synthetic data, and as the time scale of variation of the envelope for patient data (roughly a beta cycle, 0.05 s). The mean of the drift function is inferred by considering all the data available, while SEM error bars are obtained by dividing all the data available in four segments and repeating the process on each segment (noise parameter fixed to the value used for the mean).

### Simulation of inferred envelope dynamics

The inferred dynamics are known at 300 equally spaced points between *x*_*min*_ and *x*_*max*_, so we define

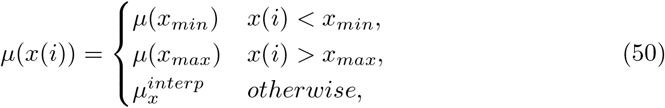

where 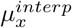 is the linear interpolation of *μ* between the closest *x* points framing *x*(*i*) at which *μ* is known. Forward simulations of the inferred envelope dynamics are then performed according to equation (8) with a Euler–Maruyama scheme (the time step is taken as *dt*). Additionally, at each integration step, only the absolute value of the next point *x*(*i* + 1) is retained to prevent the envelope from becoming negative. Five repeats of ten times the data duration are simulated. The resulting envelopes are used to compute the mean and SEM error bars of the average burst duration profile and the inverse CDF of the inferred envelope dynamics.

## Supporting information

**S1 Average burst rate.** The derivation of the relationship between average burst rate and average burst duration is presented here, as well as an expression for the average burst rate of an OU envelope model.

**S2 Supplementary Figures.** Supplementary Figures pertaining to data analysis are presented here.

**S3 Supplementary Tables.** Supplementary Tables pertaining to fits, and testing of the passage method are presented here.

## Software availability

We are sharing code at https://github.com/benoit-du/beta-burst-dyn to compute average burst duration profiles, and to infer envelope models using the passage method.

## Acknowledgements

The authors would like to thank Prof. Chris Keylock for sharing code to compute GWR surrogates. We acknowledge the use of the University of Oxford Advanced Research Computing facility in carrying out this work http://dx.doi.org/10.5281/zenodo.22558.

## Author contributions

### Conceptualization

Benoit Duchet, Filippo Ghezzi, Gihan Weerasinghe, Gerd Tinkhauser, Peter Brown, Christian Bick, Rafal Bogacz

### Formal analysis

Benoit Duchet

### Investigation

Benoit Duchet, Filippo Ghezzi

### Resources

Gerd Tinkhauser, Andrea A Kuhn, Peter Brown

### Writing – Original Draft Preparation

Benoit Duchet

### Writing – Review & Editing

Benoit Duchet, Filippo Ghezzi, Gihan Weerasinghe, Gerd Tinkhauser, Andrea A Kuhn, Peter Brown, Christian Bick, Rafal Bogacz

## Notes

### Competing Interest Statement

PB is a consultant with Medtronic.

## References

1. Brown P. Oscillatory nature of human basal ganglia activity: relationship to the pathophysiology of Parkinson’s disease. Movement Disorders. 2003;18(4):357–363. doi:10.1002/mds.10358.

2. Kuhn AA, Kupsch A, Schneider GH, Brown P. Reduction in subthalamic 8-35 Hz oscillatory activity correlates with clinical improvement in Parkinson’s disease. Eur J Neurosci. 2006;23(7):1956–1960. doi:10.1111/j.1460-9568.2006.04717.x.

3. Kühn AA, Kempf F, Brücke C, Doyle LG, Martinez-Torres I, Pogosyan A, et al. High-frequency stimulation of the subthalamic nucleus suppresses oscillatory *β* activity in patients with Parkinson’s disease in parallel with improvement in motor performance. Journal of Neuroscience. 2008;28(24):6165–6173. doi:10.1523/JNEUROSCI.0282-08.2008.

4. Kühn AA, Tsui A, Aziz T, Ray N, Brücke C, Kupsch A, et al. Pathological synchronisation in the subthalamic nucleus of patients with Parkinson’s disease relates to both bradykinesia and rigidity. Experimental Neurology. 2009;215(2):380–387. doi:10.1016/j.expneurol.2008.11.008.

5. Tan H, Pogosyan A, Anzak A, Foltynie T, Limousin P, Zrinzo L, et al. Frequency specific activity in subthalamic nucleus correlates with hand bradykinesia in Parkinson’s disease. Experimental Neurology. 2013;240(1):122–129. doi:10.1016/j.expneurol.2012.11.011.

6. Steiner LA, Neumann WJ, Staub-Bartelt F, Herz DM, Tan H, Pogosyan A, et al. Subthalamic beta dynamics mirror Parkinsonian bradykinesia months after neurostimulator implantation. Movement Disorders. 2017;32(8):1183–1190. doi:10.1002/mds.27068.

7. Brittain JS, Brown P. Oscillations and the basal ganglia: motor control and beyond. Neuroimage. 2014;85 Pt 2:637–647. doi:10.1016/j.neuroimage.2013.05.084.

8. Khawaldeh S, Tinkhauser G, Shah SA, Peterman K, Debove I, Nguyen TAK, et al. Subthalamic nucleus activity dynamics and limb movement prediction in Parkinson’s disease. Brain. 2020;143(2):582–596. doi:10.1093/brain/awz417.

9. Feingold J, Gibson DJ, Depasquale B, Graybiel AM. Bursts of beta oscillation differentiate postperformance activity in the striatum and motor cortex of monkeys performing movement tasks. Proceedings of the National Academy of Sciences of the United States of America. 2015;112(44):13687–13692. doi:10.1073/pnas.1517629112.

10. Sherman MA, Lee S, Law R, Haegens S, Thorn CA, Hämäläinen MS, et al. Neural mechanisms of transient neocortical beta rhythms: Converging evidence from humans, computational modeling, monkeys, and mice. Proceedings of the National Academy of Sciences of the United States of America. 2016;113(33):E4885–E4894. doi:10.1073/pnas.1604135113.

11. Shin H, Law R, Tsutsui S, Moore CI, Jones SR. The rate of transient beta frequency events predicts behavior across tasks and species. eLife. 2017;6. doi:10.7554/eLife.29086.

12. Tinkhauser G, Pogosyan A, Tan H, Herz DM, Kühn AA, Brown P. Beta burst dynamics in Parkinson’s disease off and on dopaminergic medication. Brain. 2017;140(11):2968–2981. doi:10.1093/brain/awx252.

13. Tinkhauser G, Pogosyan A, Little S, Beudel M, Herz DM, Tan H, et al. The modulatory effect of adaptive deep brain stimulation on beta bursts in Parkinson’s disease. Brain. 2017;140(4):1053–1067. doi:10.1093/brain/awx010.

14. Torrecillos F, Tinkhauser G, Fischer P, Green AL, Aziz TZ, Foltynie T, et al. Modulation of beta bursts in the subthalamic nucleus predicts motor performance. Journal of Neuroscience. 2018;38(41):8905–8917. doi:10.1523/JNEUROSCI.1314-18.2018.

15. Lofredi R, Tan H, Neumann WJ, Yeh CH, Schneider GH, Kühn AA, et al. Beta bursts during continuous movements accompany the velocity decrement in Parkinson’s disease patients. Neurobiology of disease. 2019;doi:10.1016/j.nbd.2019.03.013.

16. Ahn S, Zauber SE, Worth RM, Witt T, Rubchinsky LL. Neural synchronization: Average strength vs. temporal patterning. Clinical Neurophysiology. 2018;129(4):842–844. doi:10.1016/j.clinph.2018.01.063.

17. Anidi CM, O’Day JJ, Anderson RW, Afzal MF, Syrkin-Nikolau J, Velisar A, et al. Neuromodulation targets pathological not physiological beta bursts during gait in Parkinson’s disease. Neurobiology of disease. 2018;120:107–117. doi:10.1016/j.nbd.2018.09.004.

18. Deffains M, Iskhakova L, Katabi S, Israel Z, Bergman H. Longer *β* oscillatory episodes reliably identify pathological subthalamic activity in Parkinsonism. Movement Disorders. 2018;33(10):1609–1618. doi:10.1002/mds.27418.

19. Schroll H, Hamker FH. Basal Ganglia dysfunctions in movement disorders: What can be learned from computational simulations. Movement Disorders. 2016;31(11):1591–1601. doi:10.1002/mds.26719.

20. Bahuguna J, Sahasranamam A, Kumar A. Uncoupling the roles of firing rates and spike bursts in shaping the STN-GPe beta band oscillations. PLoS Computational Biology. 2020;16(3):e1007748. doi:10.1371/journal.pcbi.1007748.

21. Ahn S, Zauber SE, Worth RM, Rubchinsky LL. Synchronized beta-band oscillations in a model of the globus pallidus-subthalamic nucleus network under external input. Frontiers in computational neuroscience. 2016;10:134. doi:10.3389/fncom.2016.00134.

22. Mirzaei A, Kumar A, Leventhal D, Mallet N, Aertsen A, Berke J, et al. Sensorimotor processing in the basal ganglia leads to transient beta oscillations during behavior. Journal of Neuroscience. 2017;37(46):11220–11232. doi:10.1523/JNEUROSCI.1289-17.2017.

23. Wilson HR, Cowan JD. Excitatory and inhibitory interactions in localized populations of model neurons. Biophysical journal. 1972;12(1):1–24. doi:10.1016/S0006-3495(72)86068-5.

24. Powanwe AS, Longtin A. Determinants of Brain Rhythm Burst Statistics. Scientific Reports. 2019;9(1):1–28. doi:10.1038/s41598-019-54444-z.

25. Brown P, Oliviero A, Mazzone P, Insola A, Tonali P, Di Lazzaro V. Dopamine dependency of oscillations between subthalamic nucleus and pallidum in Parkinson’s disease. Journal of Neuroscience. 2001;21(3):1033–1038. doi:10.1523/jneurosci.21-03-01033.2001.

26. Kühn AA, Tsui A, Aziz T, Ray N, Brücke C, Kupsch A, et al. Pathological synchronisation in the subthalamic nucleus of patients with Parkinson’s disease relates to both bradykinesia and rigidity. Experimental Neurology. 2009;215(2):380–387. doi:10.1016/j.expneurol.2008.11.008.

27. Hohlefeld FU, Huebl J, Huchzermeyer C, Schneider GH, Schönecker T, Kühn AA, et al. Long-range temporal correlations in the subthalamic nucleus of patients with Parkinson’s disease. European Journal of Neuroscience. 2012;36(6):2812–2821. doi:10.1111/j.1460-9568.2012.08198.x.

28. Neumann WJ, Staub-Bartelt F, Horn A, Schanda J, Schneider GH, Brown P, et al. Long term correlation of subthalamic beta band activity with motor impairment in patients with Parkinson’s disease. Clinical Neurophysiology. 2017;128(11):2286–2291. doi:10.1016/j.clinph.2017.08.028.

29. Theiler J, Eubank S, Longtin A, Galdrikian B, Farmer JD. Testing for nonlinearity in time series: the method of surrogate data. Physica D: Nonlinear Phenomena. 1992;58(1-4):77–94. doi:10.1016/0167-2789(92)90102-S.

30. Keylock CJ. Characterizing the structure of nonlinear systems using gradual wavelet reconstruction. Nonlinear Processes in Geophysics. 2010;17(6):615–632. doi:10.5194/npg-17-615-2010.

31. Holgado AJ, Terry JR, Bogacz R. Conditions for the generation of beta oscillations in the subthalamic nucleus-globus pallidus network. J Neurosci. 2010;30(37):12340–12352. doi:10.1523/JNEUROSCI.0817-10.2010.

32. Pavlides A, Hogan SJ, Bogacz R. Improved conditions for the generation of beta oscillations in the subthalamic nucleus–globus pallidus network. Eur J Neurosci. 2012;36(2):2229–2239. doi:10.1111/j.1460-9568.2012.08105.x.

33. Pavlides A, Hogan SJ, Bogacz R. Computational Models Describing Possible Mechanisms for Generation of Excessive Beta Oscillations in Parkinson’s Disease. PLoS Comput Biol. 2015;11(12):e1004609. doi:10.1371/journal.pcbi.1004609.

34. Barnard GA. A new test for 2 x 2 tables. Nature. 1945;156(3954):177. doi:10.1038/156177a0.

35. Grebenkov DS. First exit times of harmonically trapped particles: a didactic review. Journal of Physics A: Mathematical and Theoretical. 2014;48(1):13001. doi:10.1088/1751-8113/48/1/013001.

36. Heideman SG, Quinn AJ, Woolrich MW, van Ede F, Nobre AC. Dissecting beta-state changes during timed movement preparation in Parkinson’s disease. Progress in Neurobiology. 2019; p. 101731. doi:10.1016/j.pneurobio.2019.101731.

37. Quinn AJ, van Ede F, Brookes MJ, Heideman SG, Nowak M, Seedat ZA, et al. Unpacking Transient Event Dynamics in Electrophysiological Power Spectra. Brain topography. 2019;32:1020–1034. doi:10.1007/s10548-019-00745-5.

38. Schmidt SL, Peters JJ, Turner DA, Grill WM. Continuous deep brain stimulation of the subthalamic nucleus may not modulate beta bursts in patients with Parkinson’s disease. Brain Stimulation. 2019;13(2):433–443. doi:10.1016/j.brs.2019.12.008.

39. Park C, Worth RM, Rubchinsky LL. Fine temporal structure of beta oscillations synchronization in subthalamic nucleus in Parkinson’s disease. Journal of Neurophysiology. 2010;103(5):2707–2716. doi:10.1152/jn.00724.2009.

40. Tinkhauser G, Torrecillos F, Duclos Y, Tan H, Pogosyan A, Fischer P, et al. Beta burst coupling across the motor circuit in Parkinson’s disease. Neurobiology of disease. 2018;117:217–225. doi:10.1016/j.nbd.2018.06.007.

41. Stam CJ. Nonlinear dynamical analysis of EEG and MEG: Review of an emerging field. Clinical Neurophysiology. 2005;116(10):2266–2301. doi:10.1016/j.clinph.2005.06.011.

42. Marceglia S, Foffani G, Bianchi AM, Baselli G, Tamma F, Egidi M, et al. Dopamine-dependent non-linear correlation between subthalamic rhythms in Parkinson’s disease. Journal of Physiology. 2006;571(3):579–591. doi:10.1113/jphysiol.2005.100271.

43. Jackson N, Cole SR, Voytek B, Swann NC. Characteristics of Waveform Shape in Parkinson’s Disease Detected with Scalp Electroencephalography. eNeuro. 2019; p. ENEURO. 0151–19.2019. doi:10.1523/ENEURO.0151-19.2019.

44. Cole SR, van der Meij R, Peterson EJ, de Hemptinne C, Starr PA, Voytek B. Nonsinusoidal beta oscillations reflect cortical pathophysiology in parkinson’s disease. Journal of Neuroscience. 2017;37(18):4830–4840. doi:10.1523/JNEUROSCI.2208-16.2017.

45. Lim J, Sanghera MK, Darbin O, Stewart RM, Jankovic J, Simpson R. Nonlinear temporal organization of neuronal discharge in the basal ganglia of Parkinson’s disease patients. Experimental Neurology. 2010;224(2):542–544. doi:10.1016/j.expneurol.2010.05.021.

46. Camara C, Subramaniyam NP, Warwick K, Parkkonen L, Aziz T, Pereda E. Non-Linear Dynamical Analysis of Resting Tremor for Demand-Driven Deep Brain Stimulation. Sensors. 2019;19(11):2507. doi:10.3390/s19112507.

47. Özkurt TE, Akram H, Zrinzo L, Limousin P, Foltynie T, Oswal A, et al. Identification of nonlinear features in cortical and subcortical signals of Parkinson’s Disease patients via a novel efficient measure. NeuroImage. 2020; p. 117356. doi:10.1016/j.neuroimage.2020.117356.

48. Rosa M, Arlotti M, Marceglia S, Cogiamanian F, Ardolino G, Fonzo AD, et al.. Adaptive deep brain stimulation controls levodopa-induced side effects in Parkinsonian patients; 2017.

49. Arlotti M, Marceglia S, Foffani G, Volkmann J, Lozano AM, Moro E, et al. Eight-hours adaptive deep brain stimulation in patients with Parkinson disease. Neurology. 2018;90(11):e971–e976. doi:10.1212/WNL.0000000000005121.

50. Moraud EM, Tinkhauser G, Agrawal M, Brown P, Bogacz R. Predicting beta bursts from local field potentials to improve closed-loop DBS paradigms in Parkinson’s patients. In: 2018 40th Annual International Conference of the IEEE Engineering in Medicine and Biology Society (EMBC). IEEE; 2018. p. 3766–3796.

51. Cagnan H, Mallet N, Moll CKE, Gulberti A, Holt AB, Westphal M, et al. Temporal evolution of beta bursts in the parkinsonian cortical and basal ganglia network. Proceedings of the National Academy of Sciences. 2019;116(32):16095–16104. doi:10.1073/pnas.1819975116.

52. Gillies A, Willshaw D, Li Z. Subthalamic-pallidal interactions are critical in determining normal and abnormal functioning of the basal ganglia. Proc Biol Sci. 2002;269(1491):545–551. doi:10.1098/rspb.2001.1817.

53. Terman D, Rubin JE, Yew AC, Wilson CJ. Activity Patterns in a Model for the Subthalamopallidal Network of the Basal Ganglia. Journal of Neuroscience. 2002;22(7):2963–2976. doi:10.1523/jneurosci.22-07-02963.2002.

54. Tachibana Y, Iwamuro H, Kita H, Takada M, Nambu A. Subthalamo-pallidal interactions underlying parkinsonian neuronal oscillations in the primate basal ganglia. European Journal of Neuroscience. 2011;34(9):1470–1484. doi:10.1111/j.1460-9568.2011.07865.x.

55. Nevado-Holgado AJ, Mallet N, Magill PJ, Bogacz R. Effective connectivity of the subthalamic nucleus-globus pallidus network during Parkinsonian oscillations. Journal of Physiology. 2014;592(7):1429–1455. doi:10.1113/jphysiol.2013.259721.

56. Leblois A, Boraud T, Meissner W, Bergman H, Hansel D. Competition between feedback loops underlies normal and pathological dynamics in the basal ganglia. Journal of Neuroscience. 2006;26(13):3567–3583. doi:10.1523/JNEUROSCI.5050-05.2006.

57. McCarthy MM, Moore-Kochlacs C, Gu X, Boyden ES, Han X, Kopell N. Striatal origin of the pathologic beta oscillations in Parkinson’s disease. Proceedings of the National Academy of Sciences of the United States of America. 2011;108(28):11620–11625. doi:10.1073/pnas.1107748108.

58. Broadie M, Glasserman P, Steven K. A continuity correction for discrete barrier options. Mathematical Finance. 1997;7(4):325–349. doi:10.1111/1467-9965.00035.

59. Taillefumier T, Magnasco MO. A Fast Algorithm for the First-Passage Times of Gauss-Markov Processes with Hölder Continuous Boundaries. Journal of Statistical Physics. 2010;140(6):1130–1156. doi:10.1007/s10955-010-0033-6.

60. Drugowitsch J. Fast and accurate Monte Carlo sampling of first-passage times from Wiener diffusion models. Scientific Reports. 2016;6. doi:10.1038/srep20490.

61. Herrmann S, Zucca C. Exact Simulation of the First-Passage Time of Diffusions. Journal of Scientific Computing. 2019;79(3):1477–1504. doi:10.1007/s10915-018-00900-3.

62. Collin-Dufresne P, Goldstein RS. Do credit spreads reflect stationary leverage ratios? Journal of Finance. 2001;56(5):1929–1957. doi:10.1111/0022-1082.00395.

63. Larralde H. A first passage time distribution for a discrete version of the Ornstein-Uhlenbeck process. Journal of Physics A: Mathematical and General. 2004;37(12):3759–3767. doi:10.1088/0305-4470/37/12/003.

64. Ditlevsen S, Lansky P. Comparison of statistical methods for estimation of the input parameters in the Ornstein-Uhlenbeck neuronal model from first-passage times data. In: AIP Conference Proceedings. vol. 1028; 2008. p. 171–185.

65. Lundqvist M, Rose J, Herman P, Brincat SLL, Buschman TJJ, Miller EKK. Gamma and Beta Bursts Underlie Working Memory. Neuron. 2016;90(1):152–164. doi:10.1016/j.neuron.2016.02.028.

66. Brady B, Power L, Bardouille T. Age-Related Trends in Neuromagnetic Transient Beta Burst Characteristics During a Sensorimotor Task and Rest in the Cam-CAN Open-Access Dataset. NeuroImage. 2020;222:117245. doi:10.1016/j.neuroimage.2020.117245.

67. Smelyanskiy VN, Luchinsky DG, Stefanovska A, McClintock PVE. Inference of a Nonlinear Stochastic Model of the Cardiorespiratory Interaction. Physical review letters. 2005;94(9):98101. doi:10.1103/PhysRevLett.94.098101.

68. Smelyanskiy VN, Luchinsky DG, Timucin DA, Bandrivskyy A, Timuçin DA, Bandrivskyy A. Reconstruction of stochastic nonlinear dynamical models from trajectory measurements. Physical Review E. 2005;72(2):26202. doi:10.1103/PhysRevE.72.026202.

69. Stankovski T, Duggento A, McClintock PVE, Stefanovska A. Inference of time-evolving coupled dynamical systems in the presence of noise. Physical review letters. 2012;109(2):24101. doi:10.1103/PhysRevLett.109.024101.

70. Callaham JL, Loiseau JC, Rigas G, Brunton SL. Nonlinear stochastic modeling with Langevin regression. arXiv e-prints. 2020; p. arXiv:2009.01006.

71. Wilting J, Priesemann V. Inferring collective dynamical states from widely unobserved systems. Nature Communications. 2018;9(1). doi:10.1038/s41467-018-04725-4.

72. Fernández-Ruiz A, Oliva A, de Oliveira EF, Rocha-Almeida F, Tingley D, Buzsáki G. Long-duration hippocampal sharp wave ripples improve memory. Science. 2019;364(6445):1082–1086. doi:10.1126/science.aax0758.

73. Sporn S, Hein T, Ruiz MH. Alterations in the amplitude and burst rate of beta oscillations impair reward-dependent motor learning in anxiety. eLife. 2020;9:1–40. doi:10.7554/eLife.50654.

74. Luo H, Huang Y, Xiao X, Dai W, Nie Y, Geng X, et al. Functional dynamics of thalamic local field potentials correlate with modulation of neuropathic pain. European Journal of Neuroscience. 2020;51(2):628–640. doi:10.1111/ejn.14569.

75. Benjamini Y, Hochberg Y. Controlling the false discovery rate: a practical and powerful approach to multiple testing. Journal of the royal statistical society Series B (Methodological). 1995; p. 289–300.

76. Storey JD, Taylor JE, Siegmund D. Strong control, conservative point estimation and simultaneous conservative consistency of false discovery rates: a unified approach. Journal of the Royal Statistical Society: Series B (Statistical Methodology). 2004;66(1):187–205. doi:10.1111/j.1467-9868.2004.00439.x.

77. Benjamini Y, Krieger AM, Yekutieli D. Adaptive linear step-up procedures that control the false discovery rate. Biometrika. 2006;93(3):491–507. doi:10.1093/biomet/93.3.491.

78. Schreiber T, Schmitz A. Improved surrogate data for nonlinearity tests. Physical Review Letters. 1996;77(4):635–638. doi:10.1103/PhysRevLett.77.635.

79. Schreiber T, Schmitz A. Surrogate time series. Physica D: Nonlinear Phenomena. 2000;142(3-4):346–382. doi:10.1016/S0167-2789(00)00043-9.

80. Lancaster G, Iatsenko D, Pidde A, Ticcinelli V, Stefanovska A. Surrogate data for hypothesis testing of physical systems. Physics Reports. 2018;doi:10.1016/j.physrep.2018.06.001.

81. Keylock CJ. Hypothesis Testing for Nonlinear Phenomena in the Geosciences Using Synthetic, Surrogate Data. Earth and Space Science. 2019;6(1):41–58. doi:10.1029/2018EA000435.

82. Keylock CJ. Constrained surrogate time series with preservation of the mean and variance structure. Physical Review E - Statistical, Nonlinear, and Soft Matter Physics. 2006;73(3). doi:10.1103/PhysRevE.73.036707.

83. Keylock CJ. A wavelet-based method for surrogate data generation. Physica D: Nonlinear Phenomena. 2007;225(2):219–228. doi:10.1016/j.physd.2006.10.012.

84. Duchet B, Weerasinghe G, Cagnan H, Brown P, Bick C, Bogacz R. Phase-dependence of response curves to deep brain stimulation and their relationship: from essential tremor patient data to a Wilson–Cowan model. The Journal of Mathematical Neuroscience. 2020;10(1):4. doi:10.1186/s13408-020-00081-0.

85. Torczon V. On the convergence of pattern search algorithms. SIAM Journal on Optimization. 1997;7(1):1–25. doi:10.1137/S1052623493250780.

86. Audet C, Dennis JE. Analysis of generalized pattern searches. SIAM Journal on Optimization. 2003;13(3):889–903. doi:10.1137/S1052623400378742.

87. Gillespie DT. Exact numerical simulation of the Ornstein-Uhlenbeck process and its integral. Physical Review E. 1996;54(2):2084. doi:10.1103/PhysRevE.54.2084.

