## Supplementary material S1 for "Average beta burst duration profiles provide a signature of dynamical changes between the ON and OFF medication states in Parkinson’s disease"

### S1 – Relationship between average burst rate and average burst duration

The average burst rate $r_L$ at threshold $L$ can be directly obtained from the average burst duration at threshold $L$ (denoted $\widetilde{\tau}_L$). The average burst duration at threshold $L$ in a recording of length $T$ is given by

$$\widetilde{\tau}_L = \frac{T_L^{\text{burst}}}{n_L^{\text{burst}}}, \tag{S.1}$$

where $T_L^{\text{burst}}$ is the time spent by the envelope above $L$, and $n_L^{\text{burst}}$ the number of bursts above $L$ in the recording. The average burst rate at threshold $L$ can be obtained as

$$r_L = \frac{n_L^{\text{burst}}}{T} = \frac{1}{\widetilde{\tau}_L}\frac{T_L^{\text{burst}}}{T}. \tag{S.2}$$

The ratio $\frac{T_L^{\text{burst}}}{T}$ is the probability that the envelope is above $L$. The probability that the envelope is below $L$ is simply the cumulative distribution function of the envelope amplitude evaluated at $L$, which we denote $\Phi(L)$. We therefore have

$$r_L = \frac{1}{\widetilde{\tau}_L}\{1-\Phi(L)\} = \frac{1}{\widetilde{\tau}_L}(1-L_{\%}), \tag{S.3}$$

where $L_{\%}$ is the percentile rank. This relationship holds in patient data as shown in Fig S.1.

It follows that for an envelope described by an OU process as introduced in the main text, the average burst rate is

$$r_L \approx \frac{1}{\pi}\sqrt{\frac{\theta}{2dt}}e^{-\left\{\text{erf}^{-1}(2L_{\%}-1)\right\}^2}, \tag{S.4}$$

where we have used Equation (13) from the main text. As shown in Fig S.2, this result is very close to simulations of OU envelope models.

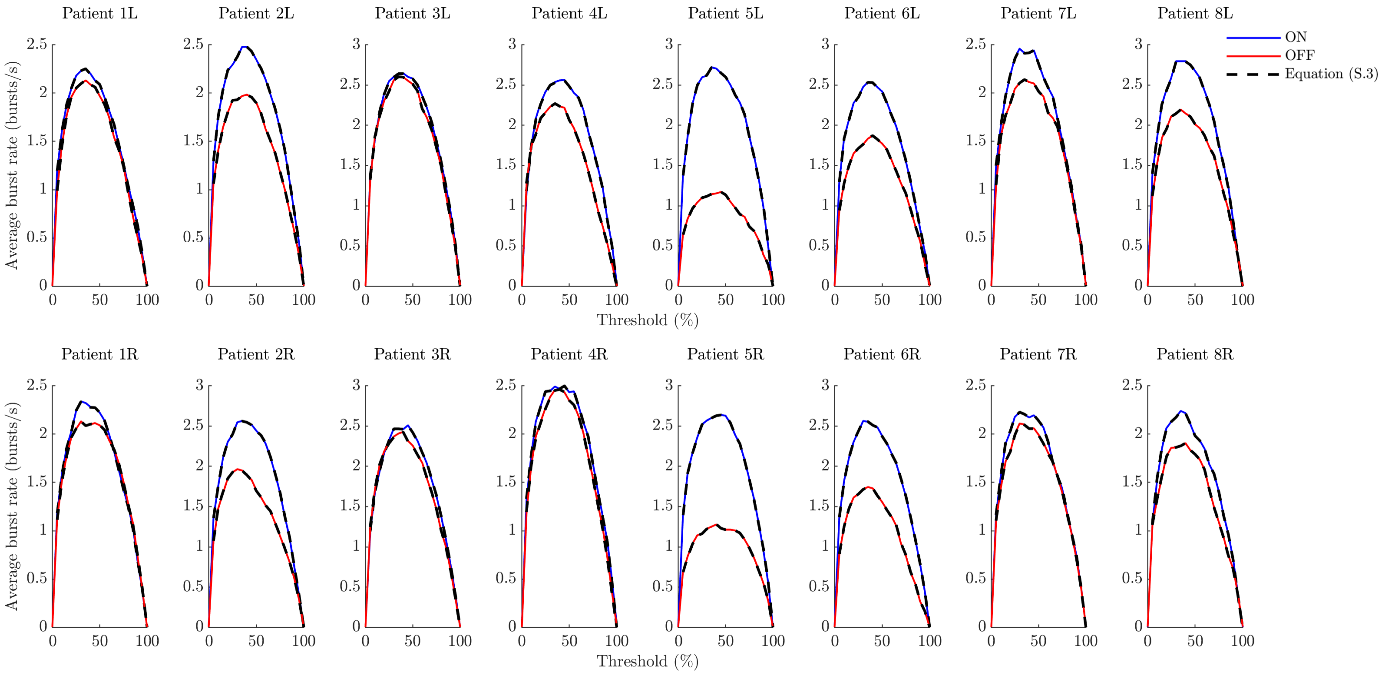


**Fig S.1. Testing the relationship between average burst rate and average burst duration in patient data.** Top panels correspond to left hemispheres, and bottom panels to right hemispheres. The average burst rate is obtained from data and is plotted as blue lines (ON medication) and red lines (OFF medication). The right hand side of Equation (S.3) is obtained using data average burst duration profiles, and is plotted as black dashed lines both ON and OFF medication. In this analysis, no minimum burst duration is enforced when calculating average burst duration and average burst rate. Since bursts are on average shorter ON medication, there are more of them and the average burst rate is larger.

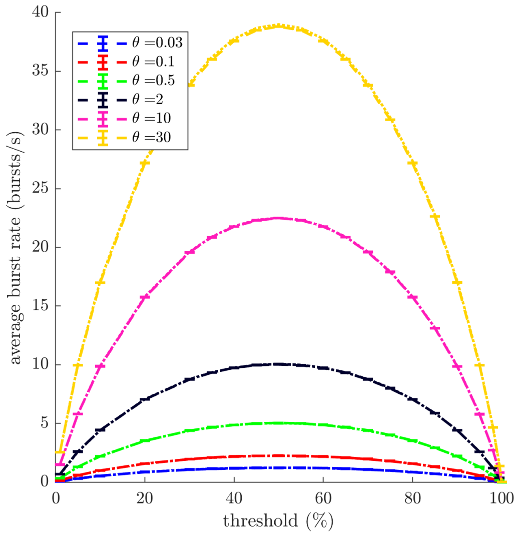


**Fig S.2. Test of the average burst rate expression for an OU envelope model.** Average burst rate profiles from simulations of OU processes are compared to equation (S.4) for a range of decay parameters and $\zeta = 1$. Simulations consist of five repeats of $10^5$ s, with a time step of 1 ms (simulations use the exact updating equation described in the main text).
