## Supplementary material S2 for "Average beta burst duration profiles provide a signature of dynamical changes between the ON and OFF medication states in Parkinson’s disease"

### S2 – Supplementary figures

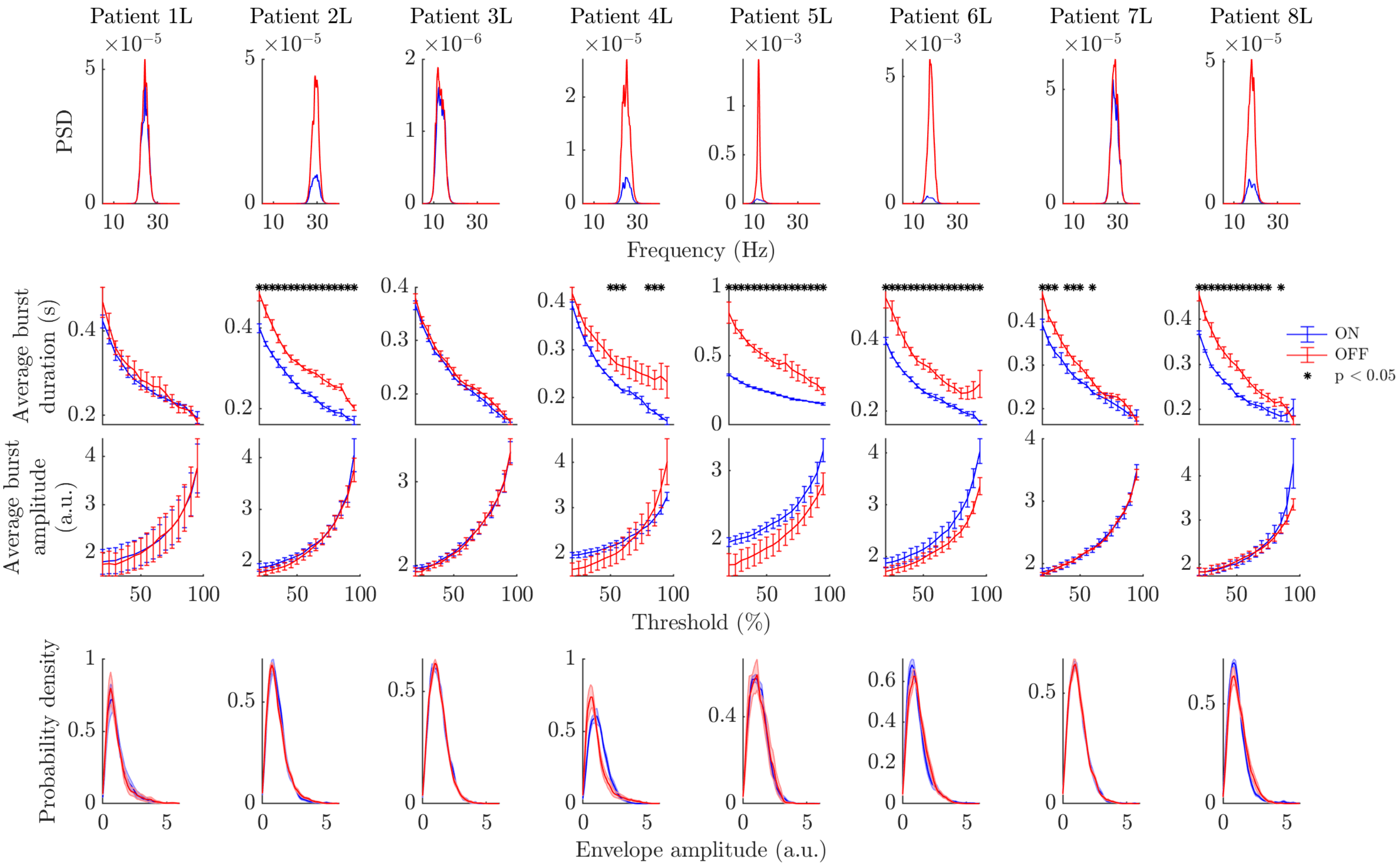


**Fig A. Power spectra and bursting features ON and OFF Levodopa (left hemispheres).** Each column corresponds to the left hemisphere of one of the eight patients. Each row corresponds to a feature, the ON state is in blue, and the OFF state in red. The first row shows power spectra, the second row average burst duration profiles, the third row average burst amplitude profiles, and last row envelope amplitude PDFs. Statistically significant differences under FDR control are indicated by black stars (three bursting features only). Error bars represent the SEM.

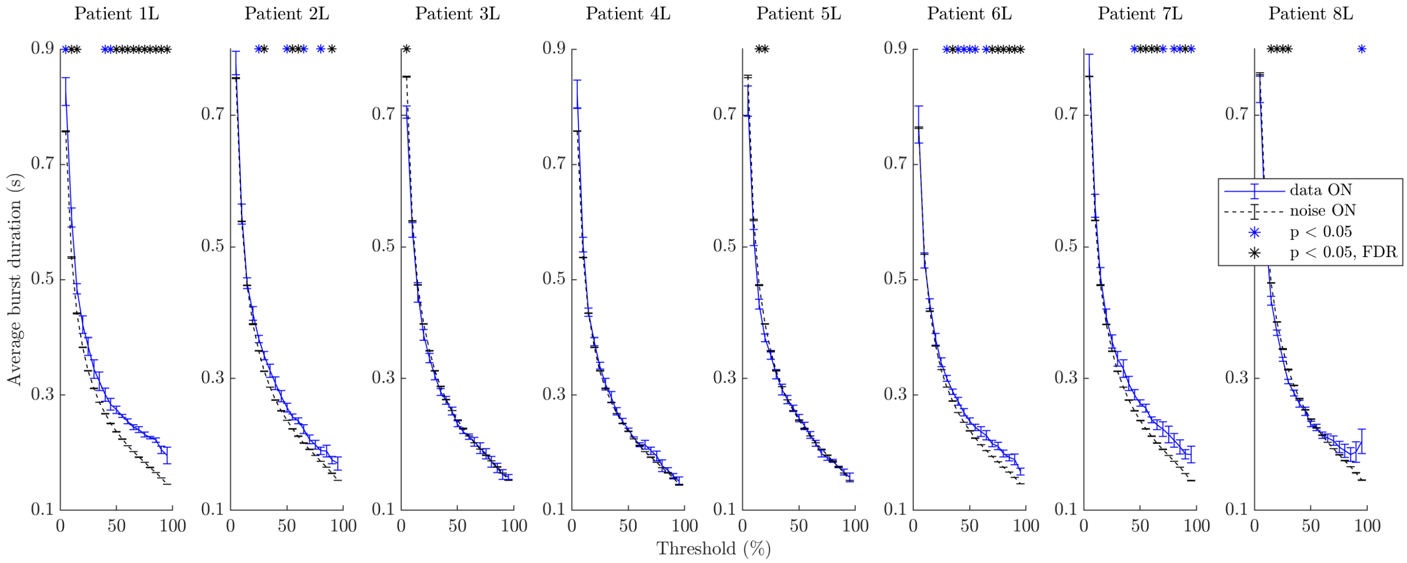


**Fig B. Average burst duration ON Levodopa compared to average burst duration of filtered pink noise (left hemispheres).** Each column corresponds to the left hemisphere of one of the eight patients. The ON state is shown by blue lines, filtered white noise by dashed black lines. Statistically significant differences under FDR control are indicated by black stars. Cases not significant under FDR control where $p < 0.05$ are indicated by blue stars. Error bars represent the SEM. Error bars are very small in the filtered pink noise case and therefore appear as single horizontal bars.

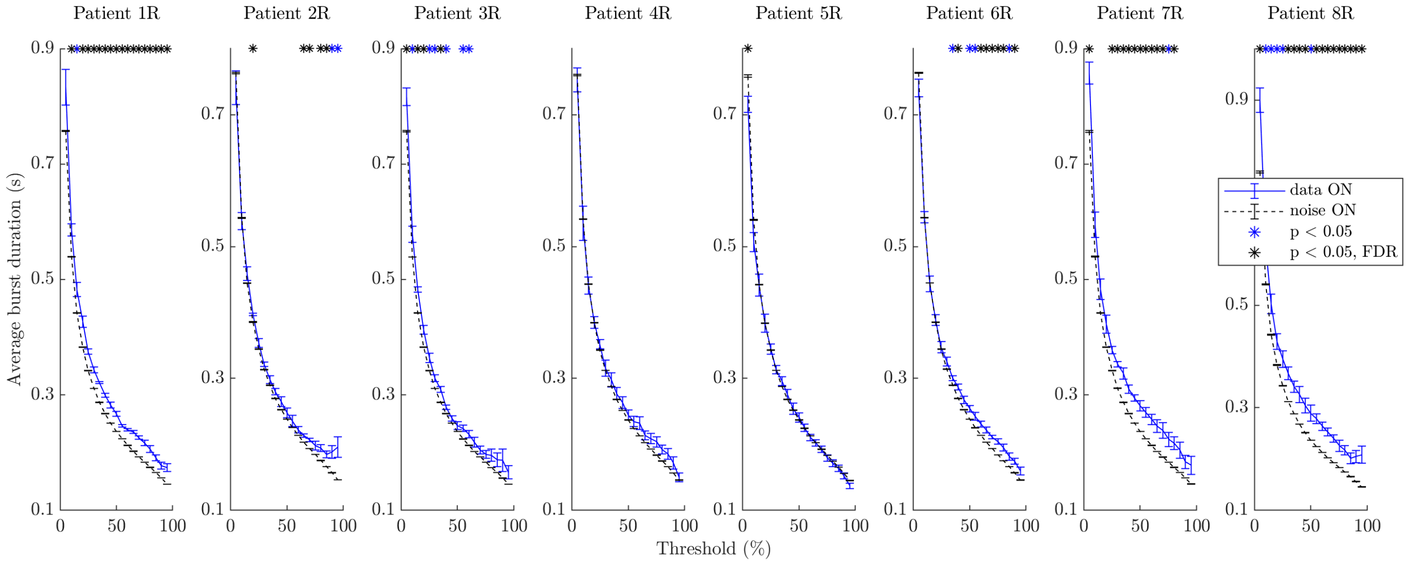


**Fig C. Average burst duration ON Levodopa compared to average burst duration of filtered pink noise (right hemispheres).** Each column corresponds to the right hemisphere of one of the eight patients. The ON state is shown by blue lines, filtered white noise by dashed black lines. Statistically significant differences under FDR control are indicated by black stars. Cases not significant under FDR control where $p < 0.05$ are indicated by blue stars. Error bars represent the SEM. Error bars are very small in the filtered pink noise case and therefore appear as single horizontal bars.

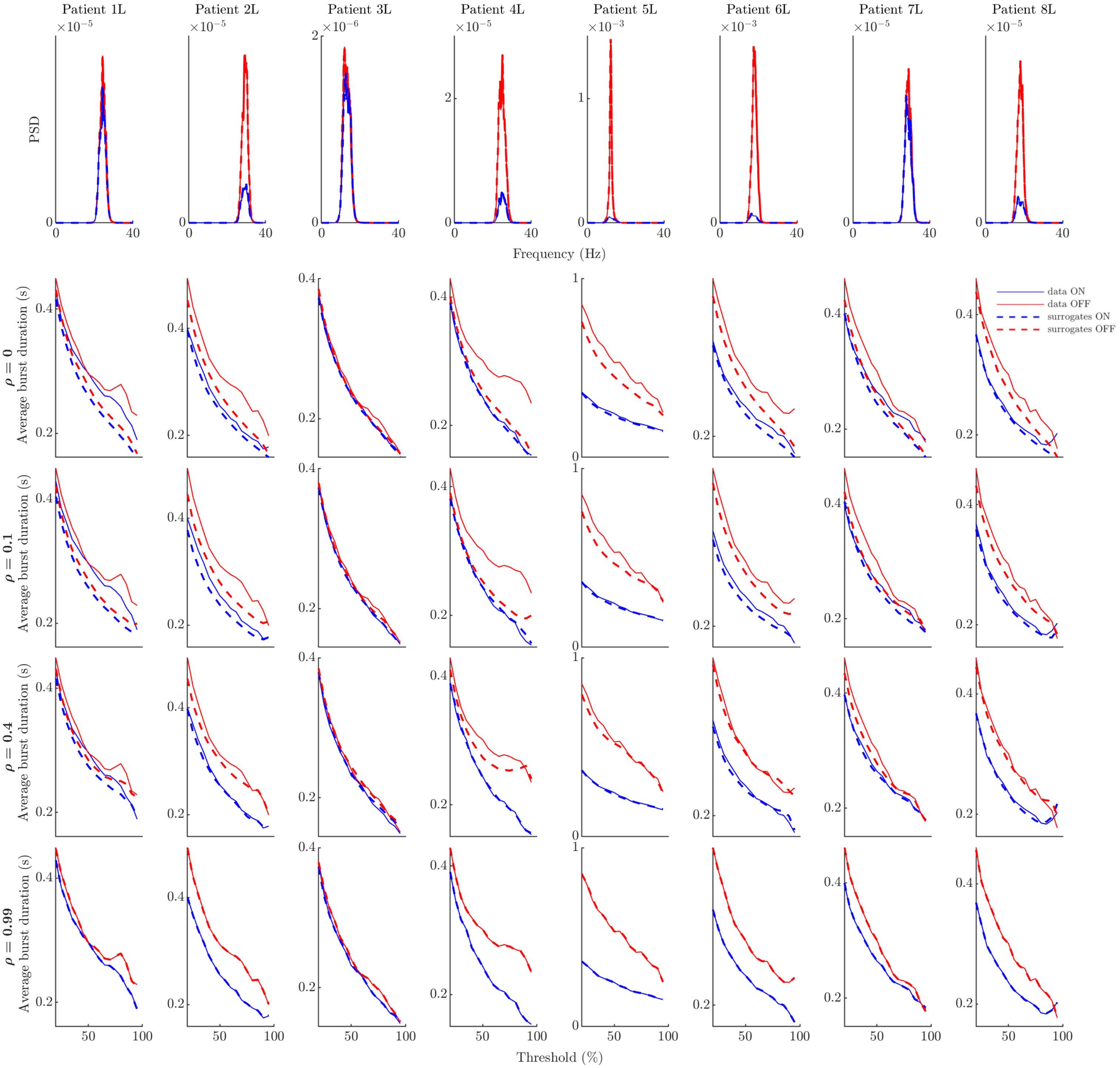


**Fig D. Power spectra and average burst duration profiles ON and OFF medication for data and GWR surrogates for a range of $\rho$ values (left hemispheres).** Each column corresponds to the left hemisphere of one of the eight patients. In all panels, data profiles are solid lines, while surrogate profiles are dashed lines. The OFF medication state is indicated in red, and the ON state in blue. Power spectra are shown in the first row (data and surrogates overlap as expected). Average burst duration profiles for various $\rho$ values are provided in the following rows. Surrogate profiles match data profiles for $\rho$ close to one as expected.

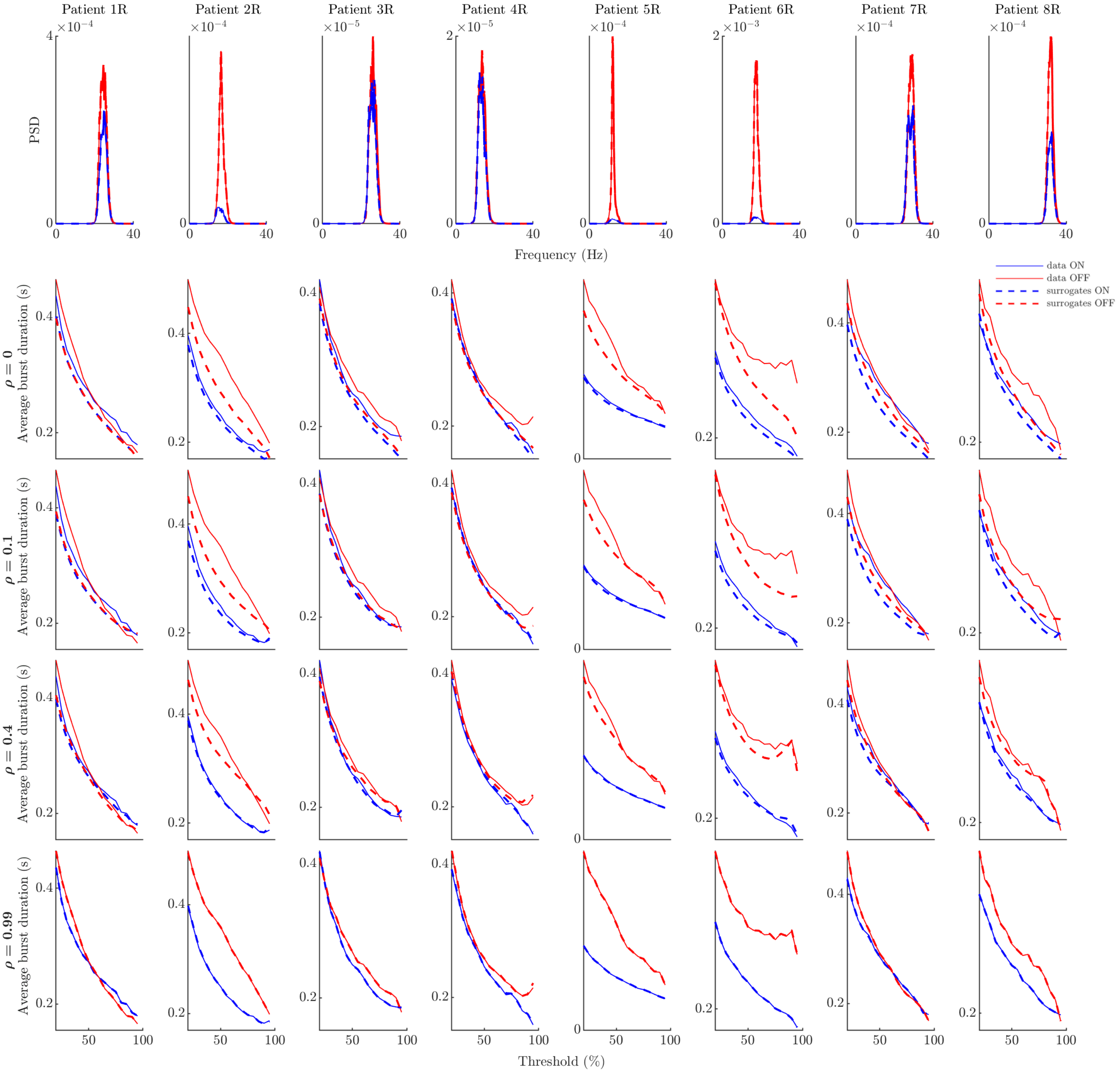


**Fig E. Power spectra and average burst duration profiles ON and OFF medication for data and GWR surrogates for a range of $\rho$ values (right hemispheres).** Each column corresponds to the right hemisphere of one of the eight patients. In all panels, data profiles are solid lines, while surrogate profiles are dashed lines. The OFF medication state is indicated in red, and the ON state in blue. Power spectra are shown in the first row (data and surrogates overlap as expected). Average burst duration profiles for various $\rho$ values are provided in the following rows. Surrogate profiles match data profiles for $\rho$ close to one as expected.

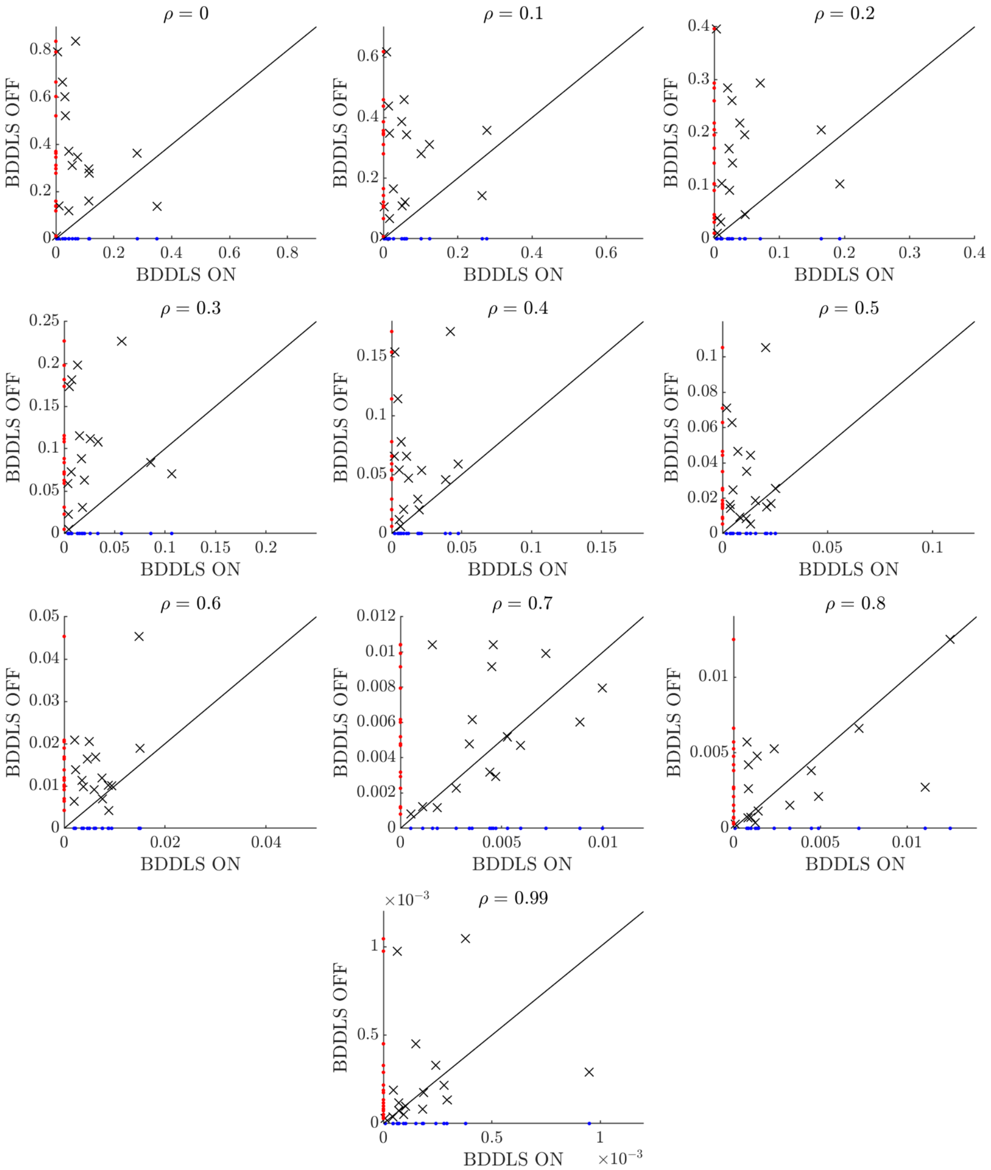


**Fig F. Comparing BDDLS between medication states.** Showing BDDLS for all 16 hemispheres OFF medication and ON medication, for all $\rho$ levels analysed. Each cross corresponds to one hemisphere. Blue and red dots are projections on the horizontal and vertical axes, and represent the ON and OFF medication BDDLS values, respectively. The black line corresponds to BDDLS OFF = BDDLS ON. Therefore, BDDLS is greater OFF medication than ON medication for hemispheres that are above this line.

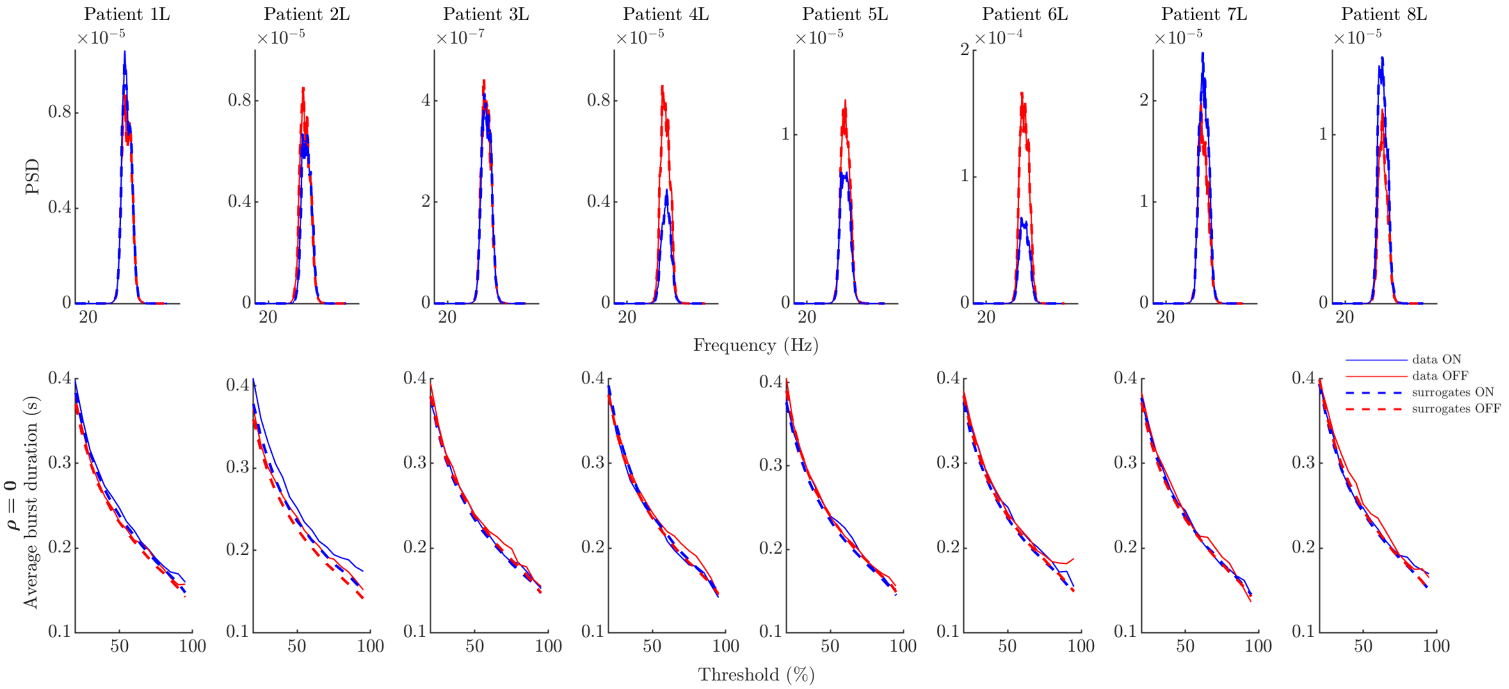


**Fig G. 35 Hz controls (left hemispheres): power spectra and average burst duration profiles ON and OFF medication.** Each column corresponds to the left hemisphere of one of the eight patients. In all panels, data profiles are solid lines, while surrogate profiles are dashed lines. The OFF medication state is indicated in red, and the ON state in blue. Power spectra are shown in the first row (data and surrogates overlap as expected). Average burst duration profiles for $\rho = 0$ are presented in the second row.

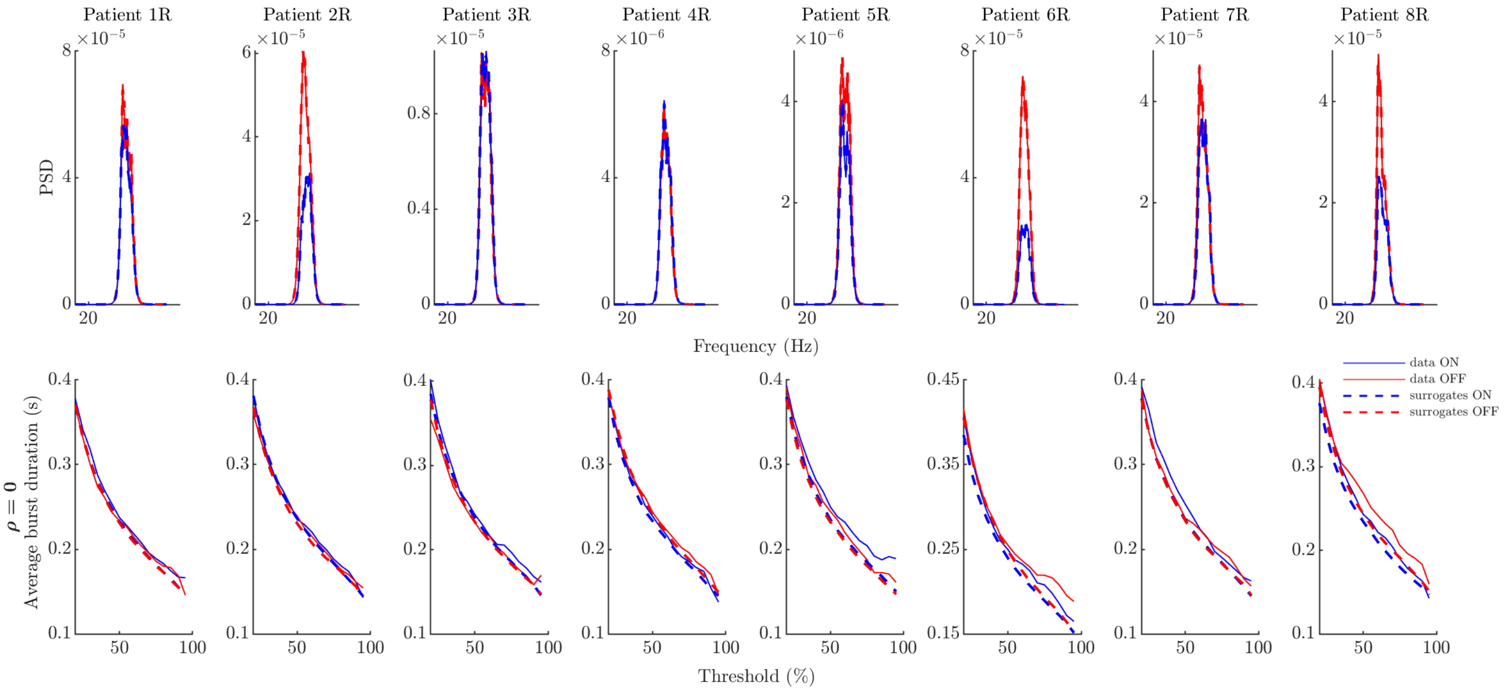


**Fig H. 35 Hz controls (right hemispheres): power spectra and average burst duration profiles ON and OFF medication.** Each column corresponds to the right hemisphere of one of the eight patients. In all panels, data profiles are solid lines, while surrogate profiles are dashed lines. The OFF medication state is indicated in red, and the ON state in blue. Power spectra are shown in the first row (data and surrogates overlap as expected). Average burst duration profiles for $\rho = 0$ are presented in the second row.

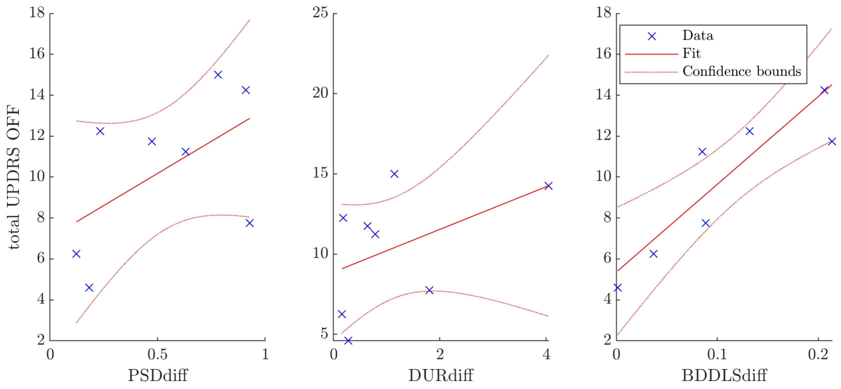


**Fig I. Total UPDRS score OFF medication as a function of the three predictors.** Spearman's correlations of 0.500, p = 0.216 (PSDdiff), 0.476, p = 0.243 (DURdiff), and 0.810, p = 0.0218* (BDDLSdiff), where * denotes significance under FDR correction. BDDLSdiff is taken at $\rho = 0.2$. Confidence bounds are at 95%.

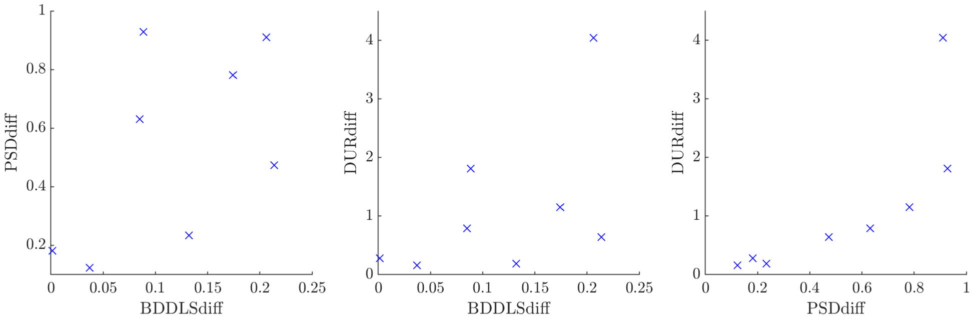


**Fig J. Correlations between the three predictors.** Spearman's correlations are 0.500, p = 0.216 between BDDLSdiff for $\rho = 0.2$ and PSDdiff (left panel); 0.476, p = 0.243 between BDDLSdiff for $\rho = 0.2$ and DURdiff (middle panel); and 0.952, p = 0.001* between DURdiff and PSDdiff (right panel), where * denotes significance under FDR correction.

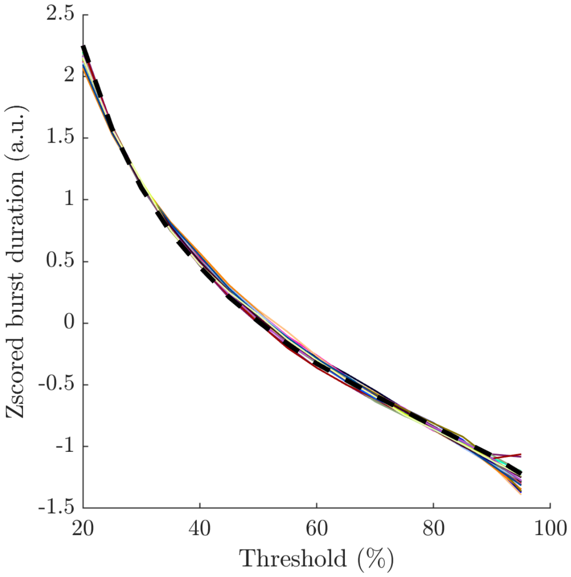


**Fig K. Z-scored average burst duration profiles of data linear surrogates ($\rho = 0$).** Shown for all patients and hemispheres available. The 16 average burst duration profiles (colored lines) closely match, highlighting that average burst duration profiles of linear systems are the same shape (potentially scaled and shifted).The black dashed line is the z-scored average burst duration profile of an OU process, indicating that the average burst duration profile of linear systems can be represented by an OU process.

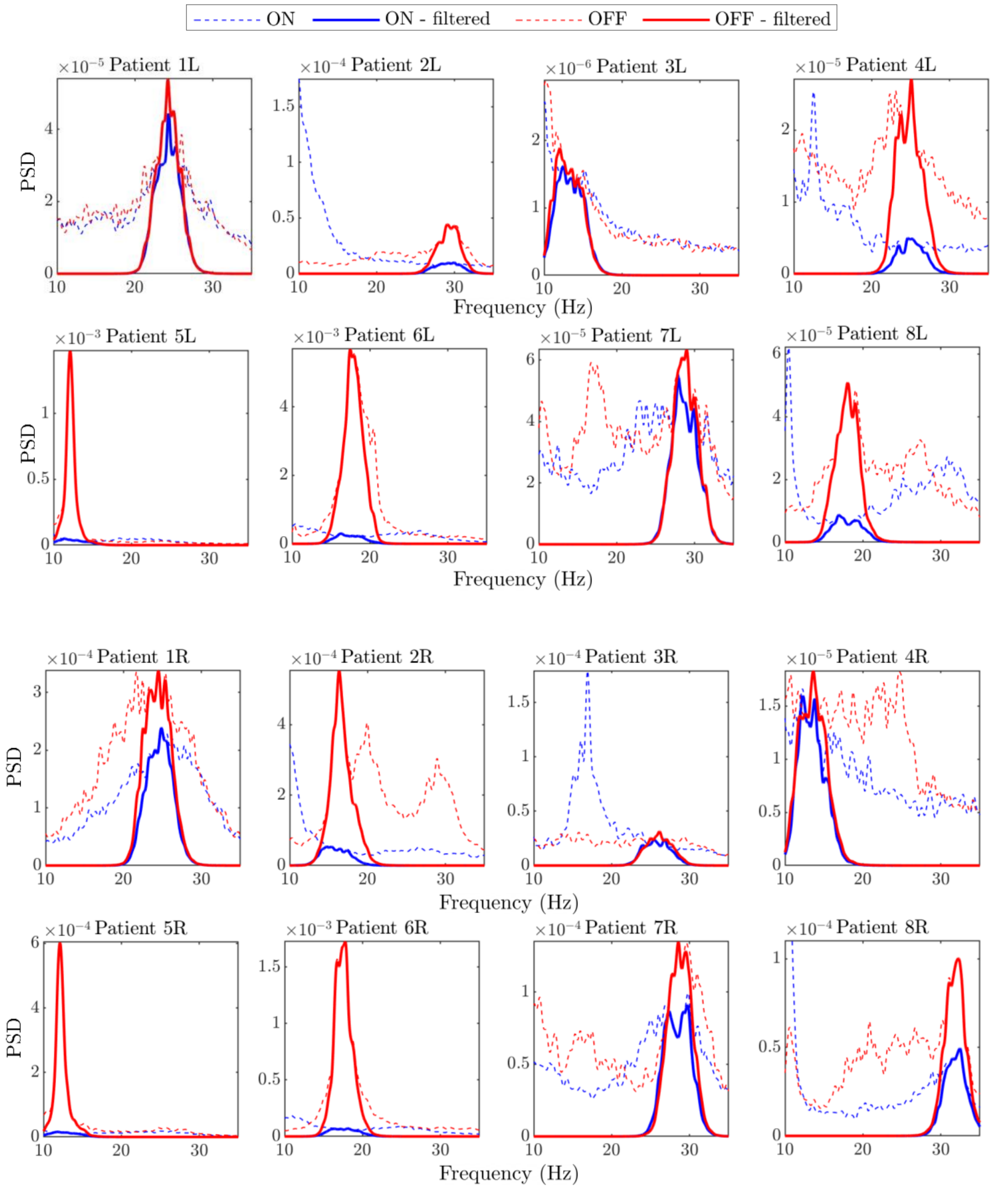


**Fig L. Power spectra of filtered and unfiltered data.** Showing the PSDs of the centered, unfiltered data (dashed lines), and of the filtered data (solid lines). All hemispheres are shown ON and OFF medication. The peak seen at 12.6 Hz ON medication for patient 4L is a harmonic of a peak at 4.2 Hz, and the peak seen at 16.6 Hz ON medication for patient 3R is a harmonic of a peak at 8.3 Hz.
