## Supplementary material S3 for "Average beta burst duration profiles provide a signature of dynamical changes between the ON and OFF medication states in Parkinson’s disease"

### S3 – Supplementary tables

| Parameter | Symbol | Best fit values | | | |
|---|---|---|---|---|---|
| | | Patient 6RON | Patient 4LON | Patient 6ROFF | Patient 4LOFF |
| I to E weight | $w_{IE}$ | 9.764 | 30.340 | 21.349 | 13.131 |
| E to I weight | $w_{EI}$ | 12.698 | 39.455 | 26.810 | 10.462 |
| I to I weight | $w_{II}$ | 0.318 | 0.570 | 0.0986 | 0.0050 |
| activation function slope | $\beta$ | 5.185 | 1.360 | 2.439 | 3.600 |
| E time constant (s) | $\Omega_E$ | 0.637 | 0.382 | 0.556 | 0.244 |
| I time constant (s) | $\Omega_I$ | 0.405 | 0.243 | 0.519 | 0.295 |
| Constant input to E | $\lambda_E$ | 0 | 0 | 0 | 0 |
| Constant input to I | $\lambda_I$ | 0 | 0 | 0 | 0 |
| Noise standard deviation | $\zeta$ | 0.0612 | 0.0161 | 0.131 | 0.0202 |

**Table A. Best parameters for fits of the linear WC model to patients 6RON, 4LON, 6ROFF, and 4LOFF.**

| Parameter | Symbol | Best fit values | |
|---|---|---|---|
| | | Patient 6ROFF | Patient 4LOFF |
| I to E weight | $w_{IE}$ | 1.215 | 20.465 |
| E to I weight | $w_{EI}$ | 2.118 | 13.364 |
| I to I weight | $w_{II}$ | 1.144 | 14.558 |
| Scaling parameter | $\eta$ | 4.100 | 0.481 |
| Steepness parameter | $\beta$ | 4.777 | 4.464 |
| E time constant (s) | $\Omega_E$ | 0.0170 | 0.0099 |
| I time constant (s) | $\Omega_I$ | 0.165 | 0.165 |
| Constant input to E | $\lambda_E$ | 2.662 | 3.578 |
| Constant input to I | $\lambda_I$ | 2.304 | 0.106 |
| Delay from I to E (s) | $\Delta_{IE}$ | 0.0039 | 0.0011 |
| Delay from E to I (s) | $\Delta_{EI}$ | 0.0005 | 0.0018 |
| Delay from I to I (s) | $\Delta_{II}$ | 0.0445 | 0.0306 |
| Noise standard deviation | $\zeta$ | 0.0710 | 0.0063 |

**Table B. Best parameters for fits of the non-linear WC model with delays to patients 6ROFF, and 4LOFF.**

| Patient | | OU | degree 2 | degree 3 | degree 4 | degree 5 |
|---|---|---|---|---|---|---|
| 1 | left | 51.30% | 77.63% | 68.99% | 92.55% | 95.70% |
| | right | 99.36% | | | | |
| 2 | left | 86.20% | 92.76% | 98.80% | | |
| | right | 82.90% | 98.82% | | | |
| 3 | left | 98.11% | | | | |
| | right | 89.15% | 94.44% | 88.79% | 97.61% | |
| 4 | left | -4.51% | 71.24% | 96.61% | | |
| | right | 88.45% | 84.91% | 92.17% | 99.33% | |
| 5 | left | 91.85% | 98.21% | | | |
| | right | 97.34% | | | | |
| 6 | left | 84.54% | 88.21% | 80.85% | 99.22% | |
| | right | -116.47% | 50.58% | 97.76% | | |
| 7 | left | 95.69% | | | | |
| | right | 98.70% | | | | |
| 8 | left | 92.85% | 96.35% | | | |
| | right | 84.29% | 94.85% | 97.76% | | |

**Table C. Average burst duration $\mathbf{R^2}$ in envelope model fits, OFF medication.** Showing both hemispheres of all patients. Cells highlighted in green correspond to $R^2 > 95\%$.

| Patient | | OU | degree 2 | degree 3 | degree 4 | degree 5 |
|---|---|---|---|---|---|---|
| 1 | left | 74.98% | 93.61% | 98.61% | | |
| | right | 95.98% | | | | |
| 2 | left | 96.96% | | | | |
| | right | 95.61% | | | | |
| 3 | left | 99.55% | | | | |
| | right | 97.19% | | | | |
| 4 | left | 99.05% | | | | |
| | right | 97.65% | | | | |
| 5 | left | 98.29% | | | | |
| | right | 99.41% | | | | |
| 6 | left | 92.80% | 96.13% | | | |
| | right | 95.43% | | | | |
| 7 | left | 88.73% | 93.69% | 99.60% | | |
| | right | 92.41% | 97.81% | | | |
| 8 | left | 86.87% | 85.71% | 96.59% | | |
| | right | 89.48% | 96.48% | | | |

**Table D. Average burst duration $\mathbf{R^2}$ in envelope model fits, ON medication.** Showing both hemispheres of all patients. Cells highlighted in green correspond to $R^2 > 95\%$.

| Patient | | OU | degree 2 | degree 3 | degree 4 | degree 5 |
|---|---|---|---|---|---|---|
| 1 | left | -96.1 | -103.0 | -95.0 | -117.8 | -121.1 |
| | right | -152.7 | | | | |
| 2 | left | -108.0 | -112.7 | -138.8 | | |
| | right | -101.5 | -138.7 | | | |
| 3 | left | -146.2 | | | | |
| | right | -118.2 | -123.3 | -109.4 | -134.1 | |
| 4 | left | -90.5 | -105.6 | -137.0 | | |
| | right | -117.5 | -107.7 | -115.4 | -154.7 | |
| 5 | left | -92.0 | -110.7 | | | |
| | right | -107.6 | | | | |
| 6 | left | -103.7 | -102.4 | -91.9 | -143.1 | |
| | right | -73.3 | -91.4 | -138.1 | | |
| 7 | left | -125.6 | | | | |
| | right | -142.3 | | | | |
| 8 | left | -119.3 | -124.5 | | | |
| | right | -102.8 | -115.0 | -125.6 | | |

**Table E. Average burst duration BIC in envelope model fits, OFF medication.** Showing both hemispheres of all patients. Models with the lowest BIC for a given patient and hemisphere are highlighted in green.

| Patient | | OU | degree 2 | degree 3 | degree 4 | degree 5 |
|---|---|---|---|---|---|---|
| 1 | left | -105.3 | -120.5 | -143.3 | | |
| | right | -130.7 | | | | |
| 2 | left | -136.5 | | | | |
| | right | -132.9 | | | | |
| 3 | left | -168.9 | | | | |
| | right | -137.5 | | | | |
| 4 | left | -155.5 | | | | |
| | right | -141.9 | | | | |
| 5 | left | -147.8 | | | | |
| | right | -162.0 | | | | |
| 6 | left | -125.0 | -129.3 | | | |
| | right | -132.2 | | | | |
| 7 | left | -119.0 | -122.7 | -164.2 | | |
| | right | -120.5 | -134.8 | | | |
| 8 | left | -120.6 | -113.7 | -133.8 | | |
| | right | -115.5 | -127.4 | | | |

**Table F. Average burst duration BIC in envelope model fits, ON medication.** Showing both hemispheres of all patients. Models with the lowest BIC for a given patient and hemisphere are highlighted in green.

| Parameter | Symbol | Best fit values | | | | | | | |
|---|---|---|---|---|---|---|---|---|---|
| | | 1LOFF | 2LOFF | 3LOFF | 4LOFF | 5LOFF | 6LOFF | 7LOFF | 8LOFF |
| Coefficient of $x^5$ | $d_5$ | - 10478516.144 | | | | | | | |
| Coefficient of $x^4$ | $d_4$ | 1604261.443 | | | | | - 23.211 | | |
| Coefficient of $x^3$ | $d_3$ | - 80454.031 | - 120.331 | | - 5224.078 | | 50.299 | | |
| Coefficient of $x^2$ | $d_2$ | 1395.819 | 51.834 | | 574.056 | - 4.988 | - 31.205 | | - 55.630 |
| Coefficient of $x^1$ | $d_1$ | - 4.397 | - 5.951 | - 6.9154 | - 20.923 | 0.118 | 4.156 | - 4.748 | 0.326 |
| Coefficient of 1 | $d_0$ | - 0.0552 | 0.0957 | 0.0378 | 0.204 | - 0.00133 | 0 | 0.140 | - 0.00203 |
| Noise standard deviation | $\zeta$ | 0.0514 | 0.0343 | 0.00671 | 0.0264 | 0.0985 | 0.384 | 0.0293 | 0.0498 |

**Table G. Best parameters for minimal envelope model fits to left hemispheres, OFF medication.**

| Parameter | Symbol | Best fit values | | | | | | | |
|---|---|---|---|---|---|---|---|---|---|
| | | 1ROFF | 2ROFF | 3ROFF | 4ROFF | 5ROFF | 6ROFF | 7ROFF | 8ROFF |
| Coefficient of $x^5$ | $d_5$ | | | | | | | | |
| Coefficient of $x^4$ | $d_4$ | | | - 9557.629 | - 160402.774 | | | | |
| Coefficient of $x^3$ | $d_3$ | | | 2812.027 | 23046.149 | | - 102.404 | | - 42.415 |
| Coefficient of $x^2$ | $d_2$ | | - 13.699 | - 218.749 | - 957.661 | | 69.085 | | 7.218 |
| Coefficient of $x^1$ | $d_1$ | - 5.200 | 0.567 | 2.078 | 9.195 | - 1.942 | -15.329 | - 4.956 | - 2.710 |
| Coefficient of 1 | $d_0$ | 0.400 | - 0.0433 | 0 | 0 | 0.103 | 0.964 | 0.203 | 0.0502 |
| Noise standard deviation | $\zeta$ | 0.0784 | 0.137 | 0.0453 | 0.0319 | 0.0356 | 0.119 | 0.0429 | 0.0583 |

**Table H. Best parameters for minimal envelope model fits to right hemispheres, OFF medication.**

| Parameter | Symbol | Best fit values | | | | | | | |
|---|---|---|---|---|---|---|---|---|---|
| | | 1LON | 2LON | 3LON | 4LON | 5LON | 6LON | 7LON | 8LON |
| Coefficient of $x^5$ | $d_5$ | | | | | | | | |
| Coefficient of $x^4$ | $d_4$ | | | | | | | | |
| Coefficient of $x^3$ | $d_3$ | - 55.261 | | | | | | - 37.816 | - 6855.253 |
| Coefficient of $x^2$ | $d_2$ | 32.229 | | | | | - 26.939 | 31.788 | 704.566 |
| Coefficient of $x^1$ | $d_1$ | - 4.980 | - 6.198 | - 7.353 | - 6.825 | - 7.302 | 0.191 | - 6.679 | - 21.287 |
| Coefficient of 1 | $d_0$ | 0.0688 | 0.0774 | 0.0390 | 0.0623 | 0.190 | 0.0183 | 0.127 | 0.157 |
| Noise standard deviation | $\zeta$ | 0.0447 | 0.0143 | 0.00631 | 0.0106 | 0.0333 | 0.147 | 0.0539 | 0.0133 |

**Table I. Best parameters for minimal envelope model fits to left hemispheres, ON medication.**

| Parameter | Symbol | Best fit values | | | | | | | |
|---|---|---|---|---|---|---|---|---|---|
| | | 1RON | 2RON | 3RON | 4RON | 5RON | 6RON | 7RON | 8RON |
| Coefficient of $x^5$ | $d_5$ | | | | | | | | |
| Coefficient of $x^4$ | $d_4$ | | | | | | | | |
| Coefficient of $x^3$ | $d_3$ | | | | | | | | |
| Coefficient of $x^2$ | $d_2$ | | | | | | | - 40.062 | - 61.285 |
| Coefficient of $x^1$ | $d_1$ | - 5.635 | - 6.491 | - 6.155 | - 6.581 | - 7.098 | - 6.400 | 0.277 | 0.574 |
| Coefficient of 1 | $d_0$ | 0.310 | 0.187 | 0.122 | 0.0995 | 0.113 | 0.219 | 0.000306 | - 0.00593 |
| Noise standard deviation | $\zeta$ | 0.0618 | 0.0332 | 0.0221 | 0.0181 | 0.0193 | 0.0398 | 0.0784 | 0.0515 |

**Table J. Best parameters for minimal envelope model fits to right hemispheres, ON medication.**

| Parameter | Symbol | Value |
|---|---|---|
| Coefficient of $x^5$ | $d_5$ | -12.67 |
| Coefficient of $x^4$ | $d_4$ | 49.73 |
| Coefficient of $x^3$ | $d_3$ | -64.47 |
| Coefficient of $x^2$ | $d_2$ | 30.12 |
| Coefficient of $x^1$ | $d_1$ | -3.81 |
| Coefficient of 1 | $d_0$ | 0 |
| Noise standard deviation | $\zeta$ | 0.828 |

**Table K. Parameters of the fifth degree polynomial drift used to generate synthetic data to test the passage method.**
